# 2’-*O*-methylation of the second transcribed nucleotide within mRNA 5’ cap impacts protein production level in a cell specific manner and contributes to RNA immune evasion

**DOI:** 10.1101/2022.02.03.478939

**Authors:** Karolina Drazkowska, Natalia Baran, Marcin Warminski, Rafal Tomecki, Anaïs Depaix, Dominik Cysewski, Renata Kasprzyk, Joanna Kowalska, Jacek Jemielity, Pawel J. Sikorski

## Abstract

In higher eukaryotes, m7G-adjacent nucleotides undergo extensive modifications. Ribose of the first or first and second transcribed nucleotides can be subjected to 2’-*O*-methylation to form cap1 or cap2, respectively. Additionally, when the first transcribed nucleotide is adenosine, it can not only undergo 2’-*O*-methylation but can also be methylated at position N6 forming N6,2’-*O*-dimethyladenosine (m6Am). Recent studies have shed some light on the functions of cap1, showing that cap1 in mammalian cells plays a crucial role in distinguishing between ‘self’ and ‘non-self’ RNA during viral infection. Here, we attempted to understand the impact of other cap methylations on RNA-related processes. Therefore, we synthesized tetranucleotide cap analogs and used them for efficient co-transcriptional RNA capping during in vitro transcription. Using this tool, we found that 2’-*O*-methylation of the second transcribed nucleotide within the mRNA 5’ cap influences protein production levels in a cell-specific manner. The presence of this modification can strongly hamper protein biosynthesis or do not influence protein production levels. Interestingly, 2’-*O*-methylation of the second transcribed nucleotide as well as the presence of N6,2’-*O*-dimethyladenosine as the first transcribed nucleotide serve as determinants that define transcripts as ‘self’ and contribute to transcript escape from the host innate immune response. Additionally, cap methylation status does not influence transcript affinity towards translation initiation factor 4E or *in vitro* susceptibility to decapping by DCP2; however what we observe is resistance of RNA capped with cap2 to DXO-mediated decapping and degradation.

**Significance Statement:** Methylation of mRNA cap structure regulates protein biosynthesis in a cell-dependent manner. Among the three known m7G cap modifications, the 2’-*O*-methylation is dominant. 2’-*O*-methylation of the first transcribed nucleotide can boost protein production, whereas the same modification of the second transcribed nucleotide can strongly decrease translation. Interestingly, we show that in the JAWS II cell line, 2’-*O*-methylation of mRNA cap had a prominent impact on the composition of the protein interactome associated with the RNA bearing mentioned modifications. Further analysis revealed that 2’-*O*-methylation of the second transcribed nucleotide and N6-methylation of adenosine as the first transcribed nucleotide serve as determinants defining transcripts as ‘self’ and contribute to transcript escape from the host innate immune response.

## Introduction

A key feature of eukaryotic mRNA is the presence of a 5’ cap structure, which is indispensable for several biological processes such as pre-mRNA splicing, mRNA export, and translation (1, 2). The 5’ cap of mRNA consists of 7-methylguanosine (m7G) linked by a 5’,5’-triphosphate bridge to the first transcribed nucleotide. This m7GpppN modification, referred to as cap0, is formed enzymatically during transcription. Cap0 can be further modified by nuclear cap 2’-*O*-methyltransferase 1 (CMTR1) to form cap1 (m7GpppNm), wherein the ribose of the first transcribed nucleotide is methylated at the 2’-*O* position (3). Transcripts bearing cap1 can be additionally subjected to ribose 2’-*O*-methylation at the second transcribed nucleotide. This reaction is catalyzed by another methyltransferase, CMTR2, resulting in the formation of cap2 (m7GpppNmpNm) (4). Interestingly, studies have shown that CMTR2 can also act on substrates lacking 2’-*O*-methylation at the m7G-adjacent nucleotide *in vitro* (4), thereby forming an m7GpppNpNm capping structure called cap2-1. In addition to 2’-*O*-methylation, when adenosine is the first transcribed nucleotide, it can be methylated at the N6 position of nucleobase by phosphorylated CTD-interacting factor 1 (PCIF1) to form N6,2’-*O*-dimethyladenosine (m6Am) (5-7). Importantly, transcriptome-wide mass spectrometry-based studies have shown that in some mammalian cells, N6-methyladenosine (m6A) is as abundant as the first transcribed nucleotide as m6Am (8). However, it is unclear whether PCIF1 can add an N6-methyl group to adenosine without prior ribose 2’-*O*-methylation, or conversely, if m6A presence within the cap is a consequence of unknown demethylase activity.

For a long time, the role of extensive methylation of mammalian cap structure was not characterized adequately. Recently, it was reported that the first transcribed nucleotide 2’-*O*-methylation is important for the differentiation between ‘self’ and ‘non-self’ RNA during viral infection in mammalian cells (9-11). Among the sensors of viral RNAs, interferon (IFN)-induced proteins with tetratricopeptide repeats (IFITs) can be distinguished. IFIT1 competes with eukaryotic translation initiation factor 4E (eIF4E) for m7G-capped RNAs, and lack of methylation at the 2’-*O*-position of the first transcribed nucleotide favors IFIT1 binding (12, 13). Cellular mRNAs are protected from IFIT1 sensing by the presence of cap1.

The 5’ cap is a crucial element in mRNA translation, which occurs in a cap-dependent manner. The eIF4F cap-binding complex, containing eIF4E cap-binding protein, promotes translation initiation when bound to the cap (1, 2). The presence of cap0 guarantees efficient mRNA translation, and it was shown that the protein production level may be further elevated by cap1 in particular cell lines (14). Recently, we also found that exogenously delivered mRNAs carrying m6A as the first transcribed nucleotide show increased protein yields in some mammalian cell lines, but only in the context of cap1 (14). However, the impact of 2’-*O*-methylation of the second transcribed nucleotide on translation remains elusive.

mRNA stability is largely controlled by the presence of the cap, as it protects transcripts from exonucleolytic degradation mediated by cellular RNases acting from the 5’-end (1, 2, 15). Enzymes of the XRN1 family are unable to degrade capped mRNAs. Independent of the type of cap modification tested to date (i.e., cap0 and cap1), transcripts are prone to the activity of DCP2, a major cellular decapping enzyme (14), which, in cooperation with its cellular partner DCP1, releases m7GDP and the monophosphorylated RNA chain (15). DXO serves as a quality control factor to eliminate aberrantly capped transcripts and acts as a decapping enzyme. However, in contrast to DCP2, DXO removes the entire cap structure by hydrolyzing the phosphodiester bond between the first and second transcribed nucleotides, resulting in the release of m7GpppN and monophosphorylated RNA, and continues to degrade RNA as an exoribonuclease in the 5’-3’ direction (16). Interestingly, DXO decapping activity was shown to be inhibited by cap1 2’-*O*-methylation (17), suggesting its potential role in distinguishing ‘self’ from ‘non-self’ RNAs. The influence of 2’-*O*-methylation of the second transcribed nucleotide on decapping by DCP2 or DXO is yet to be investigated.

One of the experimental approaches to study the impact of modifications within the mRNA 5’ cap on mRNA-related processes is the preparation of *in vitro* transcribed mRNA (IVT mRNA) and analysis of how the introduced modifications change the biological properties of the transcript. Until recently, IVT mRNA bearing methylations of m7G-adjacent nucleotides were prepared through the use of appropriate methyltransferases (3, 4, 7, 18). However, this method is time-consuming and effort-intensive, requiring the use of recombinant methyltransferases (MTases), preparation of IVT mRNA with an unmodified cap, modifying cap structure with purified enzymes, and analysis of cap modification efficiency after each attempt. Recently, we and other researchers have used a novel approach to obtain IVT mRNA bearing modifications at the first transcribed nucleotide introduced co-transcriptionally, i.e., the use of trinucleotide cap analogs as initiators of *in vitro* transcription (14, 19). However, trinucleotide cap analogs enable the preparation of transcripts with the first transcribed nucleotide modifications. Therefore, to generate capped IVT mRNA that is methylated at the second transcribed nucleotide, i.e., a transcript with cap2 or cap2-1, we generated new molecular tools – tetranucleotide cap analogs chemically decorated with 2’-*O*-methylation at the first and second transcribed nucleotides (cap2), or exclusively at the second transcribed one (cap2-1) – and used these for efficient co-transcriptional RNA capping. These new tools, together with previously synthesized trinucleotide cap analogs (14), enabled us to comprehensively study the impact of cap methylation status, i.e., 2’-*O*-methylation of the first and second transcribed nucleotides, as well as N6 nucleobase methylation of adenosine as the first transcribed nucleotide, on protein production levels in mammalian cells grown under optimal or stress conditions. The result showed that 2’-*O*-methylation of the second transcribed nucleotide regulates protein biosynthesis in a cell-dependent manner. To understand this phenomenon, we characterized the interactomes of transcripts with differently methylated cap structures in mammalian cells originating from different tissues. Moreover, we observed the resistance of RNA capped with cap2 to DXO-mediated decapping and degradation, and vulnerability to DCP2 action. We also showed that 2’-*O*-methylation of the second transcribed nucleotide and N6-methylation of adenosine as the first transcribed nucleotide serve as determinants defining transcripts as ‘self’ and contribute to transcript immune evasion.

## Results

### Tetranucleotide cap analogs enable efficient co-transcriptional capping of RNA with cap2 and cap2-1 structures *in vitro*

To investigate the influence of 2’-*O*-methylation of the second transcribed nucleotide on the biological and biochemical properties of RNA, we required a convenient tool to obtain capped RNAs with this modification. Similar to previously described trinucleotide cap analogs utilized for generating IVT mRNA efficiently capped with cap0 or cap1 (14), we chemically synthesized tetranucleotide cap analogs bearing ribose 2’-*O*-methylation at the first and second transcribed nucleotide m7GpppNmpNm – cap2, or only at the second one m7GpppNpNm – cap2-1. The first transcribed nucleotide was adenosine [cap2(A), cap2-1(A)] or adenosine methylated within the nucleobase at the N6-position [cap2(m6A) and cap2-1(m6A)] (Fig. 1A). By combining these two compounds with the previously synthesized analog of cap1, we created a set of different tri- and tetranucleotide cap analogs combining all possible cap methylation statuses (cap0, cap1, cap2 and cap2-1) with A and m6A (8, 20, 21). Tetranucleotide cap analogs were obtained via a combination of solid-phase and solution chemistry. Highly loaded solid supports generated the trinucleotide 5’-phosphates pN(m)pG(m)pG, which were conjugated with the P-imidazolide of N7-methylguanosine 5’-diphosphate (m7GDP-Im). The product was purified using ion-exchange chromatography followed by reverse-phase high-pressure liquid chromatography (RP-HPLC). The purity and identity of the compounds obtained were confirmed by RP-HPLC and high-resolution mass spectrometry (HRMS), respectively (HPLC profiles and HRMS spectra are provided in the SI Appendix, Table S1).

**Fig. 1.**
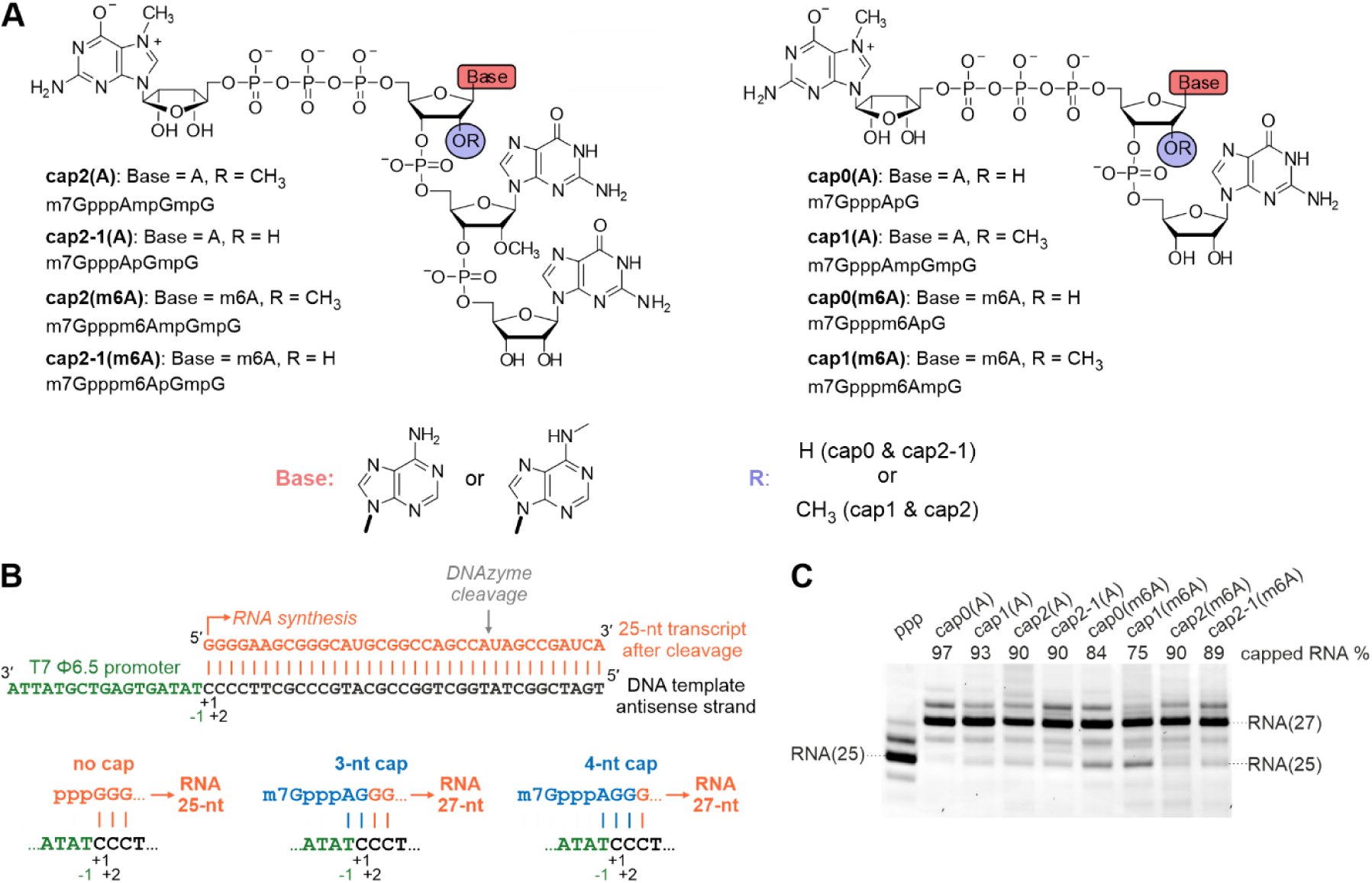
Tetranucleotide cap analogs act as initiators of *in vitro* transcription reaction. **(A)** Structure of cap analogs used in this study, newly synthetized tetranucleotide and previously obtained trinucleotide (14) cap analogs are presented on the left and right side, respectively. **(B)** Comparison of the major transcription initiation events during *in vitro* transcription reaction when either no cap analog, trinucleotide cap analog or tetranucleotide cap analog was used as an initiator. Capped RNA obtained in IVT reaction with tri-or tetranucleotide cap analog is 27-nt long, while uncapped RNA is 25-nt long. **(C)** Analysis of short RNAs obtained by *in vitro* transcription using T7 RNA polymerase in the presence of different cap analogs. The capping efficiency values (percentage) determined by densitometric quantification of the major bands corresponding to capped and uncapped RNA is shown at the top of the gel.

Next, we investigated the efficiency of incorporation of the new tetranucleotide analogs into RNA during *in vitro* transcription compared with the previously tested trinucleotide set. Annealed oligo DNA with a Φ 6.5 promoter sequence followed by 35 nucleotides served as a template for the reaction with T7 RNA polymerase, in which the concentration of the cap analogs was 3-fold higher than that of GTP. The transcripts obtained were HPLC-purified, treated with DNAzyme to trim transcripts at the 3’-end, and separated on a 15% denaturing polyacrylamide gel. Trimmed uncapped transcripts were 25 nucleotides long, whereas the respective RNAs capped with either tri or tetranucleotide analogs were 27 nucleotides long (including m7G), as expected based on the T7 promoter sequence (Fig. 1B). We observed a very high capping efficiency for the new tetranucleotide analogs at 90% for cap2(A), cap2-1(A), and cap2(m6A), and modestly less for cap2-1(m6A) (89%). The highest percentage of capped RNA was achieved in the presence of cap0(A) and cap1(A) trinucleotides (97% and 93%, respectively) (Fig. 1C). Trinucleotides with N6-methylated adenosine were incorporated with slightly reduced efficiencies at 84% and 75% for cap0(m6A) and cap1(m6A), respectively (Fig. 1B), as observed previously (14).

### Differences in cap methylation status affect protein production in a cell-specific manner

Given that new tetranucleotide cap analogs are efficiently incorporated into RNA during *in vitro* transcription reactions, we studied the impact of 2’-*O*-methylation status within the cap structure on protein production levels in mammalian cells. We prepared IVT mRNAs encoding *Gaussia* luciferase capped with differently methylated cap analogs and performed luminescence assays. However, before IVT mRNA can be used for cell transfection, it should be of sufficient purity. Double-stranded (ds) RNA impurities, which together with the uncapped version of studied mRNA could unspecifically interfere with the cell immune system, were of special concern to us (14, 22, 23). Therefore, all capped IVT mRNAs were purified by RP HPLC purification, followed by enzymatic removal of uncapped mRNAs by treatment with 5’-polyphosphatase and Xrn1 (14). Mammalian cells from the human lung carcinoma A549 cell line and murine immature dendritic JAWS II cells were transfected with the purified IVT mRNAs. In A549 cells, cap methylation status had a moderate effect on protein production (Fig. 2A,B). Notably, protein production levels were not affected by the identity of the first transcribed nucleotide (A vs. m6A) in the context of cap0 (Fig. 2B). In contrast, the presence of 2’-*O*-methylation at the first transcribed nucleotide modestly increased the total protein production for IVT mRNAs bearing cap1 relative to transcripts containing respective cap0 versions (Fig. 2A,B). Introduction of the 2’-*O*-methyl group only at the second transcribed nucleotide (cap2-1) had a similar effect on protein biosynthesis as the presence of 2’-*O*-methylation at the first transcribed nucleotide (cap1) (Fig. 2A). Interestingly, protein biosynthesis was 2 and 3-fold higher for mRNA bearing cap2-1(m6A) relative to transcripts with cap1(m6A) or cap0(m6A), respectively (Fig. 2B). Surprisingly, the combination of both 2’-*O*-methylations led to a decrease in protein production compared to mRNAs with the respective cap2-1 versions, such that protein production from mRNA with cap2(A) dropped to the level observed for transcripts with cap0(A) (Fig. 2A). Additionally, the luminescence measured for cells transfected with mRNA bearing cap2(m6A) was almost at the same level as for cells transfected with transcripts bearing cap1(m6A) (Fig. 2B).

Interestingly, differences in cap methylation status had a greater impact on protein production levels in JAWS II than in A549 cells (Fig. 2A-D). As expected, the presence of the 2’-*O*-methyl group at the first transcribed nucleotide significantly increased protein production in murine immature dendritic cells (14) (Fig. 2C,D). mRNAs with cap1(A) provided three times more protein than transcripts with the respective cap0 versions (Fig. 2C), whereas transcripts with cap1(m6A) provided 5.5 times higher luminescence compared to protein levels produced from mRNA with cap0(m6A) (Fig. 2D). In contrast to the results from A549 cells, the presence of N6-methyladenosine as the first transcribed nucleotide in JAWS II cells decreased protein production level 2-fold in the context of cap0 (Fig. 2B,D). The introduction of a methyl group at the second transcribed nucleotide had the opposite effect on protein production in JAWS II relative to A549 cells. Protein biosynthesis from mRNA with cap2-1(A) and cap2-1(m6A) was 56 and 31 times lower compared to the amount of protein produced from transcripts with the cap1 versions (Fig. 2C,D). The presence of both 2’-*O*-methyl groups within the cap structure modestly improved the translational properties of IVT mRNAs compared to transcripts with cap2-1; however, these changes were not statistically significant.

**Fig. 2.**
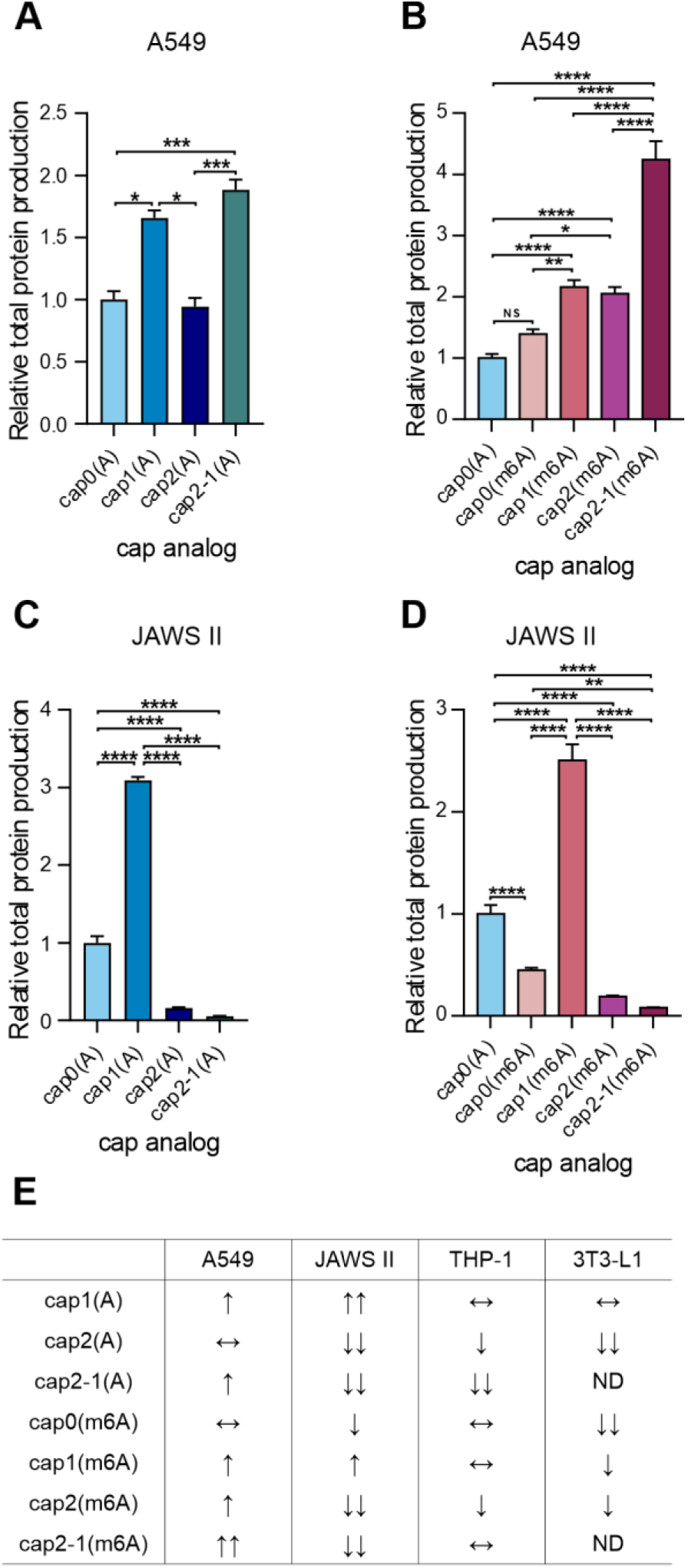
Protein production levels is affected by cap methylation status. Relative total protein production after 72 h measured in the medium from culture of **(A**,**B)** A549 and **(C**,**D)** JAWS II cells transfected with IVT mRNAs encoding *Gaussia* luciferase bearing various cap analogs at their 5’ ends. Bars represent mean value ± SEM normalized to transcripts with cap0(A). Statistical significance: NS – not significant, * P < 0.05, ** P < 0.01, *** P < 0.001, **** P < 0.0001 (one-way ANOVA with Turkey’s multiple comparisons test), (raw data from three independent biological replicates are shown in SI Appendix, Fig. S1, each independent biological replicate consisted of three independent transfections). **(E)** Summary table presenting influence of cap methylation status on protein production levels in A549, JAWS II, THP-1 and 3T3-L1 cells relative to transcript with cap0(A), ↑ – increase of protein production, ↓ – decrease of protein production, double arrow means large (at least 3-time) increase/decrease, ↔ – no significant change.

Given that differences in cap methylation status shape protein production levels in A549 and JAWS II cells, we evaluated the translational properties of differently capped mRNAs and whether these were characteristic of the two cell lines or a general phenomenon due to the presence of a 2’-*O*-methyl group at m7G-adjacent nucleotides. We also analyzed whether N6-methylation of the adenosine nucleobase influenced protein production levels in a cell-specific manner. To examine the impact of cap methylation on protein biosynthesis, we extended our analyses to two more cell lines from different tissues of origin, namely, human THP-1 monocytic cells and murine 3T3-L1 embryonic fibroblasts (SI Appendix, Fig. S3, S4). Among the two tested cell lines, the presence of N6-methyladenosine as the first transcribed nucleotide in the context of cap0 reduced protein production in 3T3-L1 cells alone. Moreover, 2’-*O*-methylation of adenosine as the first transcribed nucleotide did not influence protein production compared to the protein level obtained for mRNA with cap0(A), whereas the same modification in the context of N6-methyladenosine as an m7G-adjacent nucleotide increased the protein production level in 3T3-L1 cells alone. The addition of a second 2’-*O*-methyl group within the cap structure (cap2) decreased protein production, regardless of the identity of the first transcribed nucleotide (A vs. m6A) in both cell lines (SI Appendix, Fig. S3, S4). The only exception was the production of *Gaussia* protein from mRNA with cap2(m6A) in 3T3-L1 cells, where protein levels did not change compared to protein production from mRNA with cap0(m6A) (SI Appendix, Fig. S4). 2’-*O*-methylation of the second transcribed nucleotide alone (cap2-1) influenced protein production levels differently in THP-1 cells depending on the identity of the first transcribed nucleotide (A vs. m6A). Protein production from mRNA bearing cap2-1(m6A) did not change, but was severely impaired for transcripts with cap2-1(A) compared to protein production from IVT mRNA bearing cap0 analogs (SI Appendix, Fig. S3). The change in protein production levels caused by differences in cap methylation for the four tested cell lines is summarized in Fig. 2E.

### RNA-protein interactomes for differently capped transcripts differ substantially in JAWS II dendritic cells but not in A549 cells

To gain insight into why differences in the cap methylation status of reporter mRNA affect protein production in a cell-specific manner, we attempted to identify proteins that interacted with 5’ capped RNAs in mammalian cells. We were particularly interested in understanding how cap structure methylation influences the early stage of protein biosynthesis because m7G is indispensable for cap-dependent translation initiation. Moreover, it is widely assumed that translation regulation depends primarily on the rate-limiting initiation step (24). To identify proteins that bind to differently capped transcripts, we employed affinity purification coupled with mass spectrometry (Fig. 3A). RNAs with different cap structures were biotinylated using a pAp-biotin conjugate (25). Cell lysates were subsequently incubated with the prepared RNAs, and RNP complexes formed on capped transcripts were purified using streptavidin beads. Tandem mass spectrometry followed by label-free quantification revealed protein interactors for differently capped RNAs. In this analysis, we focused on two cell lines, A549 and JAWS II, for which the effect of cap methylation on protein production level was the most extreme (Fig. 2E). Moreover, we studied the influence of only cap 2’-*O*-methylation on protein binding to capped transcripts, as it had the greatest impact on protein biosynthesis (Fig. 2E).

**Fig. 3.**
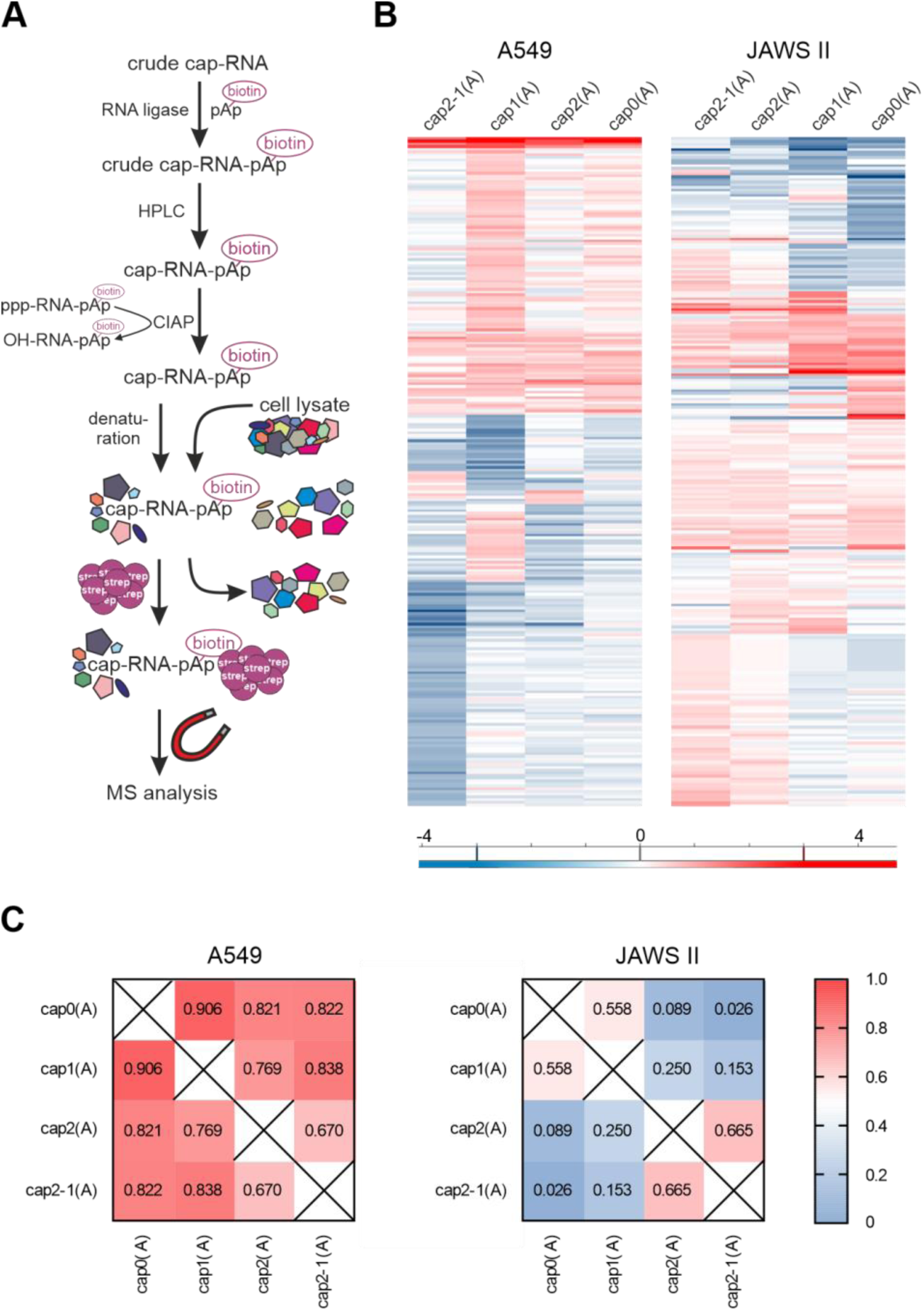
Identification of proteins binding to differently capped transcripts in A549 and JAWS II cell lysates. **(A)** Scheme of affinity purification approach used in this study. **(B)** Heatmap showing hierarchical clustering of proteins performed on log2 ratio values using Euclidean distances generated in Perseus software (42). Ratio of any protein identified and quantified was calculated in comparison to the average level in the mock samples normalized to “1.0” (Heatmaps with marked clusters are presented in SI Appendix, Fig. S4). **(C)** Pearson correlation coefficient for dentified interactomes for differently capped RNA for A549 and JAWS II cells (Exact correlations between identified interactomes for differently capped RNA are presented in SI Appendix, Fig. S5).

In the course of affinity purification experiments, we identified 248 proteins in A549, and 247 in JAWS cell line, that were observed for at least one of cap variant in higher level than in control group (mock) (SI Appendix, Datasets S1 and S2). The data were analyzed to reveal proteins that co-purified with capped transcripts in a recurrent and specific manner compared to the control group. Next, the ratio of an identified protein relative to its average level in the mock samples normalized to “1.0” was determined (Fig. 3B, SI Appendix S4). It was unclear which proteins bound to RNA with 2’-*O*-methylated cap structures among those identified were responsible for modulating protein production levels in JAWS II cells. Therefore, we analyzed the correlation between protein abundance for differently capped RNAs in both cell lines (Fig. 3C, SI Appendix, Fig. S5). Regardless of the 2’-*O*-methylation status, the subset of proteins bound to RNA was quite similar in A549 cells; the Pearson correlation coefficient ranged from 0.670 [cap0(A) vs. cap2(A)] to 0.906 [cap0(A) vs. cap1(A)]. However, the composition of RNA-protein interactomes largely depended on the cap 2’-*O*-methylation status in JAWS II cells, and the Pearson correlation coefficient ranged from 0.026 [cap0(A) vs. cap2-1(A)] to 0.665 [cap2(A) vs. cap2-1(A)]. Interestingly, capped RNA interactomes from JAWS II cells formed two subgroups: 1) interactomes for transcripts bearing an unmethylated cap [cap0(A)] or 2’-*O*-methylated first transcribed nucleotide [cap1(A)], and 2) interactomes identified for capped RNA with a 2’-*O*-methylated second transcribed nucleotide [cap2(A) and cap2-1(A)]. Interactomes in each subgroup were more similar to each other than to any of interactome in the other subgroup.

### The affinity of capped RNA for the translation machinery is not affected by the 5’ end methylation status

To gain a deeper understanding of the influence of 2’-*O*-methylation in the cap structure on protein biosynthesis, we evaluated the binding affinities of differently capped RNAs to the translation initiation factor eIF4E. We used the recently developed microscale thermophoresis (MST) assay for measuring temperature-induced changes in fluorescence for fluorescein amidite-labeled [m7Gp5OC3(5)FAM] upon competitive displacement from eIF4E by the capped transcripts (26). The MST assay confirmed previous results identifying the dissociation constants of eIF4E and trinucleotide cap analogs (14). The presence of 2’-*O*-methylation at adenosine as the first transcribed nucleotide did not influence RNA affinity for eIF4E (Fig. 4A,B). We observed a similar effect for the methylation of the second transcribed nucleotide, regardless of the 2’-*O*-methylation status of the m7G-adjacent nucleotide. Interestingly, we observed a tendency for the capped RNA-eIF4E complex to be slightly destabilized in the presence of m6A as the first transcribed nucleotide, regardless of the cap 2’-*O*-methylation status, except for transcripts with cap2-1 (Fig. 4A).

**Fig. 4.**
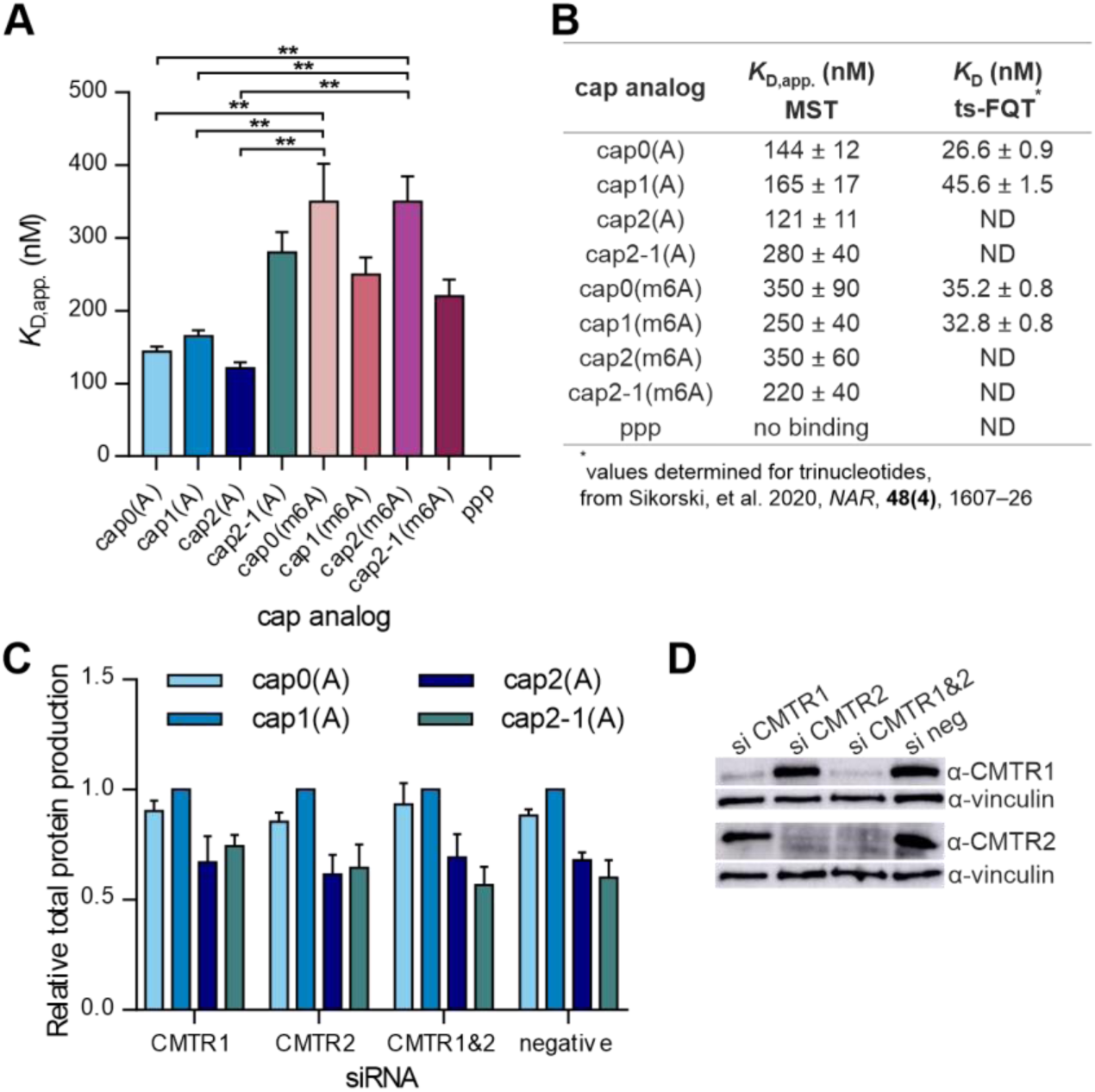
2’-O-methylation within cap structure does not influence RNA affinity to translational machinery. **(A)** Relative affinities of transcripts bearing different cap analogs for murine eIF4E determined using microscale thermophoresis (MST). Bars represent mean value ± SEM from three independent replicates. Statistical significance: ** P < 0.01 (one-way ANOVA with Turkey’s multiple comparisons test) **(B)** Comparison of apparent binding constant values *K*_D,app_ for capped RNAs in complexes with murine eIF4E measured with MST with dissociation constants of eIF4E-cap complexes obtained using time-synchronized fluorescence quenching titration (ts-FQT) (14). **(C)** Competition of differently capped IVT mRNA encoding *Gaussia* luciferase with endogenous mRNAs for translation machinery. A549 cells were pre-treated with siRNA targeting CMTR1 or 2, or both (negative siRNA control was utilized in parallel) for 48 h; then, cells were transfected with IVT mRNA and after 16 h medium was collected for luciferase activity analysis. Bars represent mean value ± SEM normalized to *Gaussia* luciferase activity measured for mock-treated cells. **(D)** Western blot analysis: detection of CMTR1, CMTR2 and vinculin in lysates from siRNA-treated A549 cells from **(C)**.

Next, to validate the biochemical data, we investigated the competition between exogenously delivered reporter transcripts and endogenous mRNAs for the translation machinery. We were interested in studying whether depriving endogenous mRNAs of cap 2’-*O*-methylation(s) would result in an increase in protein production levels for any of the studied IVT mRNAs. To this end, we knocked down the genes encoding CMTR1, CMTR2, or both. CMTR1 and CMTR2 are methyltransferases responsible for 2’-*O*-methylation of the first and second transcribed nucleotides, respectively (3, 4). Forty-eight hours after siRNA transfection, differentially capped mRNAs encoding *Gaussia* luciferase were introduced into A549 cells via lipofection. After another 16 h, the growth medium was evaluated for luminescence activity (Fig. 4C), and cell lysates were used to study the effectiveness of CMTR depletion by western blotting (Fig. 4D). Irrespective of the depleted methyltransferase(s), none of the tested IVT mRNAs preferentially interacted with the cellular translation machinery (Fig 4C). Thus, these data were consistent with our MST measurements.

### The second transcribed nucleotide 2’-*O*-methylation prevents RNA from decapping by hDXO but not by hDCP2

The presence of a cap does not only enable efficient translation but also protects transcript 5’ end from degradation. Thus, we investigated whether differences in protein production from reporter mRNAs may partially arise from the different susceptibilities of transcripts to decapping. In mammalian cells, the DCP1/DCP2 heterodimer, where DCP2 is a catalytically active subunit, is the main factor responsible for cap structure elimination from transcripts designated for degradation (15). To investigate the susceptibility of RNAs bearing differently methylated caps, short transcripts capped with tri and tetranucleotide cap analogs were prepared. Capped RNAs were then incubated with recombinant active and catalytically inactive versions of the human DCP2 (hDCP2) protein (SI Appendix, Fig. S7A) followed by polyacrylamide gel analysis of the products collected over time (Fig. 5A). We observed a reduction in the band intensity corresponding to capped transcripts with time, and the appearance of bands representing transcripts deprived of m7GDP. Importantly, this process was observed for active hDCP2 alone, indicating that decapping activity can be attributed solely to hDCP2. Moreover, to directly compare the susceptibility to hDCP2, the fraction of capped RNAs that remained in each sample was plotted as a function of time (Fig. 5B-D). The presence of one or two 2’-*O*-methylation(s) within cap did not affect susceptibility to hDCP2 activity *in vitro*. This result is in agreement with our previous studies, in which we showed that 2’-*O*-methylation of the first transcribed nucleotide did not influence the vulnerability of transcripts to decapping, regardless of the identity of the first transcribed nucleotide (14). However, we observed that the presence of N6-methylated adenosine at the m7G-adjacent position modestly increased RNA susceptibility to hDCP2 activity, irrespective of the cap 2’-*O*-methylation status.

**Fig. 5.**
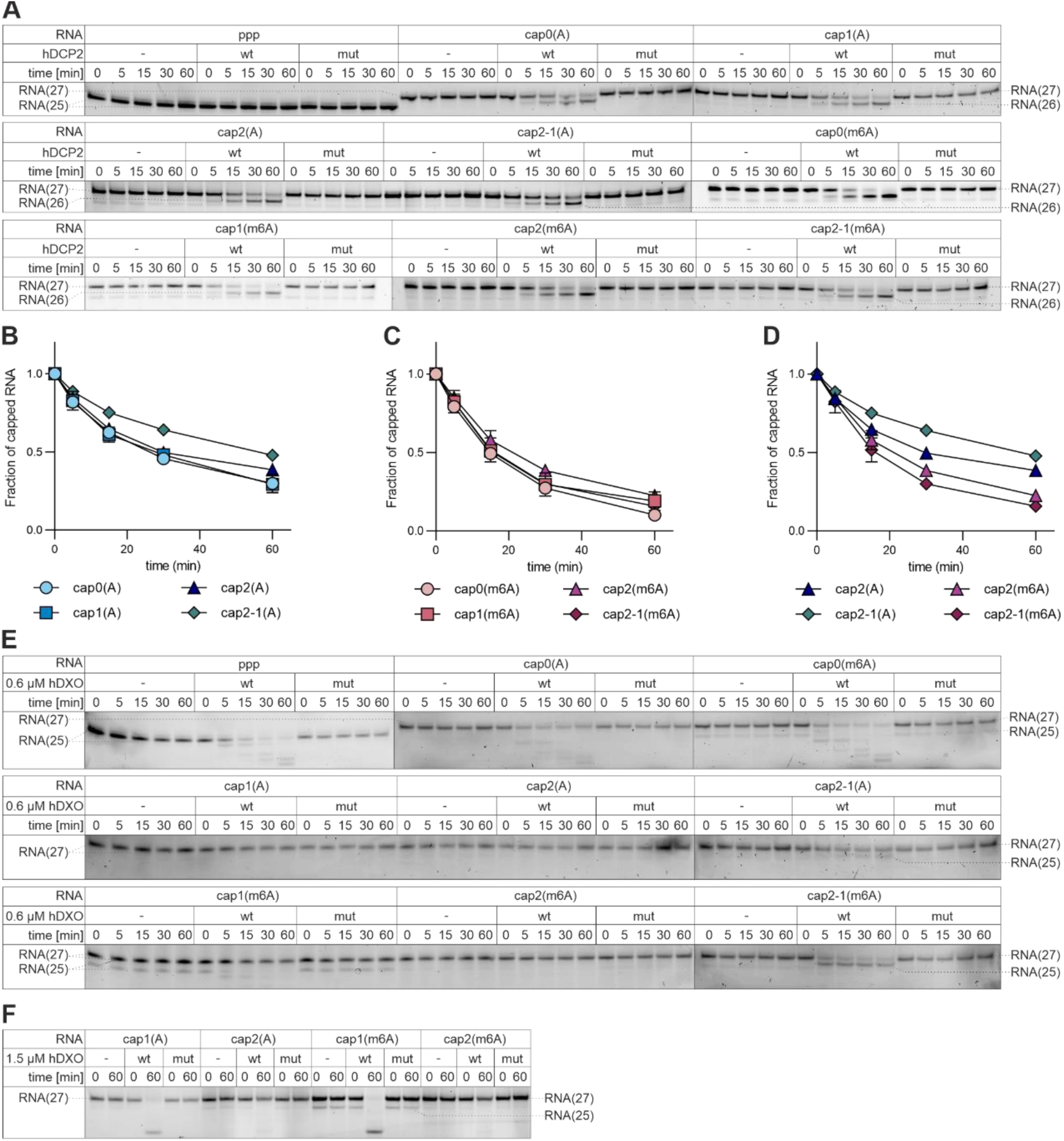
2’-*O*-methylation of the second transcribed nucleotide prevents RNA from decapping by DXO but not by DCP2. **(A-D)** Short capped RNAs were subjected to treatment with hDCP2 (wild type or mutant) over 60 min time-course. Reactions without enzyme served as controls. Aliquots from indicated time points were resolved in polyacrylamide gel and bands corresponding to capped transcripts (27-nt long) and to RNAs decapped by hDCP2 action (26-nt long) were quantified densitometrically. **(A)** Representative polyacrylamide gel analyses obtained for all tested capped RNAs (two additional repetitions with wild type hDCP2 of this experiment are shown in SI Appendix, Fig. S7). **(B–D)** Quantitative results for all studied RNAs. The fraction of capped RNA remaining in the total RNA was plotted as a function of time. Data points represent mean values ± SD from triplicate experiments. **(E)** Short capped RNAs were subjected to treatment with hDXO (wild type or mutant) over 60 min time-course. Reactions without enzyme served as controls. Aliquots from indicated time points were resolved in polyacrylamide gel. **(F)** Experimental setup as in **(E)**, however 2.5-fold higher hDXO concentration was used.

Another enzyme involved in decapping of mammalian transcripts is the DXO protein (16). In contrast to DCP2, DXO does not only decap transcripts, but also has 5’-3’ exonucleolytic activity, through which it can eliminate RNA designated for degradation. Moreover, DXO accepts uncapped 5’-triphosphorylated RNA as a substrate, and unlike DCP2, it cleaves off the entire cap structure, releasing dinucleotides linked via the triphosphate bridge. Recently, a study showed that 2’-*O*-methylation present in cap1 confers RNA resistance to human DXO (hDXO) decapping activity and further degradation (17). Moreover, the presence of the 2’-*O*-methyl group in RNA bodies hampers hDXO exonucleolytic activity. Thus, we hypothesized that the simultaneous presence of 2’-*O*-methylation at the first and second transcribed nucleotides can make RNAs more resistant compared to transcripts with cap1. To evaluate the hypothesis, we performed *in vitro* activity assays using active or catalytically inert versions of the recombinant hDXO protein (SI Appendix, Fig. S7B) and differently capped short RNAs. The reaction products from the time course were resolved using polyacrylamide gels (Fig. 5E). During the reaction, we observed decapping and subsequent degradation of RNA bearing cap0, regardless of the identity of the m7G-adjacent nucleotide (A vs m6A). As expected, the presence of the 2’-*O*-methyl group at the first transcribed nucleotide alone or both the first and second transcribed nucleotides protected transcripts from hDXO activity. Interestingly, we did not observe a decrease in decapping rate for transcripts with 2’-*O*-methylation at the second transcribed nucleotide alone; however, the presence of a methyl group at this position efficiently blocked exonucleolytic degradation. Moreover, we attempted to differentiate the susceptibility of transcripts with cap1 and cap2 to degradation by employing higher hDXO concentration. Under these experimental conditions, RNA bearing 2’-*O*-methylation at both the first and second transcribed nucleotides was more resistant to hDXO action than transcripts with cap1 (Fig. 5F).

### 2’-*O*-methylation of the second transcribed nucleotide and N6-methylation of adenosine as the first transcribed nucleotide contribute to RNA immune evasion

2’-*O*-methylation of the first transcribed nucleotide is a known mark distinguishing ‘self’ from ‘non-self’ transcripts (9, 10). IFIT proteins are the main factors responsible for regulating cap-dependent translation in stressed cells (13, 27). IFIT1 binds directly to the mRNA 5’ end and competes with eIF4E for binding to the N7-methylguanosine triphosphate cap. The 2’-*O*-methylation in cap1 prevents mRNA and IFIT1 binding and translational inhibition by this protein (12, 13). However, methylation of the second transcribed nucleotide may also contribute to this process (28). To evaluate this hypothesis, we examined the protein production levels in stressed cells transfected with IVT mRNAs encoding *Gaussia* luciferase bearing differently methylated caps. To induce stress conditions and cause upregulation of genes involved in innate immune response, which influence translation, we performed interferon (IFN) treatment, which is widely used to increase the production of antiviral proteins, including IFITs (13, 29). To ascertain whether our experimental setup induced upregulation of antiviral protein production, we verified the expression of several antiviral genes in the presence of increasing amounts of IFNα by RT-qPCR (SI Appendix, Fig. S9). Our results showed that upregulation of IFITs, 2’-5’-oligoadenylate synthetase 1 (OAS1), and protein kinase R (PKR) correlated positively with the amount of IFNα added to the cell culture medium.

Next, we examined the changes in protein production levels for differently capped IVT mRNA encoding *Gaussia* luciferase in A549 cells treated with increasing amounts of IFNα (Fig. 6A,B and SI Appendix, Fig. S10). As IFNα concentration increased, we observed a gradual decrease in protein production for all tested IVT mRNAs (SI Appendix, Fig. S10); however, the greatest drop in protein biosynthesis was observed for transcripts bearing cap0. As expected, 2’-*O*-methylation of the first transcribed nucleotide was associated with greater protein production levels compared to transcripts with the respective cap0 structures (Fig. 6A,B and SI Appendix, Fig. S10). Intriguingly, the effect of 2’-*O*-methylation of the second transcribed nucleotide depended on the identity of the first transcribed nucleotide (A vs. m6A). If adenosine was the first transcribed nucleotide [cap2(A)], the presence of two 2’-*O*-methylations within the cap resulted in the least reduction in *Gaussia* luminescence. In contrast, the combination of two 2’-*O*-methyl groups and N6-methyladenosine [cap2(m6A)] caused a decrease in protein production compared to IVT mRNA bearing the respective cap1 structure. Moreover, experiments with IVT mRNA containing cap2-1 structure indicated that the presence of 2’-O-methylation at the second transcribed nucleotide alone was sufficient to ensure high protein production levels under stress conditions. Protein production from transcripts with cap2-1(A) was similar to that from IVT mRNA bearing cap1(A), whereas *Gaussia* luminescence measured for cells transfected with transcripts bearing cap2-1(m6A) was comparable to that from cells transfected with IVT mRNA with cap2(m6A).

**Fig. 6.**
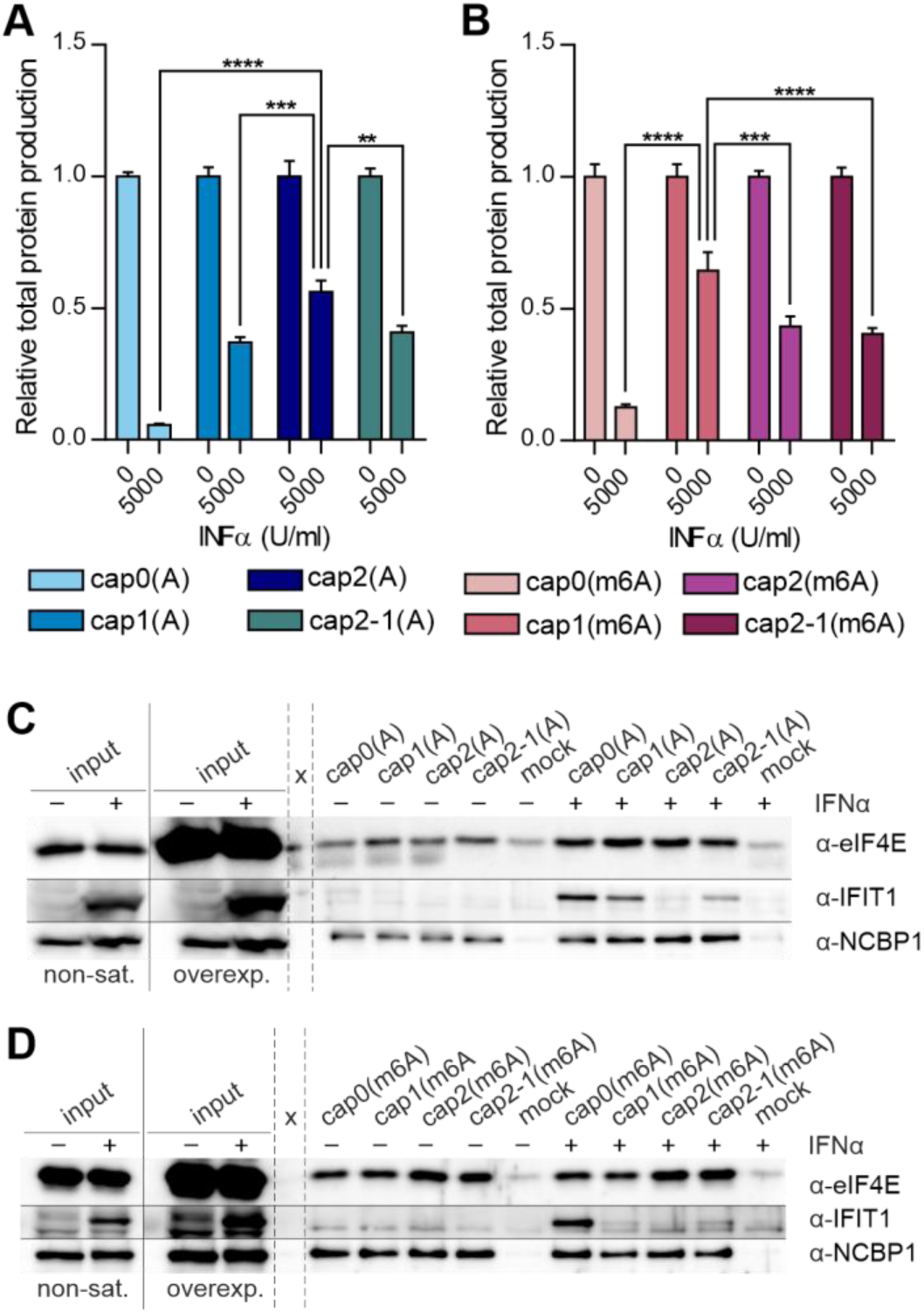
Methylation of the first transcribed nucleotides in mRNA only partially rescues translation shutdown under IFN-induced stress. **(A,B)** Relative protein production levels 4 hours after 5 hour IFNα pre-treatment in A549 cells. Bars for each transcript represent mean value ± SEM normalized to protein production in mock treated cells. Statistical significance: * P < 0.05, ** P < 0.01, *** P < 0.001, **** P < 0.0001 (one-way ANOVA with Turkey’s multiple comparisons test) (data presenting changes in relative protein production with increasing concentration of IFNα are shown in SI Appendix, Fig. S9). Co-purification of endogenous proteins from lysates of IFNα treated A549 cells with biotinylated RNA bearing cap0, cap1, cap2 or cap2-1 with **(C)** A or **(D)** m6A as the first transcribed nucleotide. Human eIF4E, IFIT1 and NCBP1 were detected in precipitates by western blotting.

Because IFIT1 competes with eIF4E for mRNAs in cells exposed to stress stimuli, we analyzed the contribution of each cap methylation type to IFIT1 binding. To this end, we used the previously mentioned approach based on biotinylation of differently capped transcripts as baits for proteins from lysates prepared using IFNα-treated and untreated A549 cells. RNP complexes formed on capped transcripts were purified using streptavidin beads and IFIT1 levels were analyzed by western blotting. As expected, the presence of 2’-*O*-methylation at the first transcribed nucleotide [cap1(A)] decreased the affinity of capped RNA for IFIT1 (Fig. 6C). The 2’-*O*-methylation of only the second transcribed nucleotide [cap2-1(A)] had a similar impact on IFIT1 binding. Importantly, IFIT1 was unable to bind to RNA bearing cap2(A). 2’-*O*-methylation did not affect eIF4E binding or interaction with the nuclear cap-binding complex. These results were in line with the protein production levels measured for differently capped reporter transcripts in IFNα-treated A549 cells (Fig. 6A,B and SI Appendix, Fig. S10). Interestingly, for transcripts with m6A as the first transcribed nucleotide, the presence of only one 2’-*O*-methylation made RNA almost unrecognizable for IFIT1 (Fig. 6D), indicating that the identity of the m7G-adjacent nucleotide affects RNA recognition by a major effector of the cellular immune defense system. Moreover, 2’-*O*-methylation of the second transcribed nucleotide resulted in either a synergistic reduction in IFIT1 binding [cap2(m6A)] or ensured similar affinity towards IFIT1 [cap2-1(m6A)] when compared to RNA bearing cap1(m6A). Taken together, 2’-*O*-methylation of the second transcribed nucleotide as well as N6-methylation of adenosine as the first transcribed nucleotide may serve as determinants defining transcripts as ‘self’, and contribute to transcript escape from the host innate immune response.

## Discussion

*In vitro* transcribed mRNA can be used as a therapeutic agent in preventive vaccines, cancer immunotherapy, or gene replacement therapy (30-32). However, the utility of IVT RNA exceeds potential clinical applications. IVT RNA can also serve as valuable tools for studying RNA-related processes *in vitro* and *in vivo* (33). Preparing IVT RNA with different modifications enables the analysis of their impact on biological processes of interest. Among the modifications that can be studied using IVT RNA are those in the cap structure, i.e., ribose 2’-*O*-methylation(s) of the first and second transcribed nucleotide(s), as well as N6 nucleobase methylation of adenosine when present as the first transcribed nucleotide. Importantly, the use of differently capped transcripts enables not only a straightforward comparison of how each cap methylation influences RNA-related processes but also enables the study of the interplay between selected modifications. Until recently, the introduction of modifications in the cap structure of IVT RNA was a time-consuming and elaborate process. For instance, transcripts with N6,2’-*O*-dimethyladenosine as the first transcribed nucleotide and bearing 2’-*O*-methylation at the second transcribed nucleotide were obtained using three consecutive enzymatic reactions. Moreover, the generation of such transcripts requires a detailed analysis of cap modification efficiency after each enzymatic step. Recently, the process of obtaining RNA bearing modifications in the cap structure has been considerably simplified using new molecular tools developed by us and other researchers, i.e., trinucleotide cap analogs as initiators of *in vitro* transcription reactions, that enable preparation of transcripts with modifications at the first transcribed nucleotide (14, 19). However, the application of trinucleotide cap analogs does not enable straightforward generation of transcripts bearing 2’-*O*-methylation at the second transcribed nucleotide. To fully appreciate the impact of cap methylation on RNA-related processes, we have described the generation of new molecular tools, namely tetranucleotide cap analogs, which enable direct preparation of RNA possessing cap 2’-*O*-methylated also at the second transcribed nucleotide.

Combining the previously generated tri (14) and newly synthesized tetranucleotide cap analogs described herein, we were able to systematically and comprehensively study the impact of all three known cap methylations on protein biosynthesis. Our data showed that the presence of 2’-*O*-methylation at the first transcribed nucleotide (cap1) either boosted (A549 and JAWS II cell lines) or did not change (THP-1 cells) protein production levels compared to those obtained for IVT mRNA bearing the respective cap0 counterparts, regardless of the identity of the first transcribed nucleotide. The only exception for cap1 relative to cap0 was observed in the 3T3-L1 cell line, where the protein production level for transcripts with adenosine as m7G-adjacent nucleotide was unchanged, whereas mRNA with cap1(m6A) showed a modestly greater protein yield than transcript bearing cap0(m6A). These results are in line with our previous observations showing that 2’-*O*-methylation enhances the biosynthesis process in JAWS II but not in HeLa or 3T3-L1 cells (14). Importantly, our data are also consistent with recent studies on the role of PCIF1, a cap methyltransferase responsible for N6-methylation of 2’-*O*-methylated adenosine as the first transcribed nucleotide, in mRNA metabolism (6, 7). Sendinc et al. and Boulias et al. showed that the loss of PCIF1 resulted in either an increase or no change in the translation of transcripts, which naturally possess m6Am in the cap structure, in the MEL624 human melanoma cell line (7) and human embryonic kidney 293T cells, respectively (6). We observed that change of A to m6A in context of cap1 led to decrease in protein production (3T3-L1 and JAWS II cells) or did not affect protein biosynthesis at all (THP-1 and A549 cells).

The presence of the 2’-*O*-methyl group at the first and second transcribed nucleotides (cap2) resulted in decreased protein production levels in the tested cell lines compared to levels for transcripts bearing the respective cap1 counterparts. However, the magnitude of reduction depended on the identity of the first transcribed nucleotide. Protein production levels for transcripts with cap2(m6A) were less affected than for mRNA with cap2(A) relative to protein levels for transcripts bearing the respective cap1 analogs. Interestingly, in A549 cells, the occurrence of two 2’-*O*-methylations in the context of N6-methyladenosine ensured the same level of protein production as the yield from mRNA with cap1(m6A). Intriguingly, our results do not support the hypothesis formulated in the 1970s that mRNAs with cap2 show the highest translation rate (34). Perry and Kelley used murine L fibroblasts in their studies (34), and we employed a similar cell line, i.e., murine 3T3-L1 embryonic fibroblasts, in our research. However, we observed a reduction in protein production levels using IVT mRNA with cap2 regardless of the identity of the first transcribed nucleotide (A vs. m6A) compared to protein levels for transcripts bearing the respective cap1 analogs. Taken together, our results indicate that protein production regulation via cap methylation is cell line-dependent. Moreover, the 2’-*O*-methylation of the second transcribed nucleotide is the major player in this mechanism.

To understand why protein production levels differed substantially among the tested cell lines, we sought to identify proteins that interacted with differently capped transcripts in A549 and JAWS II cells. We analyzed why the presence of 2’-*O*-methylation at the first transcribed nucleotide (cap1) boosted protein production in JAWS II, whereas two 2’-*O*-methylations at the first and second transcribed nucleotides (cap2) hampered protein biosynthesis in this cell line. In contrast, the same set of modifications modestly affected protein production levels in A549 cells. Among the identified cap-interacting proteins, we could not indicate factors associated with transcripts bearing a 2’-*O*-methylated cap structure that impacted protein production levels in JAWS II cells. However, a meta-analysis of the identified proteins revealed that RNA-protein interactomes for differently capped transcripts varied substantially in JAWS II cell lysates, whereas in A549 cell lysates, interactomes were similar. The biggest discrepancy among interactomes of capped RNAs was identified in JAWS II cells for transcripts differing in the 2’-*O*-methylation status of the second transcribed nucleotide. The set of proteins associated with RNA bearing either cap2(A) or cap2-1(A) was more similar to each other than to the interactomes identified for transcripts with cap1(A) or cap0(A). Interestingly, the interactomes of transcripts bearing cap0(A) and cap1(A) from JAWS II cells were very similar even though a significant difference in reporter protein production level was observed between them. Based on these results, we can assume that only protein biosynthesis was repressed (cap0/cap1 vs. cap2/cap2-1) at the level of transcript 5’ end recognition or translation initiation. In contrast, an increase in protein production due to 2’-*O*-methylation of the first transcribed nucleotide (cap0 vs. cap1) was rather not caused at the level of translation initiation. Interestingly, similar proteomic analyses have recently been performed by other researchers to characterize proteins associated with RNA bearing differently modified 5’-ends; however, in contrast to our results, Habjan et al. compared uncapped RNA to its counterparts bearing only cap0 or cap1 (13). Nevertheless, the results of MS/MS analysis from their study are in line with our observations, i.e., no difference was observed in the identified capped RNA-protein interactomes in human cervical carcinoma (HeLa) and mouse embryonic fibroblast (MEF) cell lines. Taken together, proteomics studies support the hypothesis that cap methylation status influences protein biosynthesis in a cell-dependent manner, and we show that the major determinant affecting protein production is 2’-*O*-methylation of the second transcribed nucleotide.

As protein production is affected differently in various cell lines by the cap methylation status, and 2’-*O*-methylation of the second transcribed nucleotide is the major player in this mechanism, we asked whether this modification could influence the affinity of transcripts towards the translation initiation machinery. Previously, we showed that none of the modifications of the first transcribed nucleotide affected the stability of the cap analog-eIF4E complex (14). In this study, we expanded this analysis to transcripts with all possible methylation combinations within the cap structure and investigated RNA affinity towards eIF4E. We found that none of the studied cap methylations strongly influenced the stability of the RNA-eIF4E complex. The significance of these findings was further strengthened by competition experiments, in which exogenous reporter transcripts competed with endogenous mRNAs for the translation machinery in cells depleted of cap 2’-*O*-methyltransferases. Knockdown of cap methyltransferases (CMTR1 and CMTR2, either individually or simultaneously) did not influence protein production levels for differently capped IVT mRNAs, indicating that regardless of the cap 2’-*O*-methylation status, the translation machinery showed no preference for the different transcript types. Therefore, to explain how the cap methylation status modulates RNA-related processes, we investigated transcript stability.

The cap not only ensures efficient protein biosynthesis, but also protects RNA 5’ end from uncontrolled exonucleolytic attack. DCP2 is the main decapping enzyme responsible for cap elimination from transcripts designated for degradation (15). Biochemical characterization revealed that the cap methylation status did not affect RNA susceptibility to decapping by hDCP2. However, transcripts bearing adenosine as the first transcribed nucleotide were slightly more resistant to hDCP2 action than RNA with N6-methyladenosine as the first transcribed nucleotide, regardless of the cap 2’-*O*-methylation status. This difference in susceptibility of transcripts to hDCP2 activity is supported by recent genome-wide studies in 293T cells, in which the level of transcripts, which naturally possess m6Am in the cap structure, was increased upon PCIF1 knockout (5). However, these observations have been questioned by other researchers. Sendinc et al. reported no changes in transcription or mRNA stability upon PCIF1 knockout in MEL624 cells (7). Intriguingly, we noticed that 2’-*O*-methylation impacted susceptibility to decapping by the hDXO enzyme. The presence of only one 2’-*O*-methyl group impaired hDXO action on the RNA bearing cap1 structure compared to cap0, regardless of the identity of the first transcribed nucleotide (A vs. m6A). Our observation is in line with a previous report by Picard-Jean et al., which showed that 2’-*O*-methylation of the first transcribed nucleotide impaired transcripts’ susceptibility to decapping by hDXO. Moreover, Picard-Jean et al. also noted that internal 2’-*O*-methylation drastically reduced hDXO exonucleolytic activity towards transcripts with such modifications within the RNA body (17). We found that although 2’-*O*-methylation of only the second transcribed nucleotide (cap2-1) did not affect hDXO decapping activity compared to its action on RNA with cap0, subsequent exonucleolytic degradation of the decapped transcript was impaired. Thus, we envisioned that the decapping rate of transcripts with cap2 would be similar to that for RNAs with cap1 analogs; however, 2’-*O*-methylation of the second transcribed nucleotide can block subsequent exonucleolytic hydrolysis. Interestingly, we found that the decapping rate was reduced for transcripts containing cap2 relative to RNAs bearing cap1 analogs. Apart from playing a role in the removal of improperly capped transcripts (15), DXO was recently recognized as an antiviral factor responsible for the restriction of hepatitis C virus (HCV) replication (35). Therefore, we concluded that 2’-*O*-methylation of the second transcribed nucleotide may be an additional determinant of molecule ‘selfness’ that enables endogenous transcripts to escape from being recognized by the innate immune response.

It is widely assumed that 2’-*O*-methylation of the first transcribed nucleotide is not sufficient to completely protect mRNAs from being recognized as ‘non-self’ by the cellular immune defense system (28, 36, 37) Therefore, it is an open question what are the other RNA features that mark transcripts as ‘self-ones’. One such feature could be an additional 2’-*O*-methylation present at the second transcribed nucleotide. Abbas et al. showed that only transcripts bearing caps with both m7G-adjacent nucleotides 2’-*O*-methylated are fully resistant *in vitro* to IFIT1 recognition (28), which plays a crucial role in distinguishing ‘self’ from ‘non-self’ RNAs (38). Moreover, based on the crystal structure of the IFIT1 in complex with capped RNA, the authors proposed that the presence of N6,2’-*O*-dimethyladenosine instead of 2’-*O*-methyladenosine could also provide full protection against IFIT1 binding; however, they did not provide any experimental data to support this hypothesis (28). Tartell et al. recently showed that the presence of m6Am, but not Am, as the first transcribed nucleotide of vesicular stomatitis virus (VSV) and rabies virus (RABV) transcripts enabled efficient viral replication, presumably by impeding the recognition of viral RNAs by the effectors of the innate immune system (39). Using interferon treatment in A549 cells, we showed that protein production for IVT mRNA bearing cap2 was less affected than for transcripts with cap1; however, this effect was dependent on the identity of the first transcribed nucleotide. The combination of N6-methylation and 2’-*O*-methylation of adenosine as the first transcribed nucleotide ensured the highest protein production level under stress conditions. However, N6-methylation of adenosine as the first transcribed nucleotide did not guarantee a high level of protein biosynthesis in stressed cells, and the reduction in protein production was comparable for IVT mRNA bearing cap0(m6A) and cap0(A). The result indicates that 2’-*O*-methylation is indispensable for efficient protein production under stress conditions, whereas N6-methylation positively stimulates protein biosynthesis only if adenosine is 2’-*O*-methylated. Moreover, we found that the presence of a single 2’-*O*-methyl group, regardless of the position of the modified nucleotide, was sufficient to ensure high protein production levels in stressed cells. This observation could be crucial for understanding how cell fitness is regulated by exposure to stress. It was shown that CMTR1 controls mRNA stability and promotes protein expression of certain IFN-induced genes in stressed cells (40, 41). The moderate influence of the lack of CMTR1 function on protein production could result due to redundancy in function between cap 2’-*O*-methylation at different positions. We can assume that endogenous transcripts can be efficiently translated even when lacking 2’-*O*-methylation of the first transcribed nucleotide; however, in such cases, the modification should be present at the second transcribed nucleotide.

Finally, we found that additional methylations of cap1(A) counteracted the recognition of transcripts by IFIT1. The presence of either cap2(A) or cap1(m6A) causes transcripts to be undetectable by IFIT1. Interestingly, the position of the 2’-*O*-methyl group, whether linked to the first or second transcribed nucleotide, did not influence the efficiency of RNA escape from recognition by IFIT1. This observation is consistent with the results of our protein production studies in stressed cells. Taken together, both 2’-*O*-methylation of the second transcribed nucleotide and N6-methylation of adenosine as the first transcribed nucleotide may serve as additional determinants defining transcripts as ‘self-ones’.

To summarize, we have shed light on the role of RNA cap methylation. The new tools presented here enabled the generation of efficiently capped transcripts with structures modified in a defined manner. We demonstrated the impact of specific modifications and their combinations on protein production levels, which may differ in various cell lines. We also investigated an interplay between cap methylations in the context of particular biochemical features of capped transcripts. We postulate that the combination of cap methylations enhances the identity of RNA as a ‘self-molecule’ for cellular recognition factors that contribute to immune evasion.

## Materials and Methods

**Oligonucleotides and plasmids** used in the study are listed in SI Appendix, Table S2 and S3, respectively.

### Synthesis of cap analogs

Tetranucleotide cap2 (m7GpppNmpGmpG) and cap2-1 (m7GpppNpGmpG) analogs were obtained by combining solid-phase and solution chemistry as described previously (13). Detailed procedure is provided in SI Appendix.

### RNA preparation

Short and messenger RNAs (encoding *Gaussia* luciferase) were obtained by *in vitro* transcription using annealed oligo DNA and linearized plasmid as a template, respectively. Transcripts were capped with different tri-or tetranucleotide analogs co-transcriptionally. RNAs were HPLC purified. Detailed protocol is described in SI Appendix.

### Cell lines and culture conditions

A549 human carcinoma lung cells (ATCC CCL-185), murine immature dendritic cell line JAWS II (ATCC CRL-11904), THP-1 human monocytes (ATCC TIB-202) and 3T3-L1 murine embryonic fibroblasts (ATCC CL-173) were used in the study. Cells were cultured in optimal dedicated media for normal growth conditions, while for stress conditions Universal Type I Interferon (IFNα) in particular concentrations was added. Details are provided in SI Appendix.

### Reporter mRNA translation in cells – luminescence assay

For translation studies 1×10^4^ cells were seeded at the day of experiment in 100 μl medium/well of 96-well plates, with particular concentration of IFNα if applicable. Cells were transfected with the mixture of HPLC-purified mRNA and Lipofectamine MessengerMAX Transfection Reagent in Opti-MEM. Activity of secreted luciferase was examined in medium collected 72 h after transfection, utilizing h-coelenterazine. Luminescence assay was also combined with silencing of CMRT1/2 genes using siRNAs. Detailed procedures are described in SI Appendix.

### Affinity purification with mass spectrometry and western blot analysis

Affinity purification of capped RNA-protein interactomes from A549 and JAWS II cell lysates was performed utilizing biotinylated, differently capped short RNA and streptavidin magnetic beads. Obtained complexes were analyzed by tandem mass spectrometry. Smaller scale affinity purification was conducted to assess abundance of eIF4E, IFIT1 and NCBP1 proteins by western blotting. Details of experimental procedures and MS data analysis are described in SI Appendix.

### eIF4E affinity assay – microscale thermophoresis (MST)

Microscale thermophoresis-based method, described previously in (25), with modifications, was applied to determine dissociation constant for complex encompassing eIF4E protein and capped short RNA. Fluorescent probe, murine eIF4E and capped short RNAs as ligands were utilized. Detailed protocol is available in SI Appendix.

### Recombinant hDCP2 and DXO cloning, overproduction and purification

Wild-type (WT) DXO and catalytically inactive variants of both hDCP2 and hDXO were cloned and expressed from pET28 vectors. Plasmid encoding for WT variant of hDCP2 was a kind gift from Prof. Megerditch Kiledjian. WT and mutant hDCP2 and hDXO proteins were overproduced in *E. coli* BL21-CodonPlus(DE3)-RIL strain and purified employing nickel affinity chromatography. Detailed protocols are described in SI Appendix.

### RNA decapping assays

Short capped RNAs were subjected to decapping assays with human DCP2 and DXO enzymes. Detailed procedure is described in SI Appendix.

### RT-qPCR

Following IFNα treatment, cells were lysed and lysates were then used for reverse transcription. First-strand cDNA served subsequently as a template for quantitative PCR to assess differences in expression level of stress-induced genes: *IFIT1, IFIT2, IFIT3, RIG-I, MDA5, PKR, OAS*, and *GAPDH* as a reference gene. Protocol is described in SI Appendix.

## Acknowledgments

We thank Megerditch Kiledjian (Rutgers University) for providing phDCP2(1-350)wt DNA construct. This work was supported by National Science Centre (2018/31/D/NZ1/03526 to PJS, 2019/33/B/ST4/01843 to JJ, and 2017/26/E/NZ1/00724 to RT).

