## Supplementary material for "2’-*O*-methylation of the second transcribed nucleotide within mRNA 5’ cap impacts protein production level in a cell specific manner and contributes to RNA immune evasion": SI Appendix

**Detailed Material and Methods**

**Synthesis of cap analogs**

Tetranucleotide cap2 (m7GpppNmpGmpG) and cap2-1 (m7GpppNpGmpG) analogs were synthesized according to the procedure described earlier (1). Briefly, trinucleotide 5'-phosphates pNmpGmpG and pNpGmpG were synthesized by phosphoramidite method on a high-loaded solid support (DMT-2'-*O*-TBDMS-rG$^{iBu}$ 3' lcaa PrimerSupport 5G, 308 µmol/g) using ÄKTA Oligopilot plus 10 synthesizer (GE Healthcare) or in a syringe, respectively. When using the manual method, the support was loaded directly into a syringe and the reagents and solvents were flowing through the support. Additionally, when the coupling steps were performed, the syringe containing the support and phosphoramidite solutions was shaken in a thermomixer for about 20 minutes. In the coupling steps, 2 equivalents of the phosphoramidite (Gm$^{iBu}$/Am$^{Bz}$/m6Am$^{Pac}$), 2'-OH phosphoramidite (A$^{Bz}$/m6A$^{Ac}$) or biscyanoethyl phosphoramidite and 0.30 M 5-(benzylthio)-1-H-tetrazole in acetonitrile (except for m6A$^{Ac}$ dissolved in dichloromethane/acetonitrile 2/1) were recirculated through the column for 15 minutes. A solution of 3% (v/v) dichloroacetic acid in toluene was used as a detrytilation reagent and 0.05 M iodine in pyridine for oxidation. After the last cycle of synthesis 20% (v/v) diethylamine in acetonitrile was passed through the column to remove 2-cyanoethyl protecting groups. Finally, the solid support was washed with acetonitrile and dried with argon. The product was cleaved from the solid support and deprotected with AMA (methylamine/ammonium hydroxide 1:1$_{v/v}$; 50 °C, 1 h), evaporated to dryness and re-dissolved in DMSO (200 µl). The TBDMS groups were removed using triethylammonium trihydrofluoride (TEA·3HF; 250 µl, 65 °C, 3 h), and then the mixture was cooled down and diluted with 0.25 M $NaHCO_3$(aq) (20 ml). The product was isolated by ion-exchange chromatography on DEAE Sephadex (gradient elution 0-1.2M TEAB) to afford – after evaporation – the triethylammonium salt of pNmpGmpG and pNpGmpG trinucleotide.

| Sequence | Synthesis scale [µmol] | Yield [µmol] | m/z $_{calcd.}$ | m/z $_{found}$ |
|---|---|---|---|---|
| **pAmpGmpG** | 50 | 29.60 | 1064.18198 | 1064.18216 |
| **p(m6Am)pGmpG** | 25 | 13.30 | 1078.19763 | 1078.19870 |
| **pApGmpG** | 100 | 63.80 | 1050.16633 | 1050.16740 |
| **p(m6A)pGmpG** | 150 | 116.00 | 1064.18198 | 1064.18339 |

Triethylammonium salt of pN$_{m}$pG$_{m}$pG was dissolved in DMSO to 0.05 M and *P*-imidazolide of N7-methylguanosine 5'-diphosphate [m7GDP-Im] (2) (2 eqv) and anhydrous $ZnCl_2$ (20 eqv) were added. The mixture was stirred for ca. 48 h and the reaction was quenched by addition of 10 volumes of aqueous solution of EDTA (20 mg/ml) and $NaHCO_3$ (10 mg/ml). The product was isolated by ion-exchange chromatography on DEAE Sephadex (gradient elution 0-1.2 M TEAB) and purified by semi-preparative RP HPLC (gradient elution 0-15% acetonitrile in 0.05 M ammonium acetate buffer pH 5.9) to afford – after evaporation and repeated freeze-drying from water – ammonium salt of tetranucleotide cap m7GpppNpGmpG.

For cap2-1 derivatives, P-imidazolide activation was performed on trinucleotides. The triethylammonium salt of pNpGmpG was dissolved in DMSO (0.08 M) followed by the addition of imidazole (16 eqv), 2,2'-dithiodipyridine (6 eqv), triethylamine (3 eqv) and triphenylphosphine (6 eqv). The reaction was stirred at RT, until a complete conversion was found by RP-HPLC analysis. The reaction was quenched by the addition of a cold solution of sodium perchlorate (3 eqv) in acetone (10×$V_{reaction}$). The resulting precipitate was washed with acetone, centrifuged until the supernatant was clear and dried under vacuum affording the expected product as a white powder. The resulting P-imidazolide was directly used in a coupling reaction as described for pNmpGmpG with m7GDP (1.5 eqv) instead of m7GDP-Im. The final semi-preparative RP HPLC was performed using a gradient elution from 0 to 50% acetonitrile in 120 min or 60 min for m7GpppApGmpG and m7Gppp(m6A)pGmpG, respectively.

| Sequence | Im-pNpGmpG [µmol] | m7GDP [µmol] | $ZnCl_2$ [mmol] | m7GpppNpGmpG [µmol] | Yield | m/z calcd. | m/z found |
|---|---|---|---|---|---|---|---|
| **m7GpppAmpGmpG** | 8.50 | 17.0 | 170 | 4.74 | 56% | 1503.21139 | 1503.21153 |
| **m7Gppp(m6Am)pGmpG** | 13.30 | 26.5 | 265 | 8.39 | 63% | 1517.22704 | 1517.22838 |
| **m7GpppApGmpG** | 55 | 82.5 | 1.10 | 16 | 29% | 1489.19574 | 1489.19661 |
| **m7Gppp(m6A)pGmpG** | 94.6 | 141.9 | 1.89 | 23.5 | 25% | 1503.21139 | 1503.21179 |

### RNA preparation

*Short RNA*

Short RNAs were obtained as described in (1) with some modifications. Annealed IVTshF and IVTshR oligonucleotides containing T7 promoter sequence and additional 35 nucleotides including recognition site for DNAzyme, served as a template for *in vitro* transcription reaction. Reactions were set in 80 µl mixtures which contained: RNA Pol buffer (40 mM Tris-HCl pH 7.9, 10 mM $MgCl_2$, 1 mM DTT, 2 mM spermidine); 1.25 µM annealed oligonucleotides; ATP, CTP, UTP, 3 mM each and 0.75 mM GTP (R0481, Thermo Fisher Scientific); 2.25 mM cap (analog) of interest; 2 U/µl RiboLock RNase Inhibitor (EO0381, Thermo Fisher Scientific), 4 µl of home-made T7 RNA polymerase, and were incubated at 37 °C; after 2 h additional 4 µl of home-made T7 RNA polymerase were added and reactions were conducted for another 2 h at 37 °C; next, DNase I (EN0521, Thermo Fisher Scientific) was added (0.1 U/µl) for 30 min incubation at 37 °C. Control sample – non-capped short RNA – was prepared in reaction with ATP, CTP, UTP and GTP, 3 mM each, without any cap analog. Transcripts were purified utilizing RNA Clean & Concentrator-25 kit (R1017, Zymo Research). RNAs of intended size were recovered from particular fractions collected during HPLC purification. For purification, Phenomenex Clarity® 3 µm Oligo-RP column was utilized and a linear gradient of buffer B (0.1 M triethylammonium acetate pH 7.0 and 50% acetonitrile) from 10% to 26.7% in buffer A (0.1 M triethylammonium acetate pH 7.0) over 25 min at 1 ml/min was applied. Fractions containing both uncapped and capped transcripts were collected. RNAs after HPLC were recovered by isopropanol precipitation. To generate homogenous 3' end a in these, transcripts were incubated at 1 µM concentration with 1 µM DNAzyme in 50 mM $MgCl_2$ and 50 mM Tris-HCl pH 8.0 at 37 °C for 1 h and were purified utilizing RNA Clean & Concentrator-25 kit and again HPLC-purified, when fractions containing uncapped and capped transcripts, devoid of DNAzyme, were collected. RNAs after isopropanol precipitation and resuspension in water were utilized for enzymatic assays and for examination of capping efficiency, which was analyzed in 15% acrylamide/7 M urea/urea gel.

Crude short transcripts after clean-up were used for preparation of RNAs utilized for affinity purification. Biotinylated pAp (pAp N6-PEG-biot) (3) was ligated to RNAs by T4 RNA ligase 1 (M0204, New England Biolabs). Reactions were set with 10 µM RNAs, 400 µM pAp analogs, 1 mM ATP, 10% (v/v) DMSO, 10% (v/v) ligase in 1x commercial ligase buffer overnight at 16 °C. RNAs after ligation were HPLC-purified, fractions containing transcripts with ligated biotinylated pAp analog were collected. Precipitated and re-suspended RNAs were treated with Calf Intestinal Alkaline Phosphatase (18009027, Thermo Fisher Scientific) to remove any triphosphates from 5' ends of uncapped molecules, which would interact with IFIT proteins later in the experiments. Enzyme was heat-inactivated (15 min at 65 °C).

*Messenger RNA*

mRNAs encoding *Gaussia* luciferase were obtained as previously described in (1) with modifications. Plasmid pJET_T7_Gluc_128A linearized with AarI enzyme (ER1582, Thermo Fisher Scientific) served as a template for *in vitro* transcription reactions. Reactions were set in 20 µl mixtures which contained: RNA Pol buffer (40 mM Tris-HCl pH 7.9, 10 mM $MgCl_2$, 1 mM DTT, 2 mM spermidine); 40 ng/µl of DNA template; ATP, CTP, UTP, 2 mM each and 0.5 mM GTP (R0481, Thermo Fisher Scientific); 1.5 mM cap analog of interest; 2 U/µl RiboLock RNase Inhibitor (EO0381, Thermo Fisher Scientific), 1 µl of home-made T7 RNA polymerase, and were incubated at 37 °C; after 2 h additional 1 µl of home-made T7 RNA polymerase was added and reaction was conducted for another 2 h at 37 °C; next, DNase I (EN0521, Thermo Fisher Scientific) was added (0.1 U/µl) for 30 min incubation at 37 °C. Control sample – non-capped mRNA – was prepared in reaction with ATP, CTP, UTP and GTP, 2 mM each, and no cap analog added. The crude mRNAs were purified using NucleoSpin RNA Clean-up XS (740903, Macherey-Nagel). mRNAs were HPLC-purified using RNASep Prep – RNA Purification Column (ADS Biotec) at 55 °C, a linear gradient of buffer B (0.1 M triethylammonium acetate pH 7.0 and 50% acetonitrile) from 17.5% to 25.8% in buffer A (0.1 M triethylammonium acetate pH 7.0) over 20 min and flow rate was 0.9 ml/min. mRNAs from collected fractions were recovered by isopropanol precipitation. Non-capped mRNAs from samples with capped mRNAs were removed by two separate treatments – with RNA 5'-polyphosphatase (RP8092H, Epicentre) and Xrn1 (M0338, New England Biolabs) –

including purification using NucleoSpin RNA Clean-up XS step in-between. Transcripts after enzymatic reactions were again purified utilizing NucleoSpin RNA Clean-up XS.

**Cell lines and culture conditions**

A549 human carcinoma lung cells (ATCC CCL-185) and 3T3-L1 murine embryonic fibroblasts (ATCC CL-173) were cultured in DMEM medium (Gibco) supplemented with 10% FBS, GlutaMAX (Gibco) and penicillin/streptomycin. THP-1 human monocytes (ATCC TIB-202) and murine immature dendritic cell line JAWS II (ATCC CRL-11904) were grown in RPMI 1640 (Gibco) supplemented with 10% FBS, sodium pyruvate (Gibco), penicillin/streptomycin, and in the case of JAWS cells additionally with 5 ng/ml GM-CSF (PeproTech). Cells were cultured at 37 °C in 5% $CO_2$ atmosphere.

*Stress conditions*

To stress the cells, incubation in medium with addition of Universal Type I Interferon (IFNα) (11200-1, PBL Assay Science) was performed. For RT-qPCR assessment of cellular stress phenotypes upon IFNα treatment A549 cells were seeded in 96-well plates (10 000 cells/well) 24 h before addition of IFNα for 5 h at 0, 50, 500 or 5000 U/ml concentration. Cells for protein production studies were seeded 5 h before mRNA transfection and treated with 0, 50, 500 or 5000 U/µl of IFNα during seeding; after transfection cells were cultured without exchanging medium for additional 72 h. For affinity purification experiment and subsequent western blot analysis, A549 cells were cultured in 100 mm dish to 80-90% confluency and incubated with 0 or 500 U/µl of IFNα for 5 h.

**Reporter mRNA translation in cells – luminescence assay**

For translation studies, 10 000 of A549, JAWS II, 3T3-L1 or THP-1 cells were seeded at the day of experiment in 100 µl medium/well of 96-well plates, with particular concentration of IFNα if applicable. Cells in each well were transfected with mixture containing 5 ng of HPLC-purified mRNA in 5 µl of Opti-MEM (51985026, Gibco) mixed with 0.3 µl Lipofectamine MessengerMAX Transfection Reagent (LMRNA, Invitrogen) in additional 5 µl Opti-MEM. 3 wells for each mRNA sample served as technical replicates. After 72 h incubation the medium was collected for examination of secreted luciferase activity. For detection of *Gaussia* luciferase activity, 50 µl of 10 ng/ml h-coelenterazine (301, NanoLight) in PBS was added to 10 µl of cell culture medium and the luminescence was measured with the use of Synergy H1 (BioTek) microplate reader. Total protein production for each mRNA over 4 days (cumulative luminescence) was reported as a mean value ± SD. Data was collected for 3 biological replicates.

*Silencing CMTR1/2 with siRNA*

Luminescence assay was performed for cells pre-treated with siRNA targeting CMTR1 or 2 or both 1 and 2 as follows. In the first biological replicate, A549 cells were seeded 6000/well in a 96-well plate. The next day, cells were transfected with siRNA sets targeting CMTR1 (set of 3 RNA oligo duplexes, MBS8212469), CMTR2 (set of 3 RNA oligo duplexes, MBS8235611), 1 and 2 (3+3 RNA oligo duplexes), or negative siRNA as a control (MBS8241404), all purchased from MyBioSource. Transfection mixtures contained 5 µl Opti-MEM with 30 nM siRNA, each duplex, and 0.3 µl Lipofectamine RNAiMAX (13778100, Thermo Fisher Scientific) in additional 5 µl of Opti-MEM. Cells were incubated in medium with transfection mixture for 48 h, then medium was exchanged for 100 µl of fresh DMEM. Another transfection was performed, with mRNA molecules capped with various cap structures. 5 ng of HPLC-purified *Gaussia* luciferase mRNA in 5 µl of Opti-MEM with 0.3 µl Lipofectamine MessengerMAX in additional 5 µl of Opti-MEM per well were used. Cells were incubated for 16 h and the medium was collected for luciferase activity examination. 3 wells for each mRNA sample served as technical replicates. The second and third biological replicates were conducted in 48-well plates, 15 000 cells were seeded per well in 200 µl DMEM. The next day transfection was performed for the second biological replicate with siRNA like previously (30 nM siRNA, each duplex), for the third one 30 nM of siRNA in total was used for CMTR1 or CMTR2 samples (10+10+10 nM oligo duplexes) and 30+30 nM for CMTR1+2 samples. Transfection mixtures contained siRNA in 10 µl of Opti-MEM and 0.6 µl Lipofectamine RNAiMAX in additional 10 µl of Opti-MEM. After 48 h incubation medium was exchanged for 200 µl of fresh DMEM and another transfection was performed, with mRNA (10 ng/well in 10 µl of OPTI-MEM and 0.6 µl Lipofectamine MessengerMAX in additional 10 µl of OPTI-MEM). After 16 h medium was collected for analysis of luciferase activity.

**Affinity purification with mass spectrometry and western blot analysis**

*Affinity purification*

For affinity purification/mass spectrometry (AP-MS) analysis, A549 cells were cultured to 90-95% confluency; two 100 mm culture dishes were used for each replicate. All subsequent steps were carried on ice with pre-cooled reagents, if not stated otherwise. PBS washed cells were detached with use of scraper in 800 µl of lysis buffer (20 mM Tris-HCl pH 7.4, 150 mM NaCl, 2 mM $MgCl_2$, 2 mM DTT, 0.2% IGEPAL CA-630, cocktail of protease inhibitors) per dish. Cells and buffer from each dish were collected and transferred to one Eppendorf-type tube, and this mixture was aspirated into the syringe and passed through a 26G needle 7 times. Next, lysates were centrifuged at 4 °C, 10 000 x g, for 10 min. Lysates from both plates were pooled and then aliquoted. Each portion out of 5 was mixed with freshly denatured mRNA (or lysis buffer as a control). During lysate centrifugation, short capped and biotinylated RNAs (10 picomoles of each molecule in 10 µl of water) were heated at 95 °C for 3 min and cooled down on ice to denature them. Then, to each sample, 10 µl of 2x concentrated lysis buffer was added. RNAs with lysates were incubated at 4 °C for 1 h with rotation. Then, samples were mixed with pre-washed streptavidin magnetic beads, 40 µl of 50% slurry used per sample (CMG-227, PerkinElmer), and incubated at 4 °C for 30 min with rotation. Beads pre-wash included 3 washes with lysis buffer (without protease inhibitors) and 2 washes with wash buffer (20 mM Tris-HCl pH 7.4, 150 mM NaCl, 2 mM $MgCl_2$, 2 mM DTT, 0.2% Tween-20). After incubation with lysates, beads were washed 2 times with lysis buffer and two times with wash buffer. Beads after discarding the buffer from the last wash were frozen until next steps for MS analysis were performed. For the experiment with JAWS II, lysate was prepared from approximately $12x10^6$ cells. Suspension fraction of JAWS II was collected by centrifuging medium, while adherent cells were detached with use of trypsin. Cells were pooled, washed thoroughly with PBS and transferred to two Eppendorf-type tubes. Cells from each of two pellets were suspended in 800 µl of lysis buffer and next steps of the procedure were the same as described above.

For affinity purification/western blot analysis, A549 cells were cultured to 90-95% confluency, and the lysate from one 100 mm culture dish was prepared for each condition (-/+IFN). Subsequent steps were carried as described above, but 2 picomoles of each short capped and biotinylated RNA molecules and 8 µl of bead slurry, respectively, were used per sample.

*Mass spectrometry*

To each sample, 20 µl of 100 mM $NH_4HCO_3$ and 2.5 µl of 200 mM TCEP were added, samples were vortexed and placed at horizontal shaker (10 000 rpm) at room temperature for 30 minutes. Subsequently, 2 µl of MMTS were added and samples were shaken for 20 minutes at RT. Trypsin/LysC (V5071, Promega) was solved in 8 M urea in 100 mM $NH_4HCO_3$, to final enzyme concentration 0.02 µg/µl, 50 µl was added to each sample. Samples were incubated with shaking at 37 °C for 4 h, then 300 µl of NH4HCO3 was added and the digestion was performed overnight. Samples were acidified with 10 µl of 5% TFA. The resulting peptide mixture was purified at Oasis HLB 10 mg sorbent (186000128, Waters) 96-well plates, vacuum-dried, and suspended in 60 µl of 2% acetonitrile, 0.1%TFA. Samples were measured in an online LC-MS setup of EvosepOne (Evosep Biosystems) coupled to an Orbitrap Exploris 480 Thermo Fisher Scientific mass spectrometer.
Peptide mixtures were loaded on Evotips C18 trap columns, according to the vendor protocol: activation of sorbent with 0.1% formic acid (FA) in acetonitrile, 2 minute incubation in 1-propanol, chromatographic sorbent equilibration with 0.1% FA in water, samples were loaded in 30 µl of 0.1% FA, after each step, EvoTips were centrifuged at 600 x g for one minute. Chromatographic separation was carried out at a flow rate of 500 nl/min using the 44 min (30 samples per day) performed gradient on an EV1106 analytical column (Dr. Maisch C18 AQ, 1.9 µm beads, 150 µm ID, 15 cm long, Evosep Biosystems, Odense, Denmark). Data were acquired in positive mode with a data-dependent method using the following parameters: the MS1 resolution was set to 60 000 with a normalized AGC target of 300%, auto maximum inject time and a scan range of 350 to 1400 m/z. For MS2, the resolution was set to 15 000 with a standard normalized AGC target, auto maximum inject time, and top 40 precursors within an isolation window of 1.6 m/z considered for MS/MS analysis. Dynamic exclusion was set at 20 s with an allowed mass tolerance of ±10 ppm, with a precursor intensity threshold of 5 × 1 000. Precursors were fragmented in HCD mode with a normalized collision energy of 30%. The spray voltage was set to 2.1 kV, with a funnel RF level of 40 and heated capillary temperature of 275 °C.
Raw data were analyzed with PEAKS Studio 10.6 64bit Bioinfor (4) and searched against Uniprot human (78 120 entries, for the A549 samples) or mouse (55 360 entries for the JAWS II samples) reference proteomes. Fixed modifications: methylthio (MMTS) at cysteines; variable: oxidation methionine, acetyl n-term. MS error and 0.1 Da, MS/MS level 0.2Da, FDR 1%, digestion: trypsin semi specific, max variable PTM per peptide: 3. Protein level analysis was performed using the “Label Free” PEAKS module.
Each group consisted of 3 biological replicates, average signal intensity was calculated for every condition. Data were analyzed in a way to indicate which proteins co-precipitate with the immobilized decoy in a repeatable and specific manner compared to the given control group.

Ratio of any protein identified and quantified was calculated in relation to the average level in the MOCK samples normalized to “1.0”. Heat-maps, clustering and group correlation was done in Perseus software (5). Ratio values were represented as log2 at heat-maps, rows and columns were clustered based on Euclidean distance calculation.

Data were deposited in the PRIDE under two accessions:
**Project accession:** PXD028636
**Project DOI:** 10.6019/PXD028636
Reviewer account details:
**Username:**
**Password:** P18yjuHx

**Project accession:** PXD028635
**Project DOI:** 10.6019/PXD028635
Reviewer account details:
**Username:**
**Password:** qvESgcrb

*Western blot analysis*

Beads after protein affinity purification and input samples (cell lysates) were mixed with Laemmli sample buffer (63 mM Tris-Cl (pH 6.8), 10% glycerol, 2% SDS and 0.01% bromophenol blue, 5% 2-mercaptoethanol) and heated for 5 min at 95 °C. Samples were separated in 10% SDS-PAGE. Then, proteins were transferred on to the nitrocellulose membrane (GE10600002, GE Healthcare) using the Trans-Blot Turbo Transfer System (Bio-Rad). Proteins on the membrane were stained with Ponceau S buffer (0.5% Ponceasu S, 1% acetic acid). 1 h blocking in 5% skim milk in PBST was performed. Membranes were cut horizontally into 3 pieces and each was incubated with appropriate primary antibody diluted 1 000-fold in PBST to detect eIF4E (2067, Cell Signaling), IFIT1 (PA3-848, Thermo Fisher Scientific) and NCBP1 (PA5-83607, Thermo Fisher Scientific) overnight at 4 °C. After washing with PBST, membranes were incubated with secondary goat anti-rabbit (HRP) antibody (32260, Thermo Fisher Scientific) diluted 10 000-fold in PBST for 1 h at room temperature. Detection was performed with the use of Immobilon Western Chemiluminescent HRP Substrate (WBKLS0, Merck) in Amersham Imager 600.

**eIF4E affinity assay – microscale thermophoresis (MST)**

To determine dissociation constant for eIF4E protein and capped short RNA complexes, microscale thermophoresis-based method was applied, previously described in (6), with modification of the utilized ligand form – here, capped short RNAs were analyzed. Binding assay mixtures were prepared in 20 µl as follows: 10 nM fluorescent probe m7Gp5OC3(5)FAM, 50 nM murine eIF4E and ligand – differently capped short RNAs ranging from 1.75 µM to 0.05 nM, in an MST buffer (50 mM Hepes–KOH pH 7.2, 100 mM KCl, 0.5 mM EDTA, 0.05% Tween-20). Samples after preparation, without any additional incubation, were loaded into Monolith NT.115 Capillaries (MO-KO22; NanoTemper Technologies). MST was performed using a Monolith NT.115 instrument (NanoTemper Technologies) at 25 °C. Instrument parameters were adjusted to 100% LED Blue power and Medium MST power. To determine the $K_{D,\ app}$ values, a standard 1:1 binding model was fitted to the data using PALMIST software (version 1.4.4). Confidence Intervals were determined using error-surface projection (ESP) (7).

**Recombinant hDCP2 and hDXO cloning, overproduction and purification**

*hDXO ORF cloning*

cDNA obtained from total RNA isolated from 293 Flp-In T-REx cells (R78007, Thermo Fisher Scientific) was used as a template to amplify ORF coding for the full-length human DXO WT in PCR with primer pair hDXO1For-hDXO1Rev (0.2 µM both) and using Phusion High-Fidelity DNA Polymerase (F530, Thermo Fisher Scientific) with 1x HF buffer, 0.2 mM dNTP Mix and 3% DMSO. PCR product corresponding to hDXO wild-type ORF was gel-purified using Gel-Out kit (023-50, A&A Biotechnology) and inserted by SLIC into *Bam*HI/*Xho*I sites of pET28M N-6xHis-TEV vector (8), giving rise to phDXOwt construct. *E. coli* MH1 strain (*E. coli araD lacX74 galU hsdR hsdM rpsL*) was used for transformation with SLIC products. An insert encompassing ORF encoding hDXO mut (E234A D236A) was obtained in a two-step amplification. In the first step, two PCR reactions were performed as above, but using hDXOfor-hDXOmutR and hDXOmutF-hDXOrev primer pairs and phDXOwt as a template. The two PCR products were gel-purified as above and their mixture was used in the second round of amplification. Initially, 10 cycles of PCR product joining in the absence of primers was performed, followed by addition of hDXOfor-hDXOrev primer pair and normal PCR (as for hDXO wt). Finally, the resulting PCR product corresponding to hDXO mut ORF was purified and cloned into pET28M N-6xHis-TEV vector as described above, giving rise to the phDXOmut construct.

*hDCP2 cloning*

The 1-350 amino acid coding region of hDCP2(E147Q, E148Q) was PCR amplified from the pET28a-hDCP2(E147Q, E148Q) plasmid [a gift from Megerditch Kiledjian (Addgene plasmid #72215) (9)] with primer pair hDCP2F and hDCP2R and using Phusion High-Fidelity DNA Polymerase with 1x HF buffer and 0.2 mM dNTP Mix. PCR product was purified using NucleoSpin Gel and PCR Clean-up (740609, Macherey-Nagel) and cloned into pJET1.2/blunt vector using CloneJET PCR Cloning Kit (K1231, Thermo Fisher Scientific). Chemocompetent *E.coli* TOP10 bacteria were transformed with ligation mixture. Selected plasmid containing 1-350 amino acid coding region of hDCP2(E147Q, E148Q) of interest was cut with BamHI and NotI and the insert containing hDCP2 fragment was ligated utilizing the same sites into pET28b vector giving rise to phDCP2(1-350)mut. phDCP2(1-350)mut was obtained as a counterpart for plasmid encoding for wild type 1-350 amino acid region of hDCP2 protein, herein named as phDCP2(1-350)wt, kindly gifted by Megerditch Kiledjian.

*Recombinant protein production and purification*

*E. coli* BL21-CodonPlus(DE3)-RIL strain (Agilent; *E. coli* B $F^-$ *ompT hsdS*[$r_B^-$ $m_B^-$] *dcm*$^+$ Tet$^r$ *gal λ[DE3] endA* Hte [*argU ileY leuW* Cam$^r$]) was transformed with either phDXOwt, phDXOmut, phDCP2(1-350)wt or phDCP2(1-350)mut. Transformants were grown in a standard Luria-Broth (LB) medium supplemented with 50 µg/ml kanamycin and 34 µg/ml chloramphenicol. Subsequently, 1 liter of Auto Induction Medium (AIM) Super Broth Base including Trace elements (AIMSB02, Formedium) containing 2% glycerol and both antibiotics, was inoculated with 30 ml of the starter culture. Bacteria were grown for 48 h at 18°C with shaking (150 rpm) and eventually collected by centrifugation at 4 500 rpm in a Sorvall H6000A/HBB6 swinging-bucket rotor for 15 min at 4°C.

Bacterial pellet was resuspended in 70 ml of lysis buffer (50 mM Tris-HCl pH 8.0, 200 mM NaCl, 10 mM imidazole, 10 mM 2-mercaptoethanol, 1 mM phenylmethylsulfonyl fluoride (PMSF), 0.02 µM pepstatinA, 0.02 µg/ml chymostatin, 0.006 µM leupeptin, 20 µM benzamidine hydrochloride), incubated with lysozyme (50 µg/ml; Roth) for 30 min in a cold cabinet, and then broken in an EmulsiFlex-C3 High Pressure homogenizer at 1 500 Bar. The homogenate was centrifuged in a Sorvall WX Ultra Series ultracentrifuge (F37L rotor) at 33 000 rpm for 45 min at 4°C.

The extract (supernatant after high-speed ultracentrifugation) was used for protein purification using the ÄKTA Xpress system (GE Healthcare), employing nickel affinity chromatography on an ÄKTA-compatible 5 ml column that was manually filled with Ni-NTA Superflow resin (Qiagen). The column was equilibrated with 25 ml of low-salt (LS) buffer (50 mM Tris-HCl pH 7.4, 200 mM NaCl, 10 mM imidazole, 10 mM 2-mercaptoethanol) prior to extract loading. After protein binding, the resin was sequentially washed with 40 ml of LS buffer, 25 ml of high-salt (HS) buffer (50 mM Tris-HCl pH 7.4, 1 M NaCl, 10 mM imidazole 10 mM 2-mercaptoethanol), and again 20 ml of LS buffer. Bound proteins were recovered by elution with 30 ml of buffer E (50 mM Tris-HCl pH 7.4, 200 mM NaCl, 300 mM imidazole). Pooled eluate fractions (approximately 5 ml) were dialyzed overnight at 4ºC against 2 liters of LS buffer in the presence of 100/50 µg of home-made TEV/SUMO protease (hDXO wt and mut/hDCP2 wt and mut). This mixture was afterwards subjected to second round of purification on the nickel resin, performed using ÄKTA Purifier system (GE Healthcare) and employing LS buffer for collection of the flow-through, containing protein of interest devoid of the tag, and buffer E2 (50 mM Tris-HCl pH 8.0, 1 M NaCl, 300 mM imidazole) for elution of 6xHis-tagged SUMO or TEV protease and cleaved-off 6xHis-SUMOTag or 6xHis-TEV sequence. For hDXO variants, further purification from contaminating chaperones and degradation products was achieved by separation of pooled flow-through obtained in the second-round of affinity chromatography on size exclusion Superdex 75 10/300 GL column (GE Healthcare) using 1.2 column volumes of gel-filtration (GF) buffer (20 mM Tris-HCl pH 8.0, 150 mM NaCl). Two fractions corresponding to the maximum of $A_{280}$ nm absorbance were collected after gel-filtration and pooled together. Solutions containing purified recombinant were mixed with glycerol (30% v/v), aliquoted, snap-frozen in liquid nitrogen and stored at -80ºC until needed for biochemical activity assays. Proteins were inspected in 10% SDS-PAGE stained with Coomassie Brilliant Blue R-250. PageRuler Prestained Protein Ladder, 10 to 180 kDa (26616, Thermo Fisher Scientific) was used as a molecular weight marker during electrophoresis.

**RNA decapping assays**

*hDCP2*

Purified short uncapped pppRNA (25-nt) and differently capped RNAs with their uncapped fractions (27-nt + 25-nt, respectively) were utilized as substrates in DCP2 enzymatic assay. Reactions were set in 25 µl with 1.6 U/ µl of

RiboLock in reaction buffer (50 mM Tris-HCl pH 8.0, 50 mM $NH_4Cl$, 0.01% NP-40, 5 mM $MgCl_2$, 2 mM $MnCl_2$, 1 mM DTT, where both DTT and $MnCl_2$ were always added shortly before setting the reaction) at 37 °C. 70 nM DCP2 (wild type or mutant) concentration was tested, substrates were utilized at 120 nM concentration. Reactions without enzyme served as controls. During assays, 5 µl of reaction mixtures were collected at each time point (0, 5, 15, 30, 60 min), and reactions were stopped by mixing with formamide loading dye (for PAA electrophoresis) containing 20 mM EDTA and snap freezing in liquid nitrogen. Samples were separated in denaturing 15% urea TBE PAA gels. After staining with SYBR Gold Nucleic Acid Gel Stain (S11494,Thermo Fisher Scientific), gels were scanned in Typhoon FLA9500 Imager (GE Healthcare).

*DXO*

Purified short uncapped pppRNA (25-nt) and differently capped RNAs with its uncapped fractions (27-nt + 25-nt, respectively) were utilized as substrates in DXO enzymatic assay. Reactions were set in 25 µl in reaction buffer (10 mM Tris-HCl pH 7.9, 50 mM KOAc, 2 mM $Mg(OAc)_2$, 2 mM $MnCl_2$, 1 mM DTT, where both DTT and $MnCl_2$ were always added shortly before setting the reaction) at 37 °C. 0.6 µM and 6 µM DXO (wild type or mutant) concentrations were tested, substrates were utilized at 70 nM concentration. Reactions without enzyme served as controls. Assays and samples analysis were conducted as those with DXO protein, described above.

**RT-qPCR**

Cells were incubated for 5 h with various concentrations of IFNα and then lysed utilizing SingleShot Cell Lysis Kit (1725080, Bio-Rad). Lysates were used for reverse transcription (RT) reactions with oligo($dT_{20}$) primer and M-MLV Reverse Transcriptase (28025, Invitrogen). cDNA generated in RT served as templates for quantitative PCR (qPCR) reactions performed utilizing SsoAdvanced Universal SYBR Green Supermix (1725271, Bio-Rad) with primer pairs for amplifying *IFIT1, IFIT2, IFIT3, RIG-I, MDA5, PKR, OAS*, and *GAPDH* – the latter was employed as a reference gene.

**Supplementary Figures and Tables**

**A**

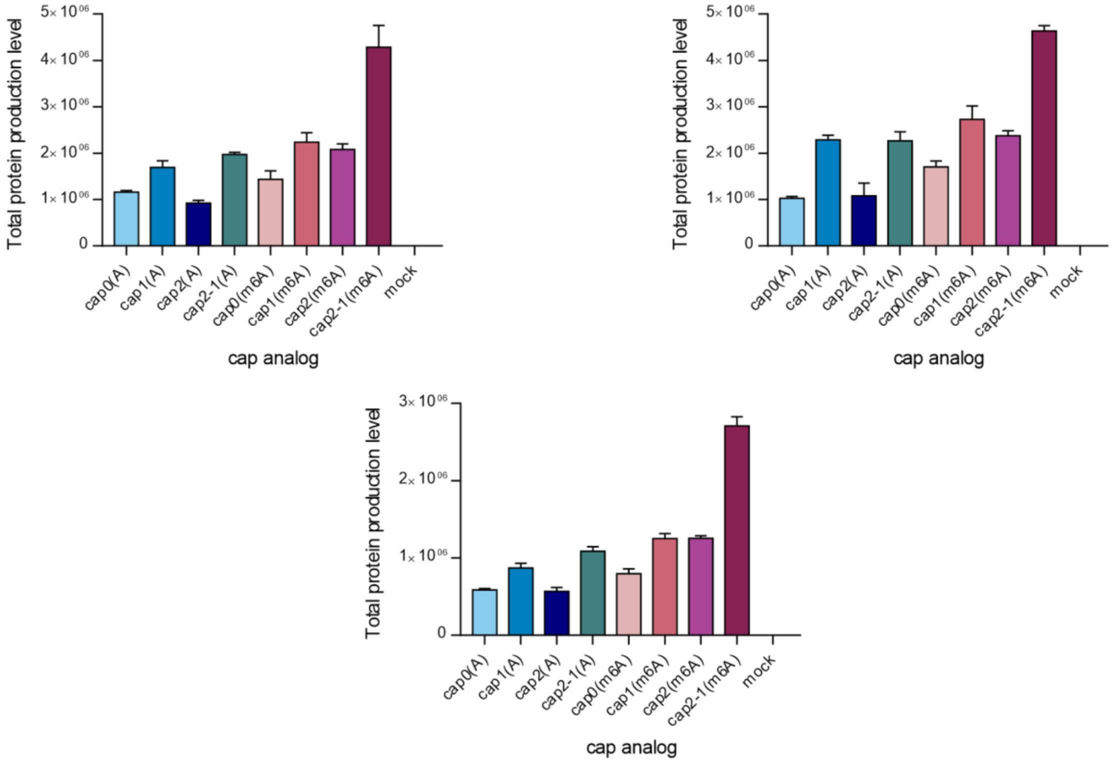


**B**

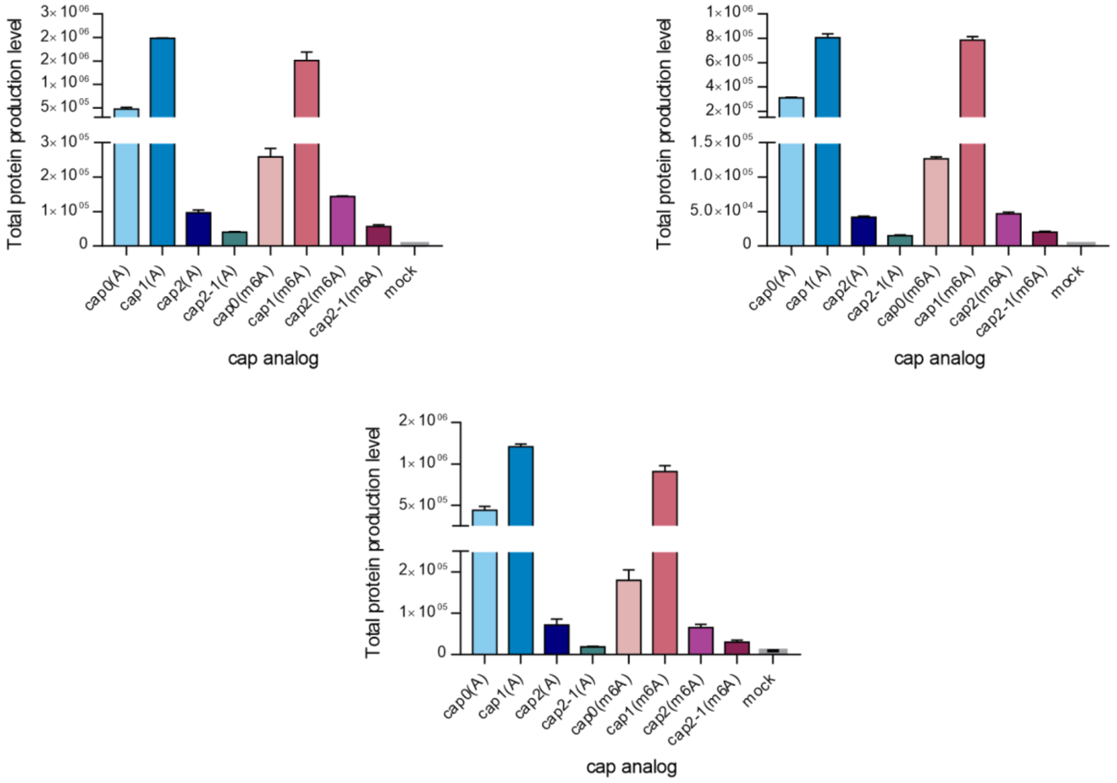


**Fig. S1** Total protein production levels after 72 h measured in the medium from culture of **(A)** A549 and **(B)** JAWS II cells transfected with IVT mRNAs encoding *Gaussia* luciferase bearing various cap analogs at their 5' ends. Three independent biological replicates are presented, each consists of three independent transfections. Bars represent mean value ± SEM.

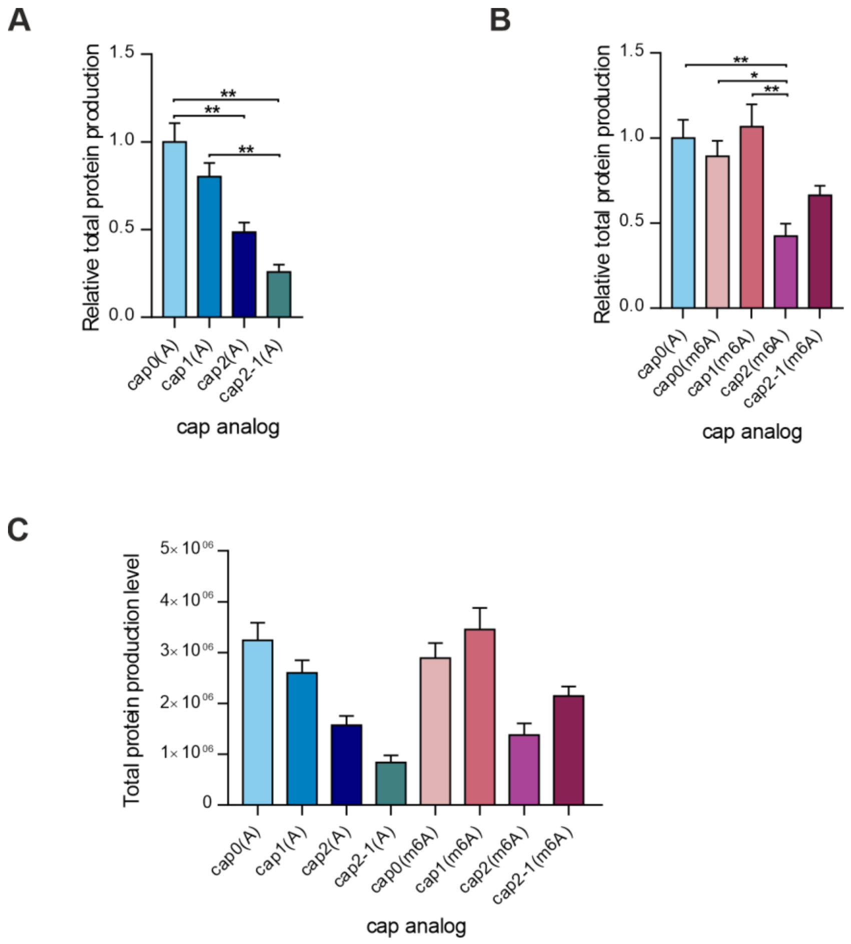


**Fig. S2** Protein production levels after 72 h measured in the medium from culture of THP-1 cells transfected with IVT mRNAs encoding *Gaussia* luciferase bearing various cap analogs at their 5' ends. **(A,B)** Bars represent mean value ± SEM normalized to transcripts with cap0(A). Statistical significance: * P < 0.05, ** P < 0.01, (one-way ANOVA with Turkey's multiple comparisons test). **(C)** Total protein production level used to calculate values on **(A,B)**

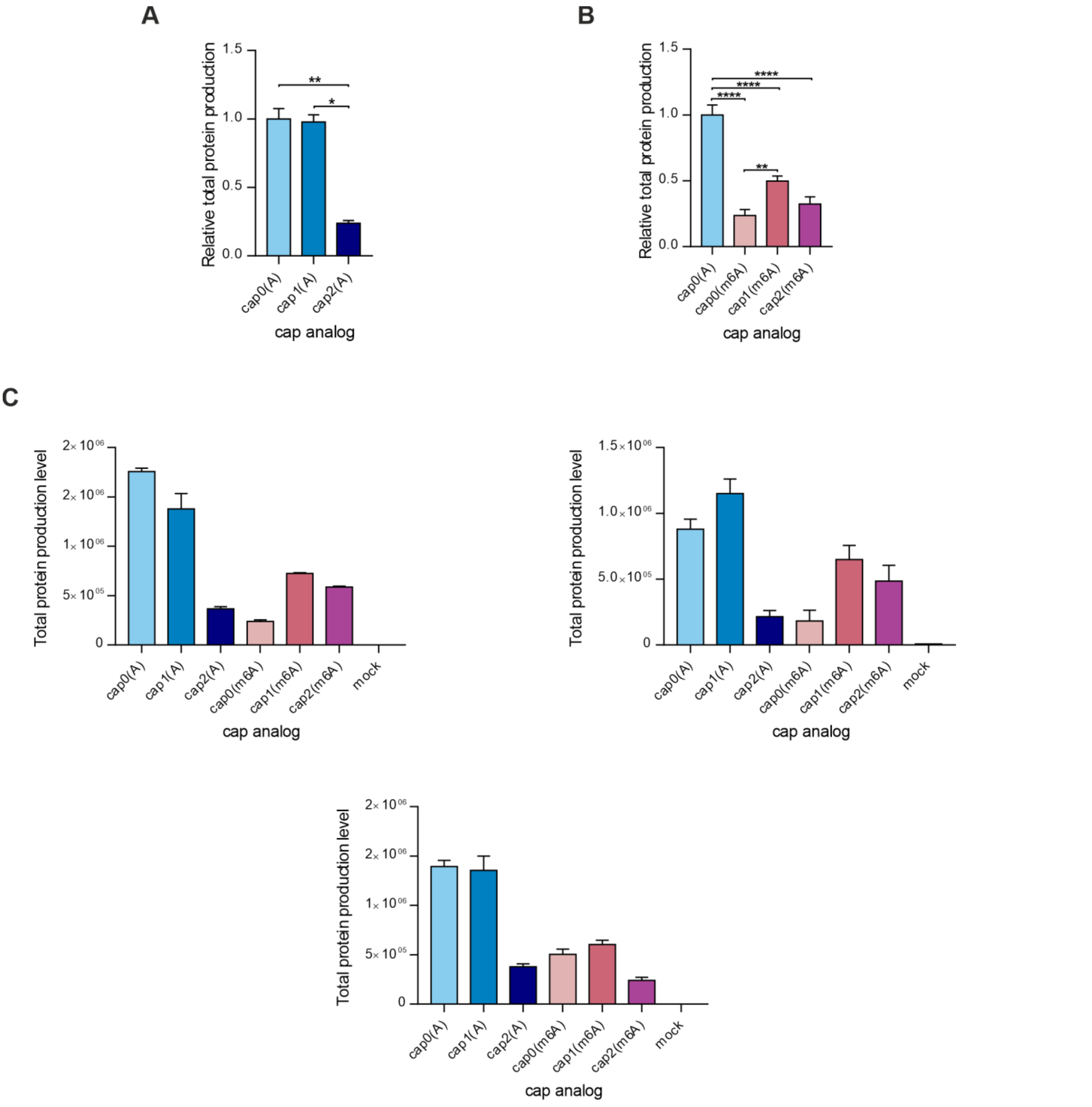


**Fig. S3** Protein production levels after 72 h measured in the medium from culture of 3T3-L1 cells transfected with IVT mRNAs encoding *Gaussia* luciferase bearing various cap analogs at their 5' ends. **(A,B)** Bars represent mean value ± SEM normalized to transcripts with cap0(A). Statistical significance: NS – not significant, * P < 0.05, ** P < 0.01, *** P < 0.001, **** P < 0.0001 (one-way ANOVA with Turkey's multiple comparisons test) **(C)** Three independent biological replicates are presented, each consists of three independent transfections. Bars represent mean value ± SEM

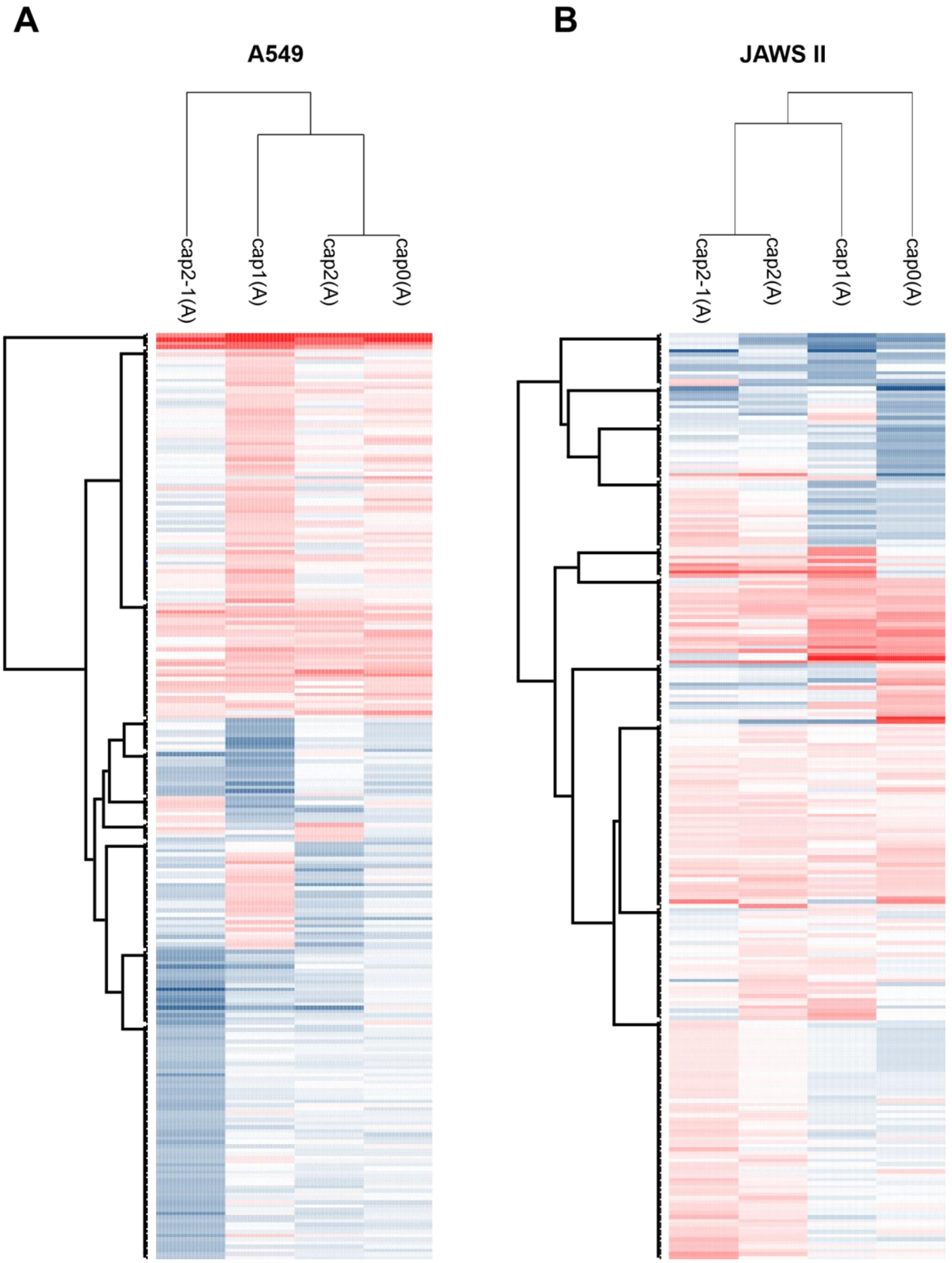


**Fig. S4** Heatmap showing hierarchical clustering of proteins performed on log2 ratio values using Euclidean distances generated in Perseus software (5). Ratio of any protein identified and quantified was calculated in comparison to the average level in the mock samples normalized to “1.0”.

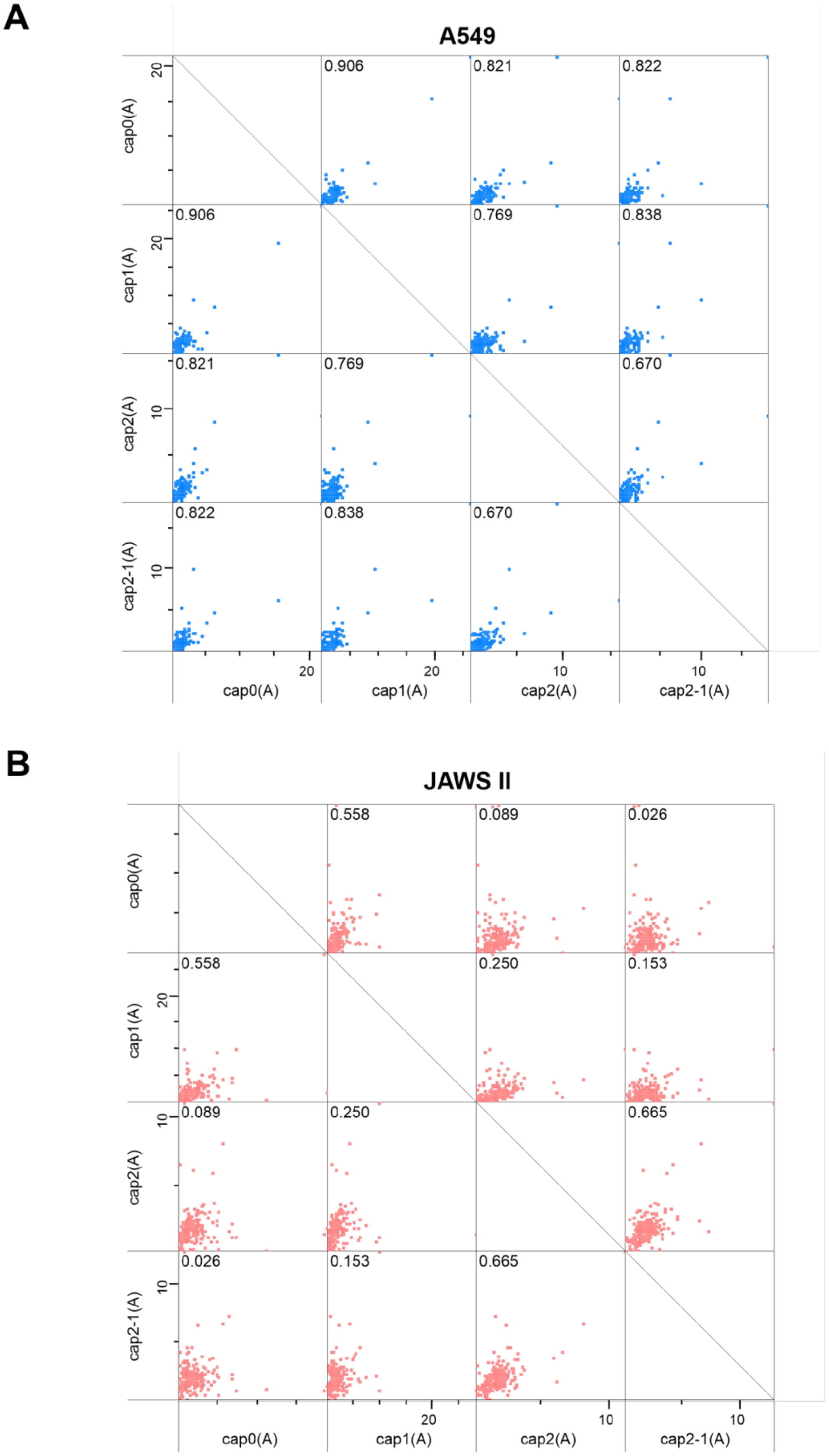


**Fig. S5** Correlation between identified interactomes for differently capped RNA for **(A)** A549 and **(B)** JAWS II cells. Pearson correlation coefficient for each comparison is presented in the left graph upper corner.

**cap0(A)-RNA**

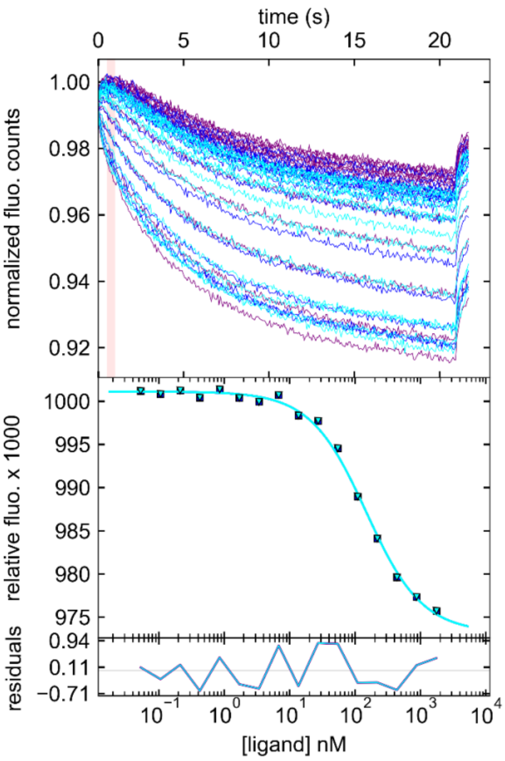


**cap1(A)-RNA**

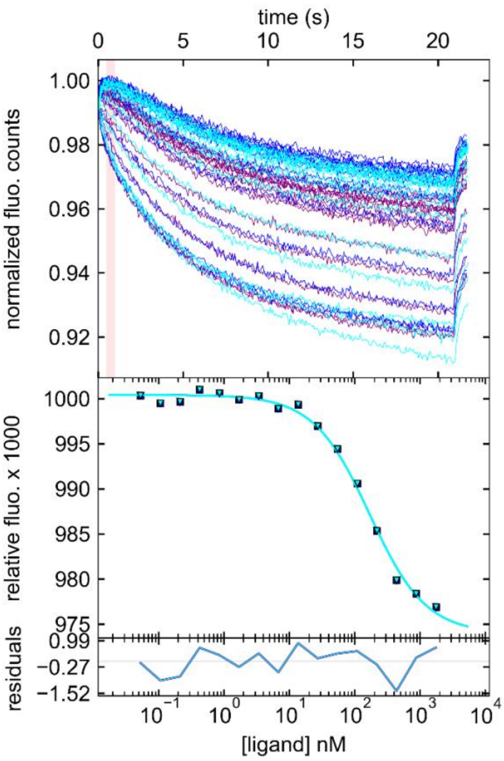


**cap2(A)-RNA**

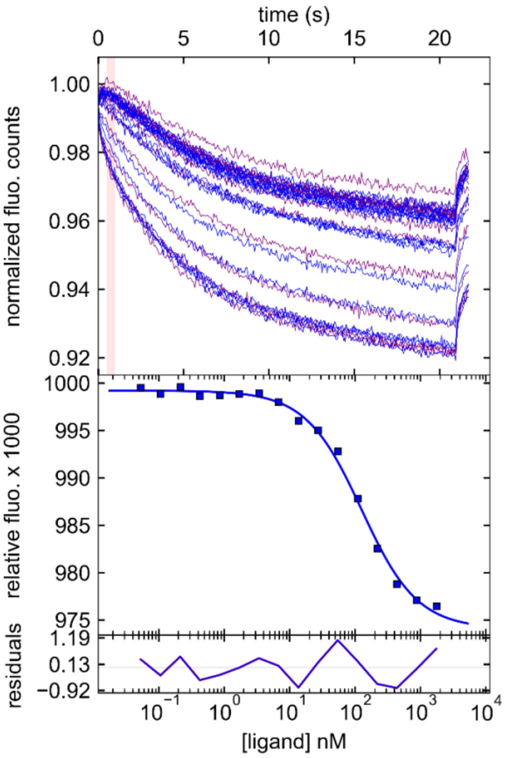


**cap2-1(A)-RNA**

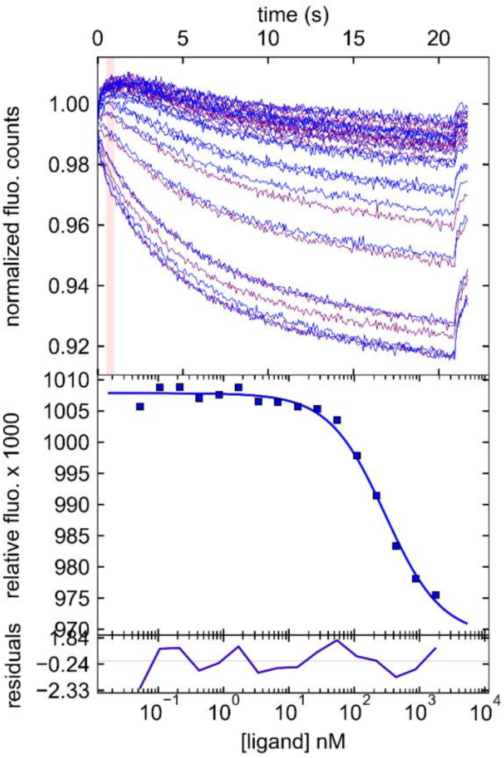

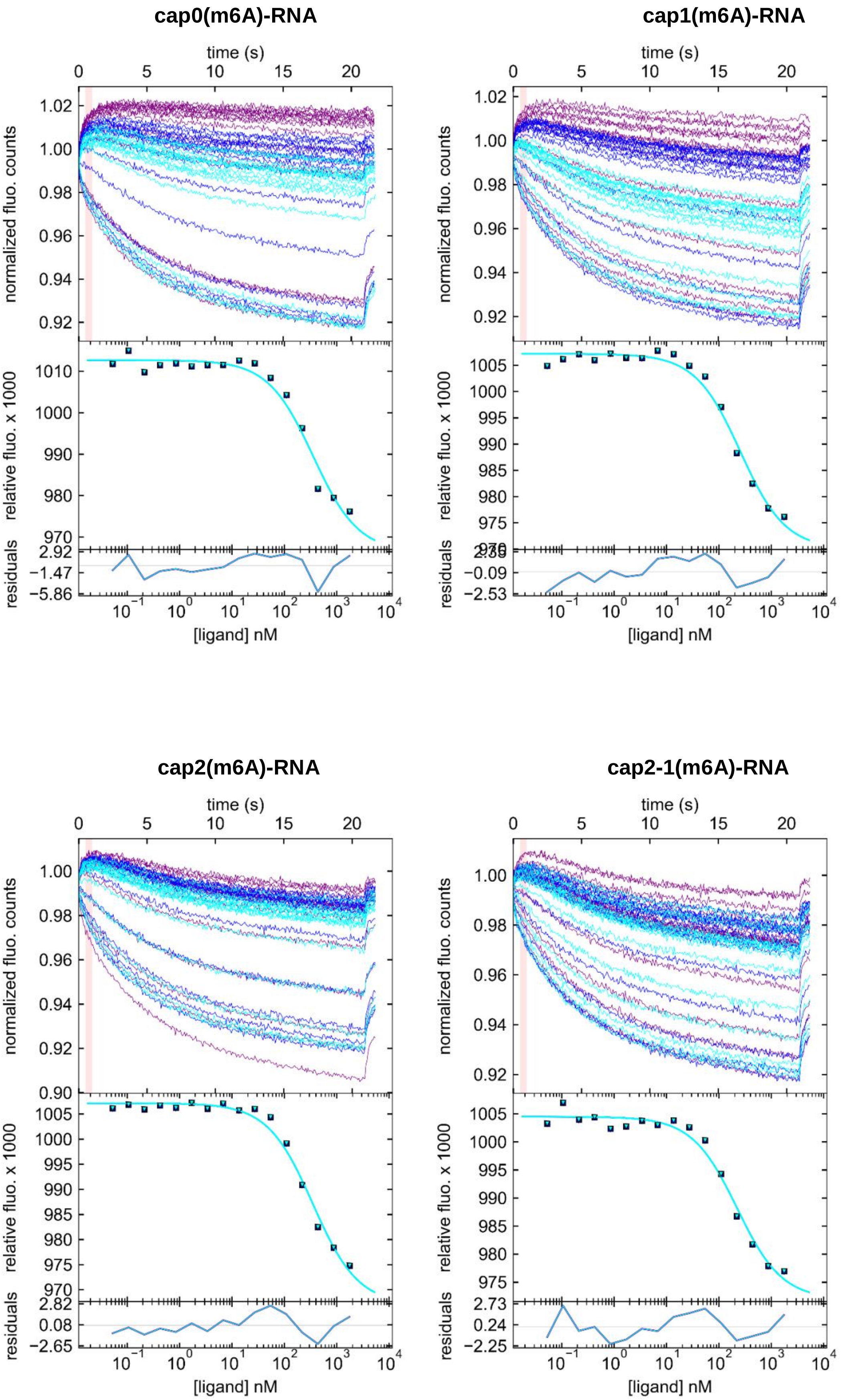
cap0(m6A)-RNA
cap1(m6A)-RNA
cap2(m6A)-RNA
cap2-1(m6A)-RNA
time (s)
normalized fluo. counts
relative fluo. x 1000
residuals
[ligand] nM

### ppp-RNA

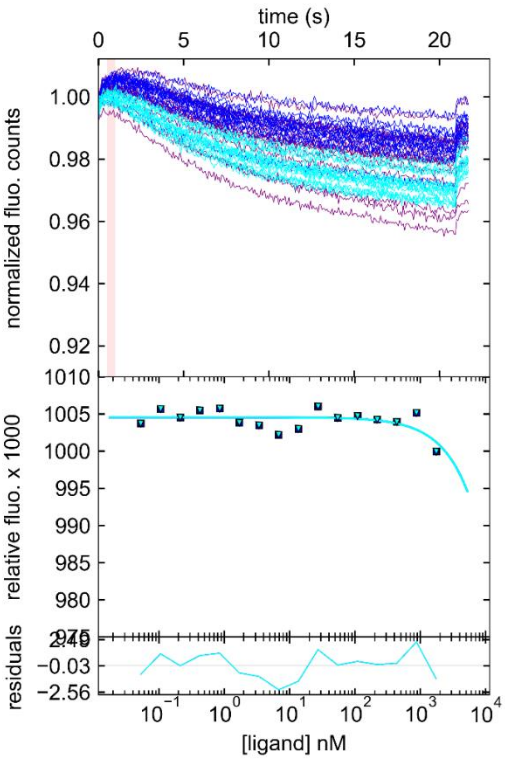


**Fig. S6** Determination of murine eIF4E-capped RNA complexes dissociation constants by microscale thermophoresis (MST) competition assay. Representative MST curves and competitive binding curves obtained for each RNA in the experiment are presented.

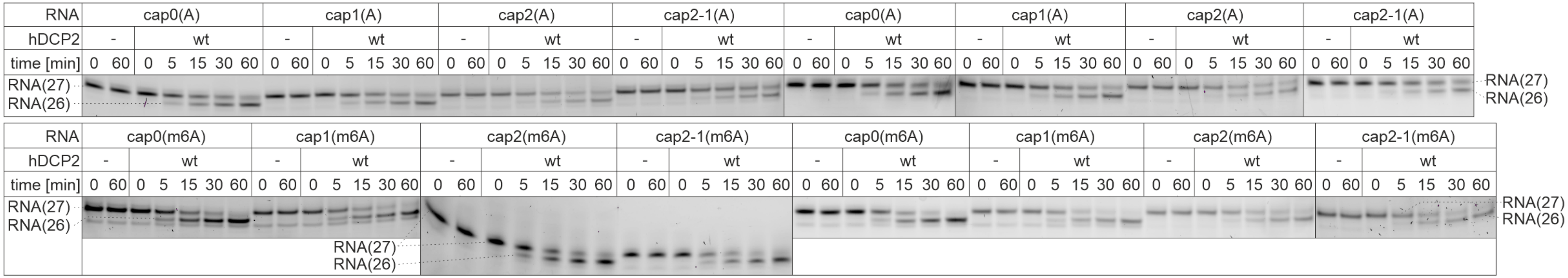


**Fig. S7** Susceptibility of short transcripts to recombinant hDCP2 action. Short capped RNAs were subjected to treatment with wild type hDCP2 for 60 min. Reactions without enzyme served as controls. Aliquots from indicated time points were resolved in polyacrylamide gel and bands corresponding to capped (27-nt long) and to RNAs decapped by hDCP2 action (26-nt long). Two independent replicates are presented. One more replicate of this experiment is shown in Fig. 5A.

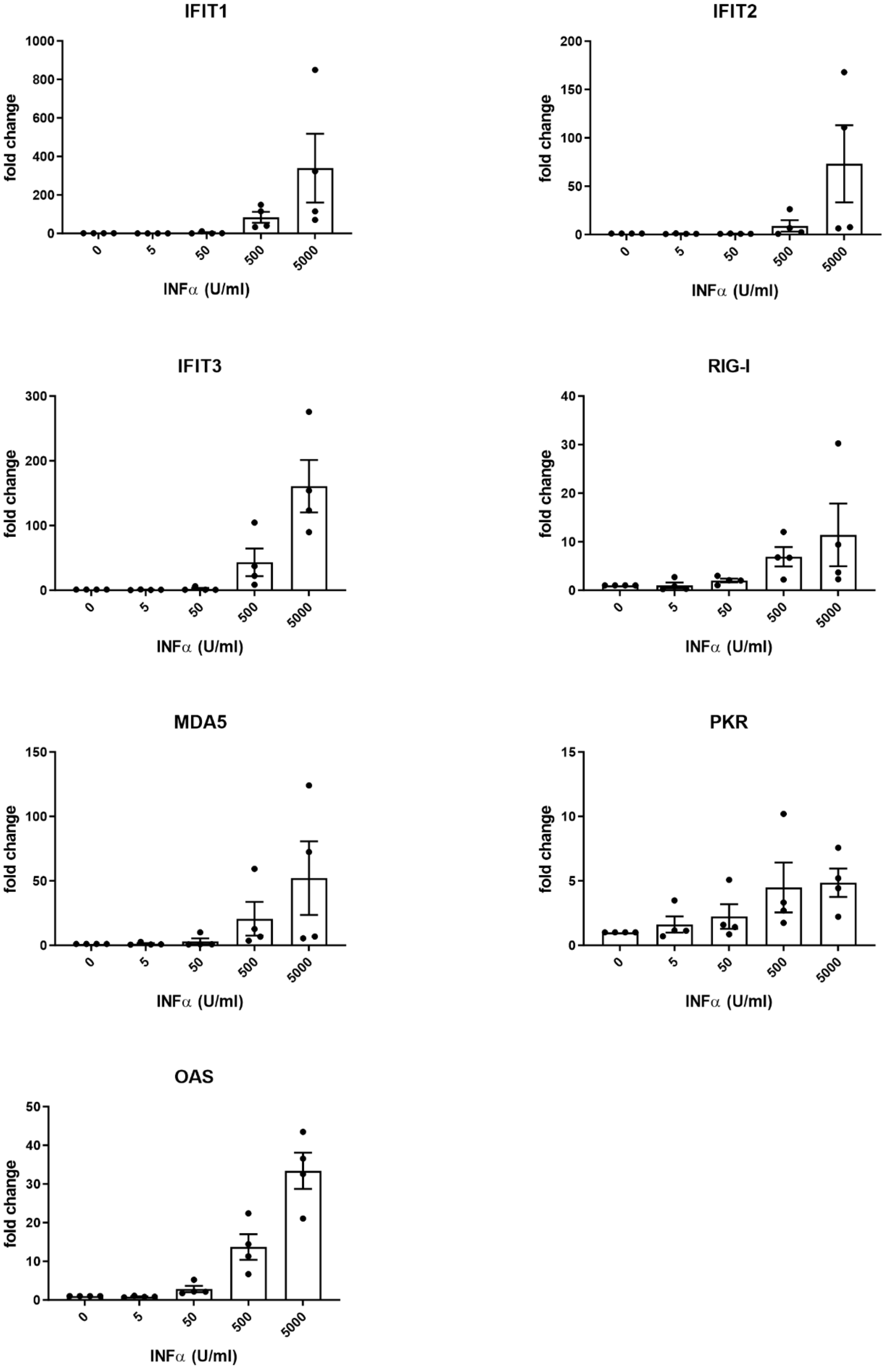


**Fig. S8** Changes in gene expression upon treatment with increasing concentration of IFNα for 5 h. mRNA expression analysis for the indicated genes was carried out using RT-qPCR. Bars represent the mean value of mRNA level change (fold change) ± SEM from four independent biological replicates, each independent biological replicate consists of a single transfection reaction. Each point represents data from one independent biological replicate.

**A**

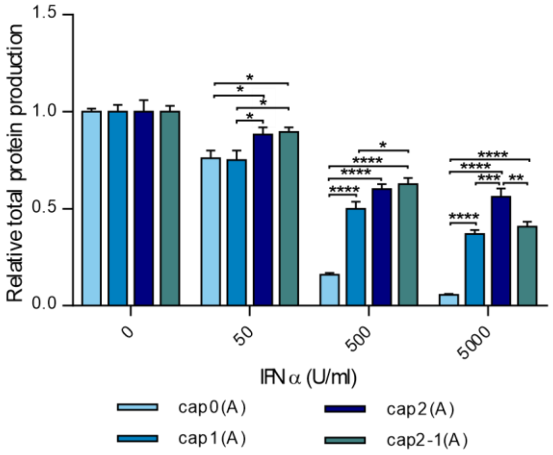


**B**

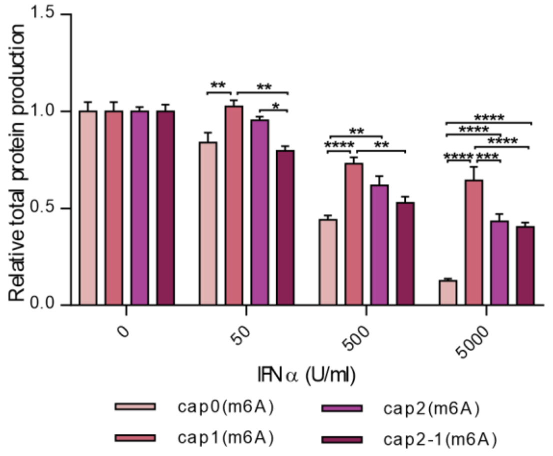


**Fig. S9 (A,B)** Changes in relative protein production in response to increasing concentration of IFNα in A549 cells. Relative protein production levels over 24 hours after 5 hour IFNα pre-treatment in A549 cells. Bars for each transcript represent mean value ± SEM normalized to protein production in mock treated cells. Statistical significance: * P < 0.05, ** P < 0.01, *** P < 0.001, **** P < 0.0001 (one-way ANOVA with Turkey's multiple comparisons test)

**Table S1** HPLC profiles and HRMS spectra of synthetized compounds

| | **pAmpGmpG** |
|---|---|
| **Chemical structure** | 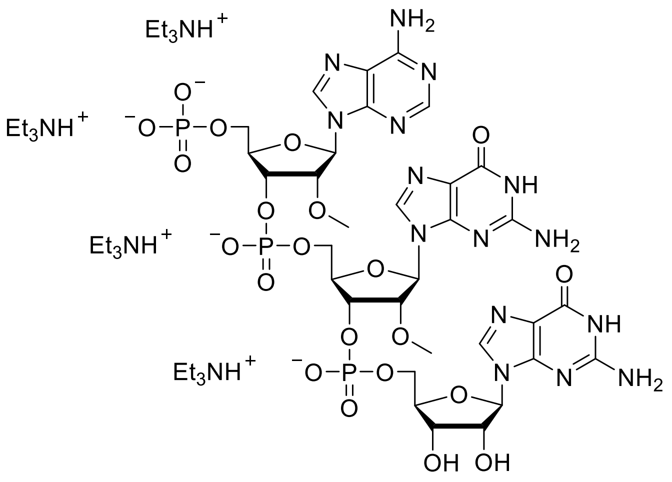 |
| **RP HPLC** | 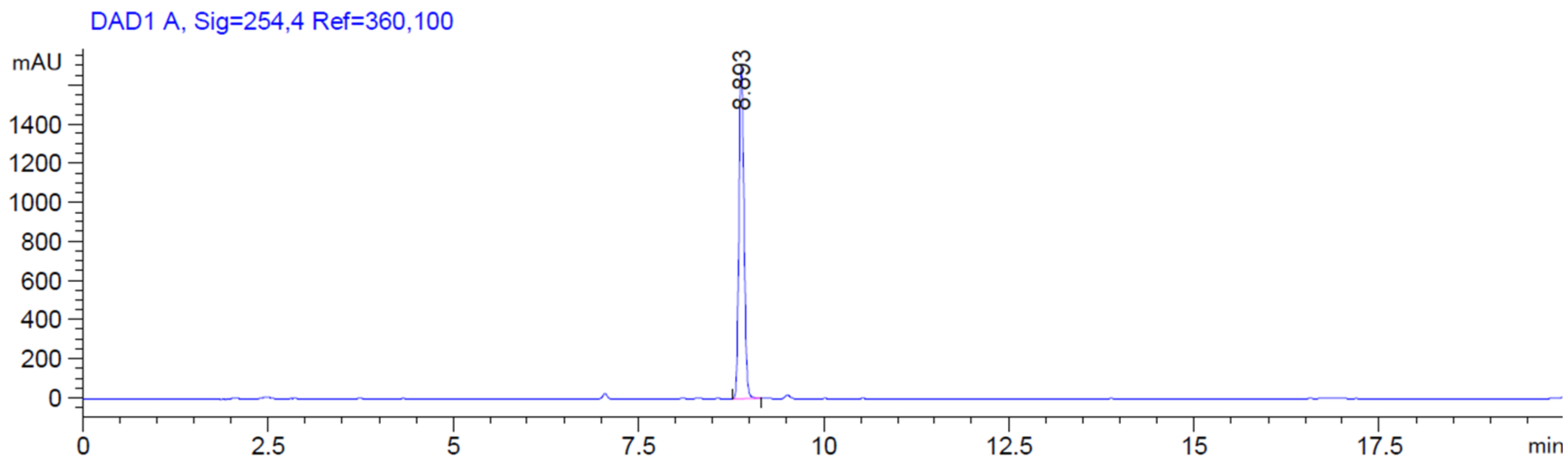  |

| | |
|---|---|
| **HR ESI (-) MS Calc. [M-H]$^-$ $C_{32}H_{41}N_{15}O_{21}P_3^-$ 1064.18198)** | 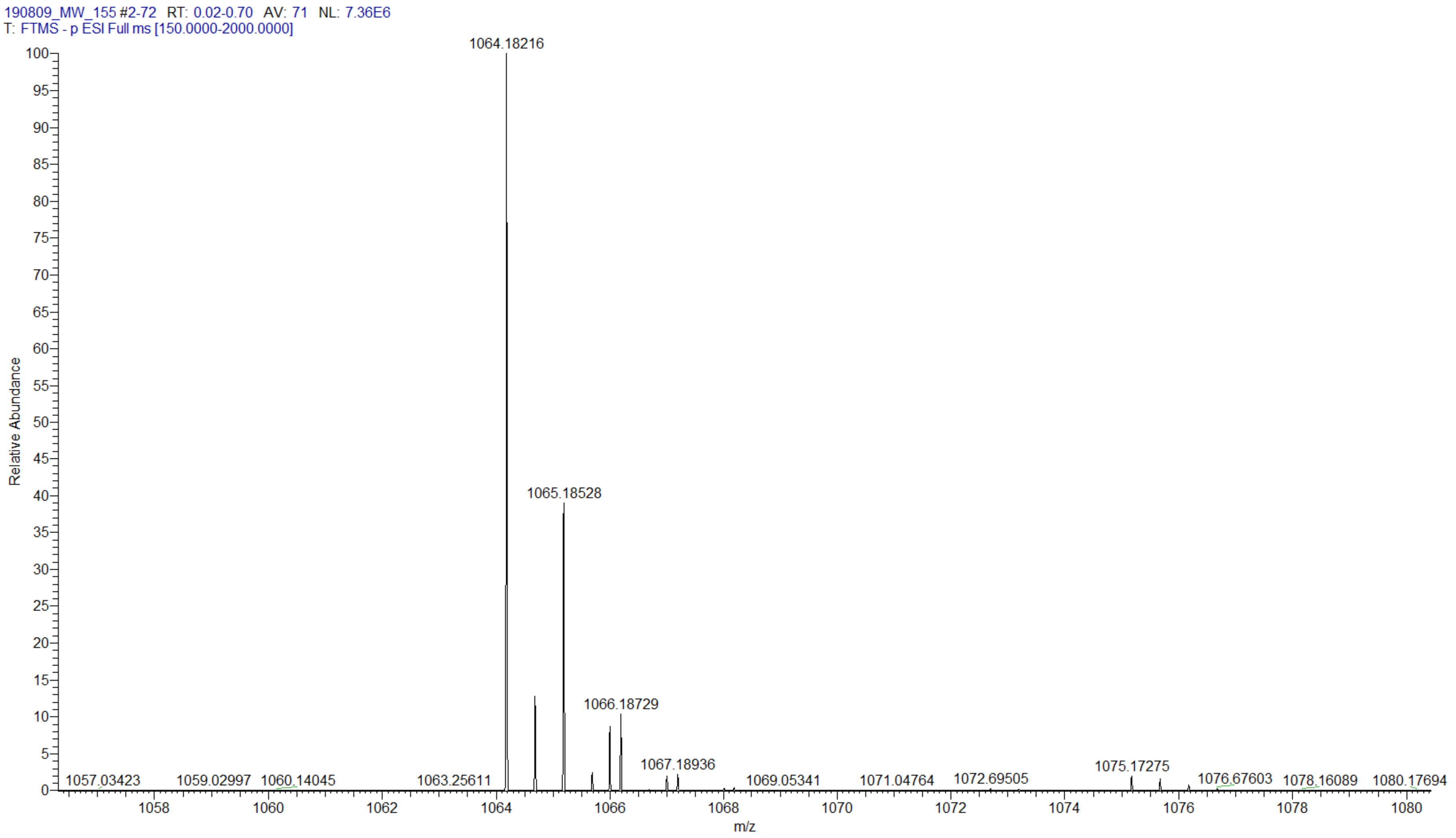 |

| p(m6Am)pGmpG | |
|---|---|
| **Chemical structure** | 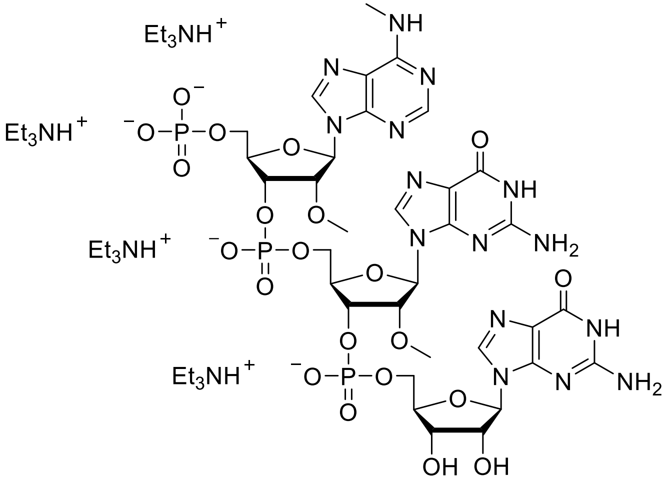 |
| **RP HPLC** | 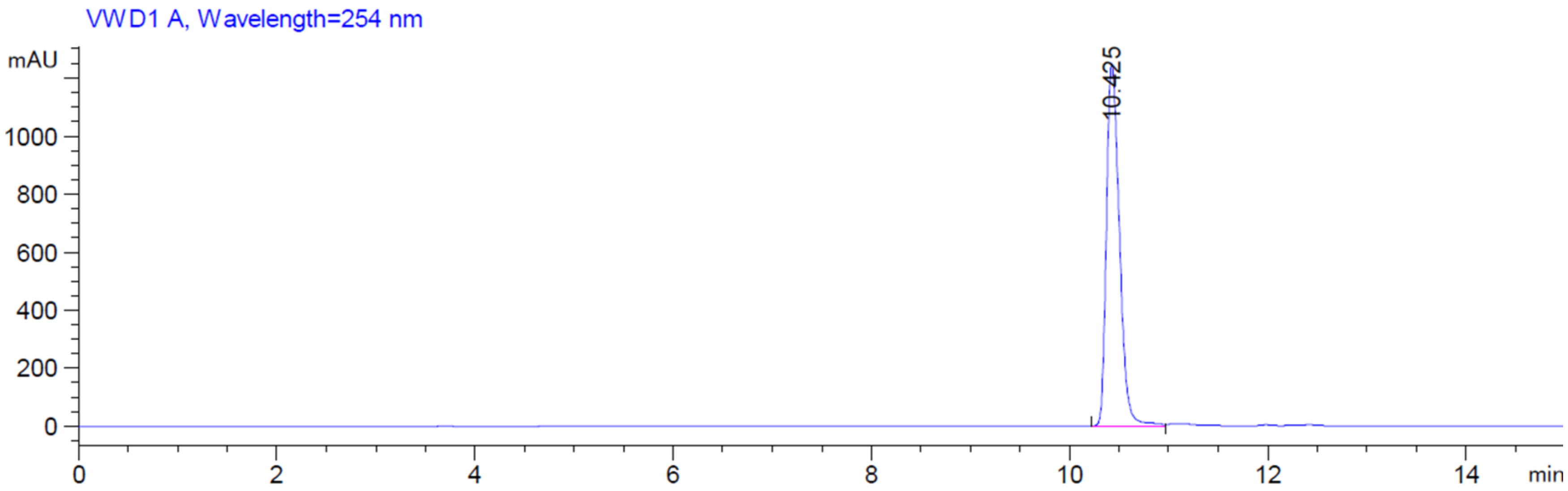 |

**HR ESI(-) MS**

**Calc. [M-H]⁻ $C_{33}H_{43}N_{15}O_{21}P_3^-$ 1078.19763)**

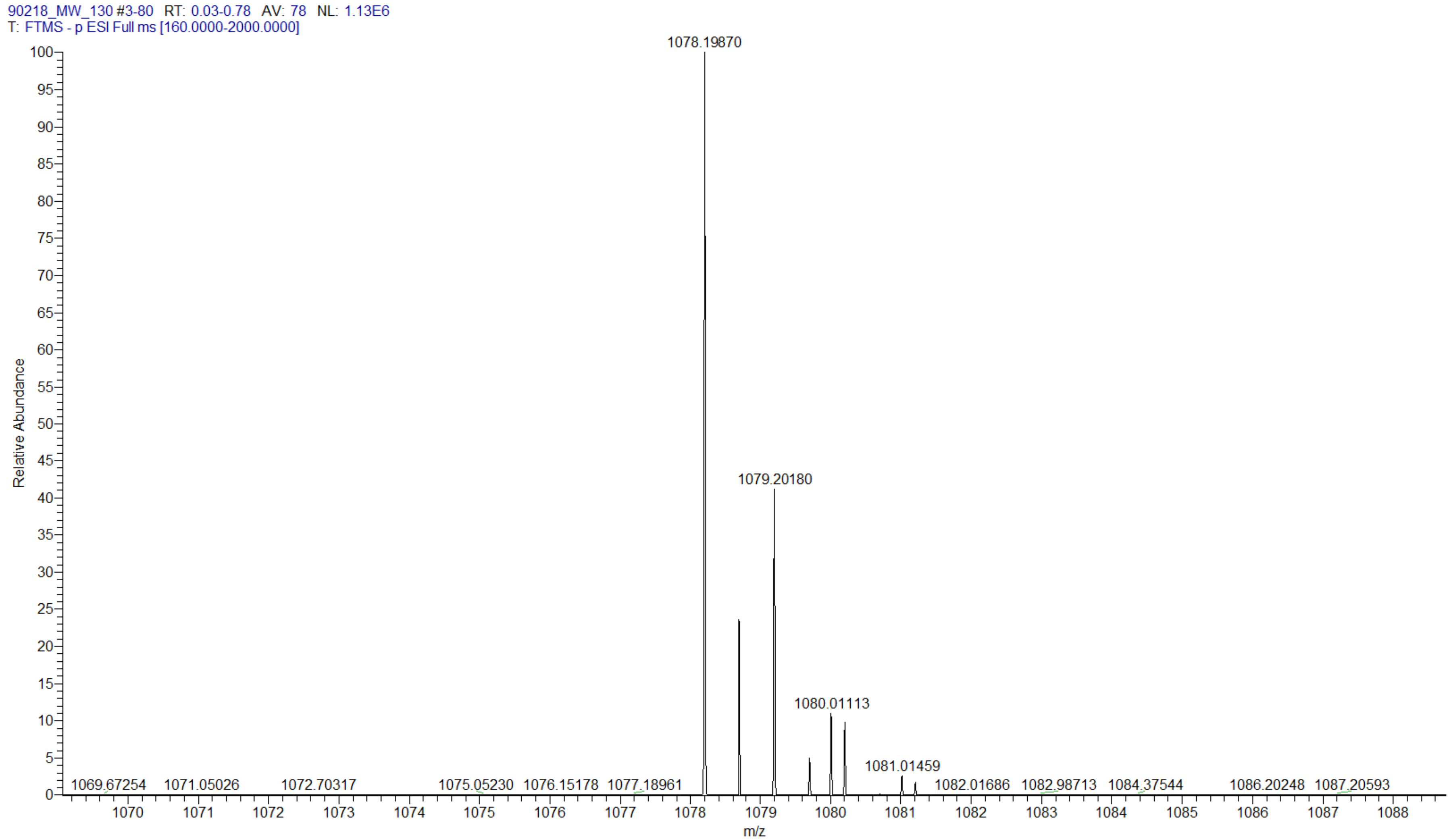


**pApGmpG**

| | |
|---|---|
| **Ch em ica l str uct ur e** | 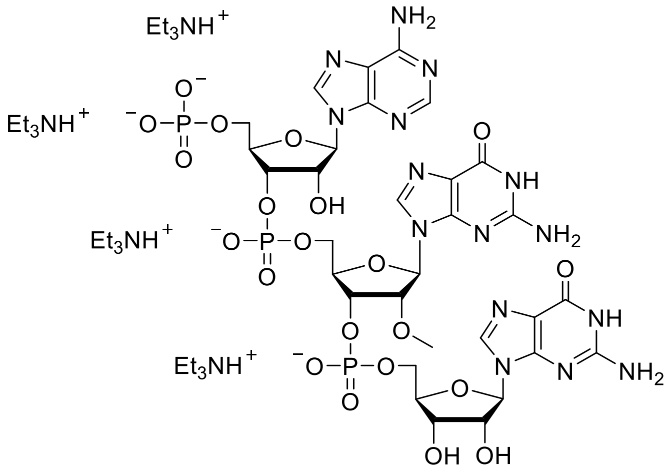 |
| **RP HP LC** | 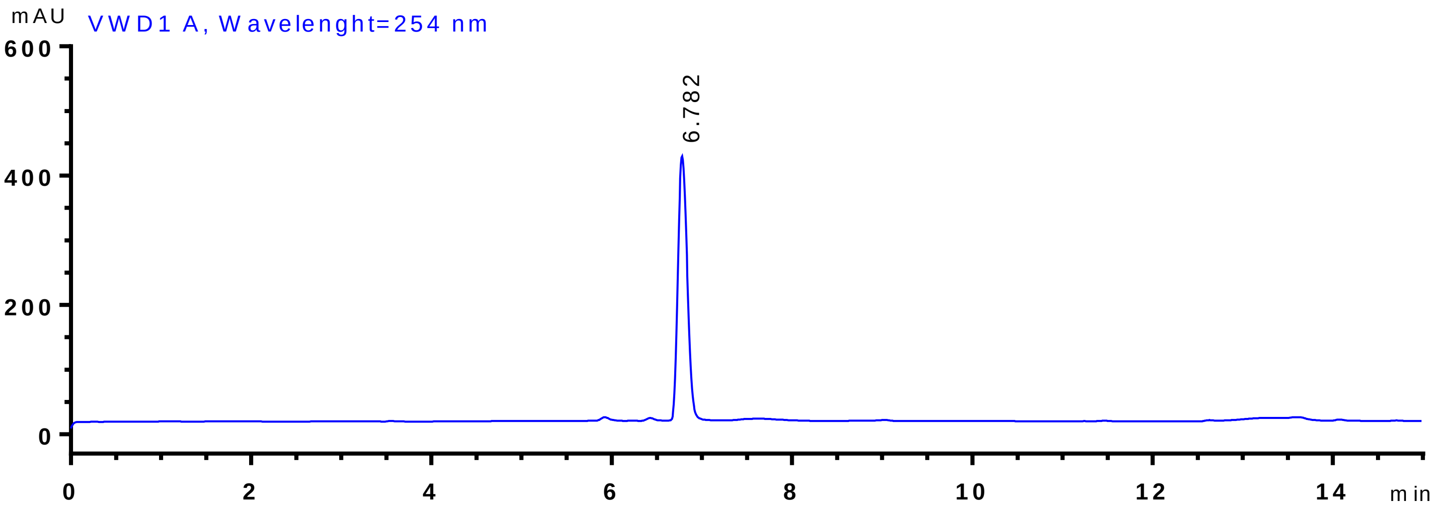 |

**HRESI(-)MS Calc. [M-H]$^{-}$ $C_{31}H_{39}N_{15}O_{21}P_3^{-}$ 1050.16633)**

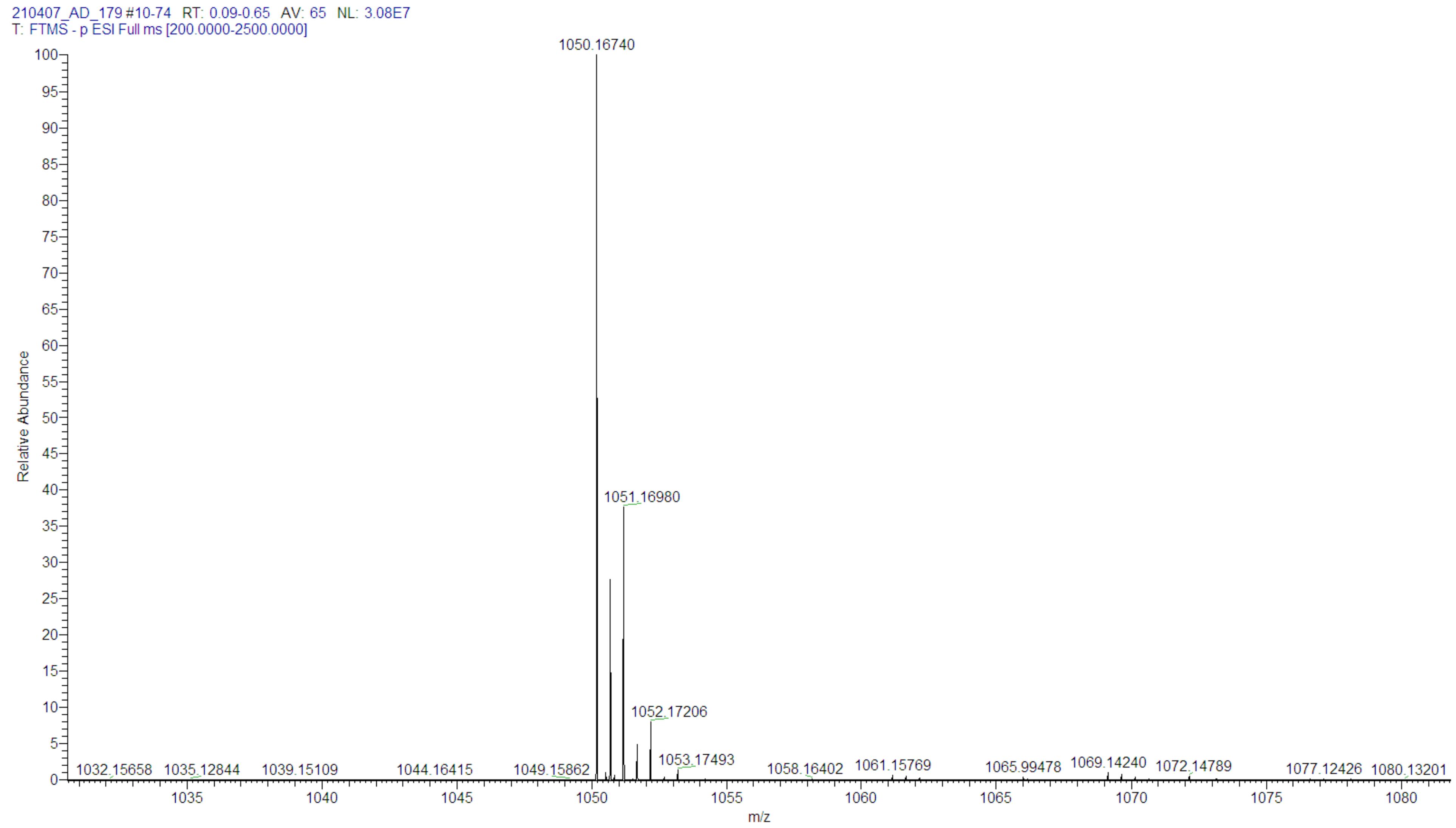

| p(m6A)pGmpG | |
|---|---|
| Chemical structure | 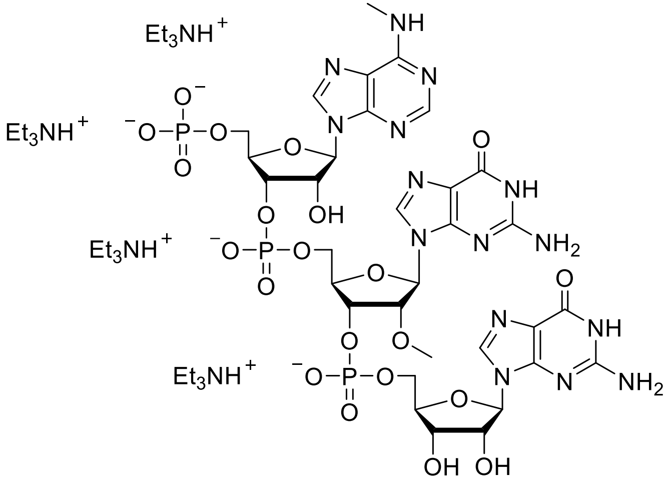 |
| RP HPLC | 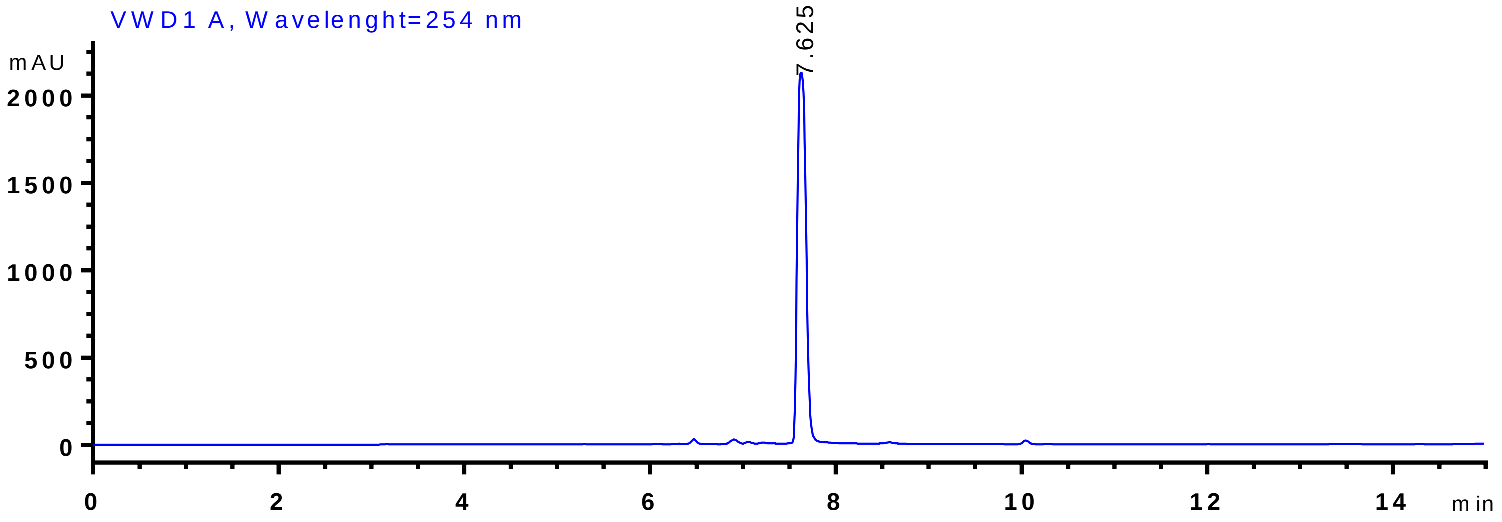 |

| | |
|---|---|
| **HRESI(-)MS Calc. [M-H]$^-$ $C_{32}H_{41}N_{15}O_{21}P_3^-$ 1064.18198)** | 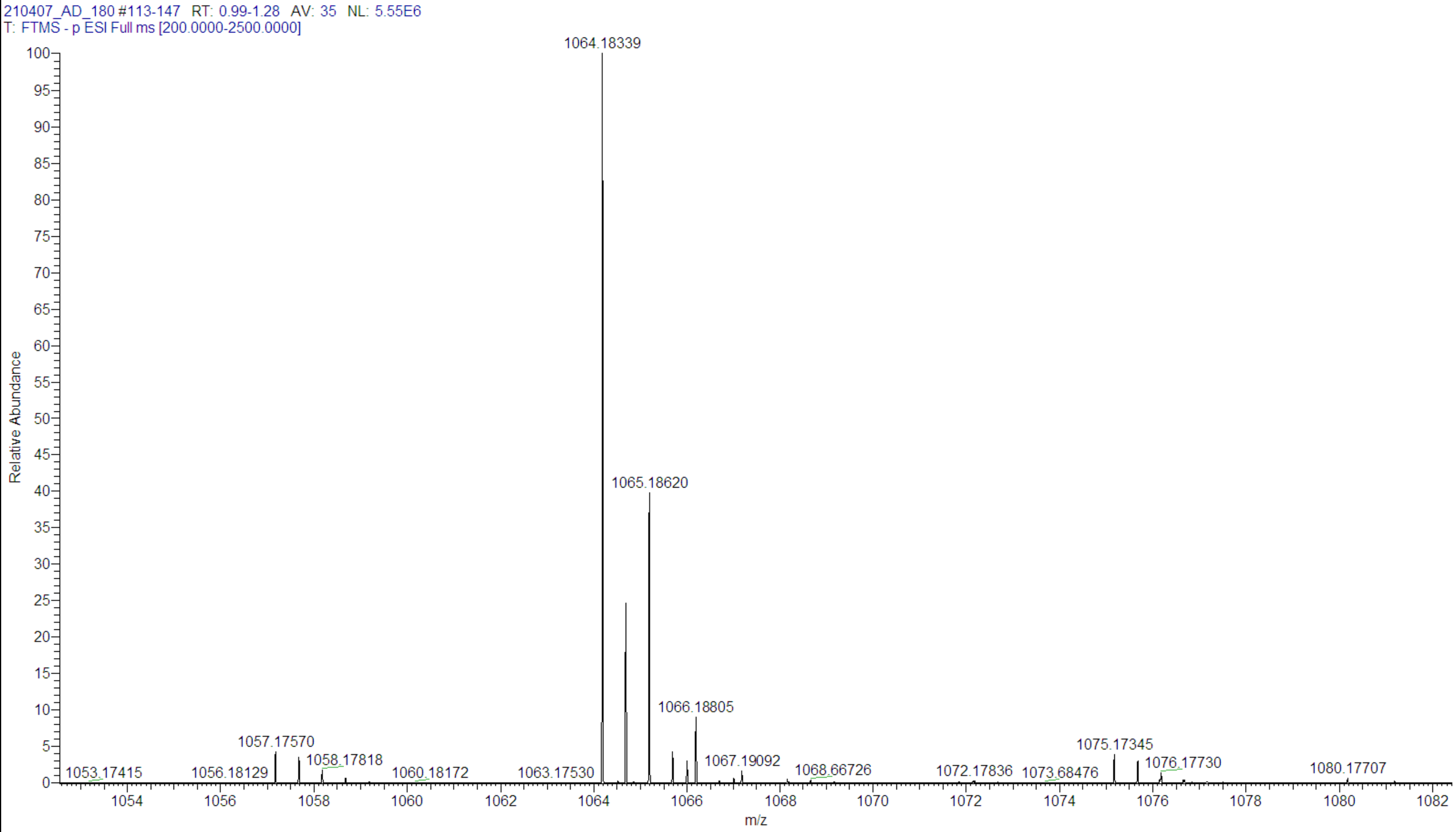 |

| m7GpppAmpGmpG | |
| --- | --- |
| Chemical structure | 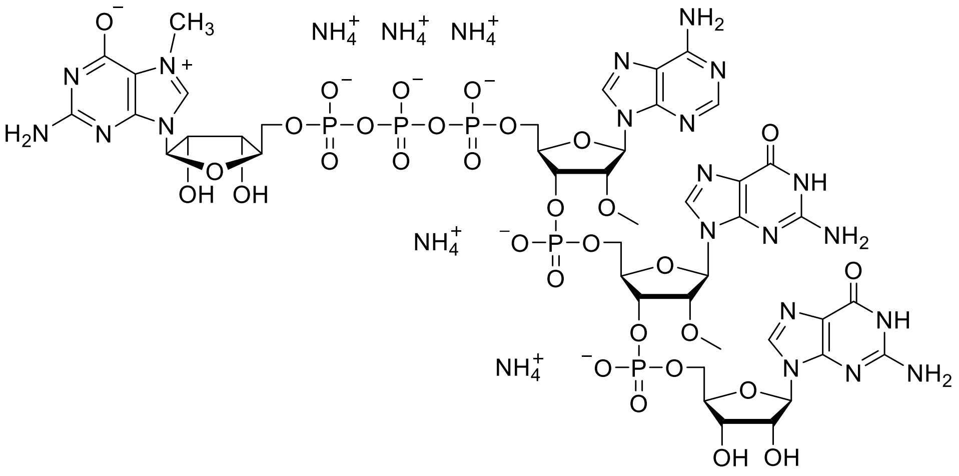 |
| RP HPLC | 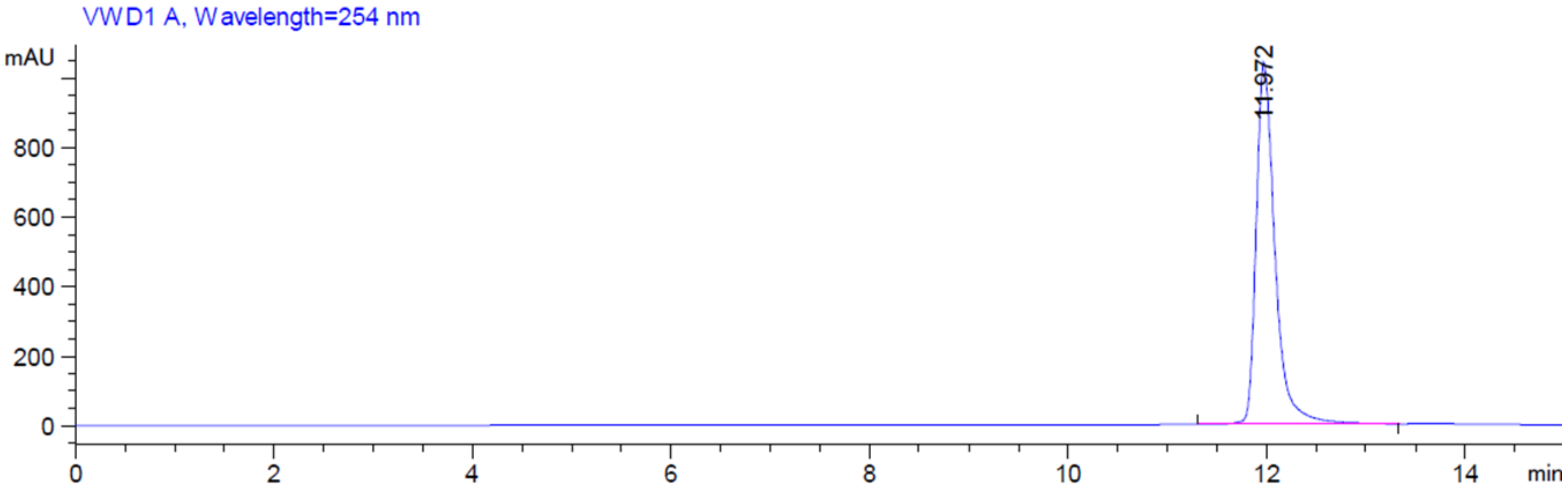  |

| | |
|---|---|
| **HR ESI (-) MS Calc. [M-H]⁻ $C_{43}H_{56}N_{20}O_{31}P_5^-$ 1503.21139)** | 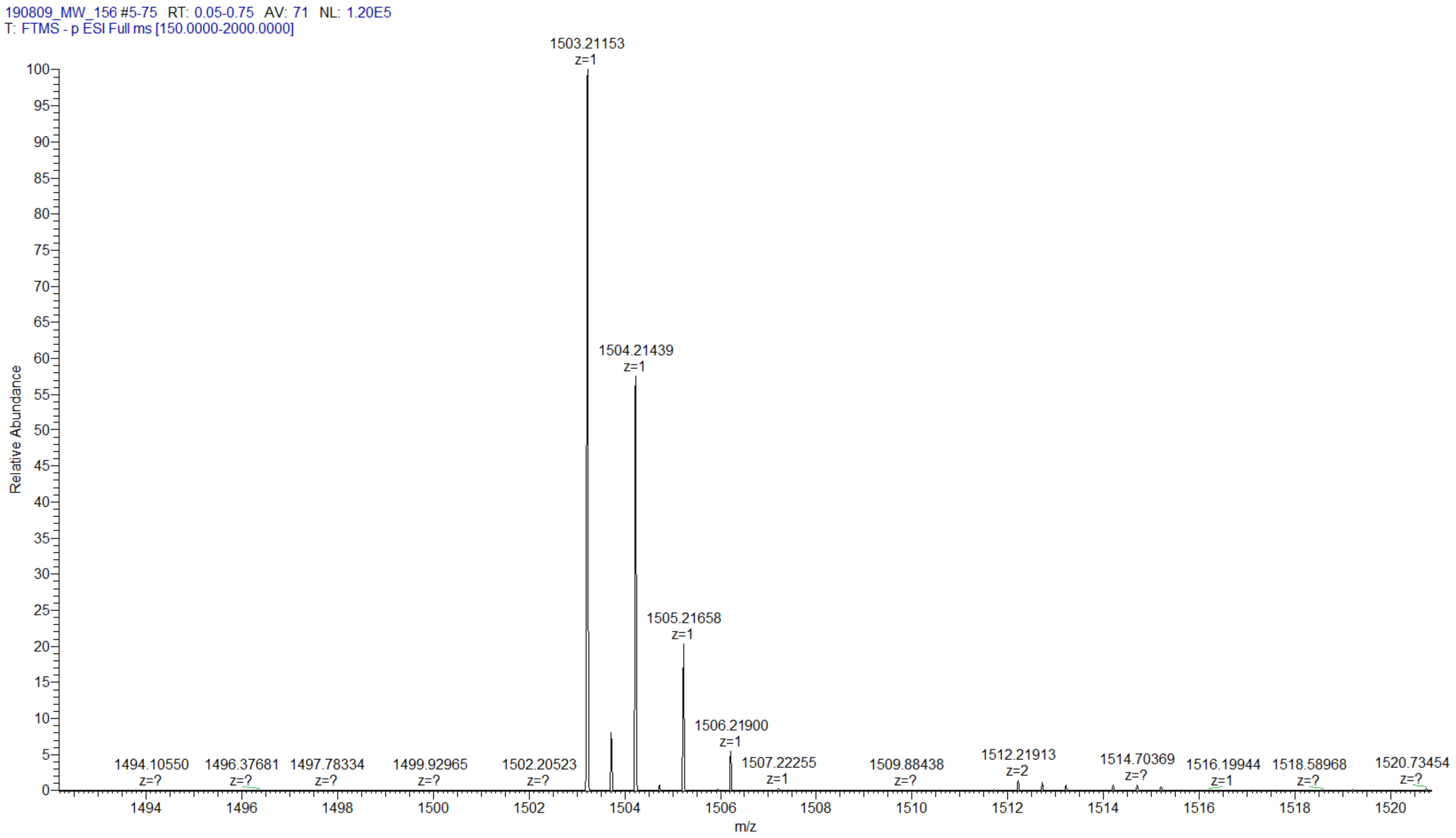 |

| m7Gppp(m6Am)pGmpG | |
|---|---|
| **Chemical structure** | 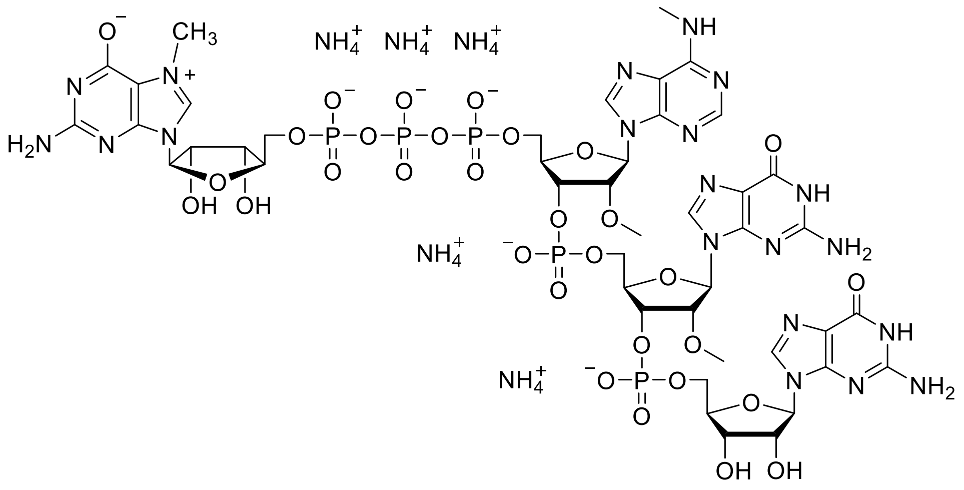 |
| **RP HPLC** | 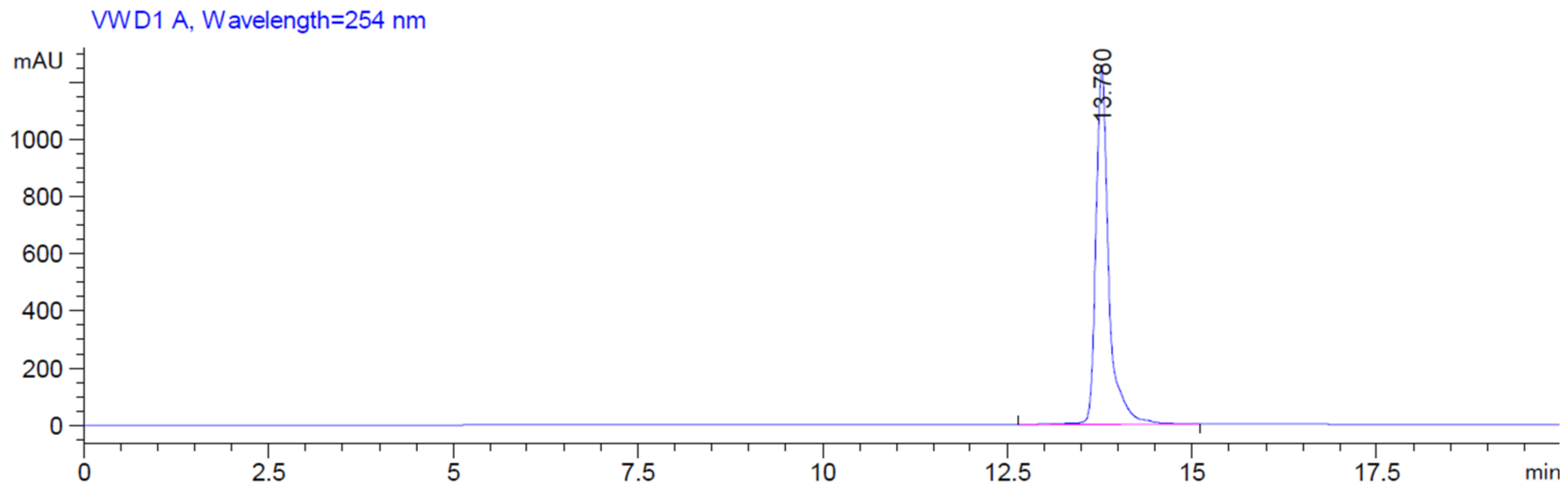  |

HRESI(-)MS Calc. [M-H]$^-$ $C_{44}H_{58}N_{20}O_{31}P_5^-$ 1517.2270 4)

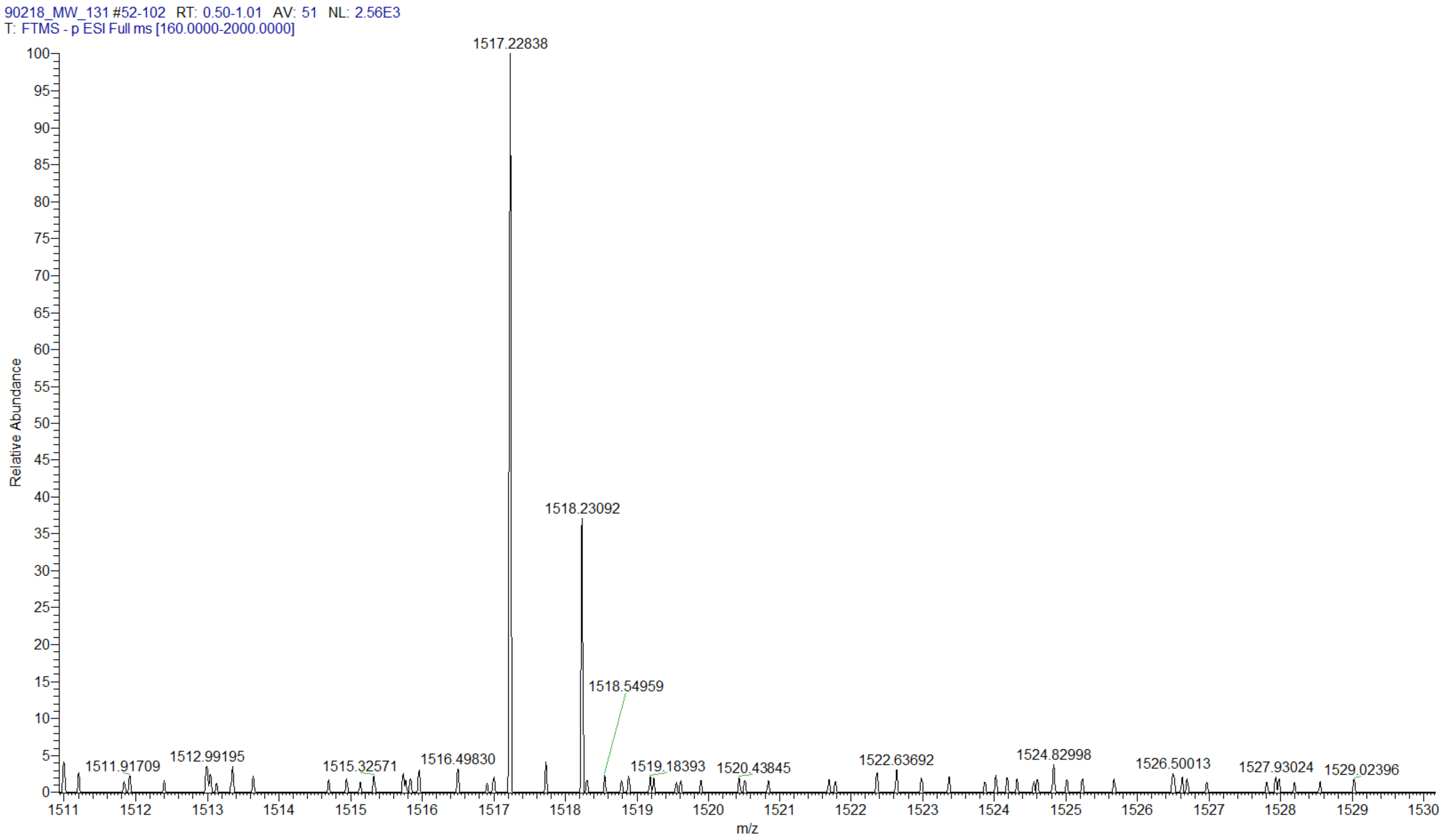

| m7GpppApGmpG | |
|---|---|
| Chemical structure | 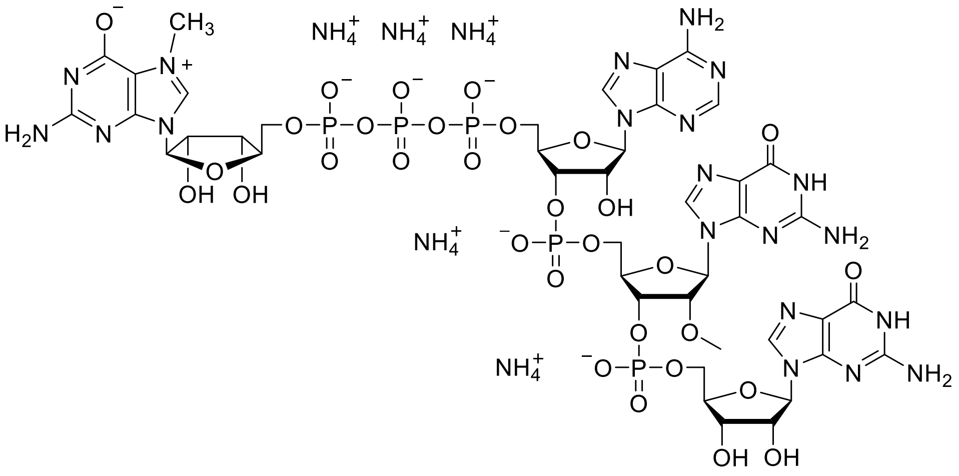 |
| RP HPLC | 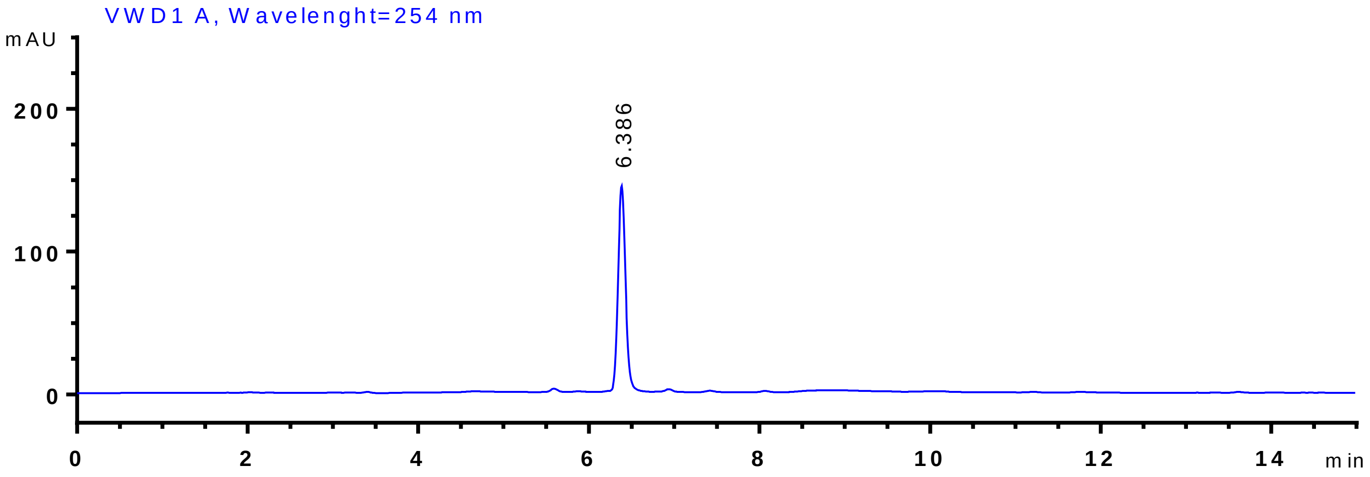 |

HR ESI (-) MS Calc. [M-H]$^-$ $C_{42}H_{54}N_{20}O_{31}P_5^-$ 1489.19574)

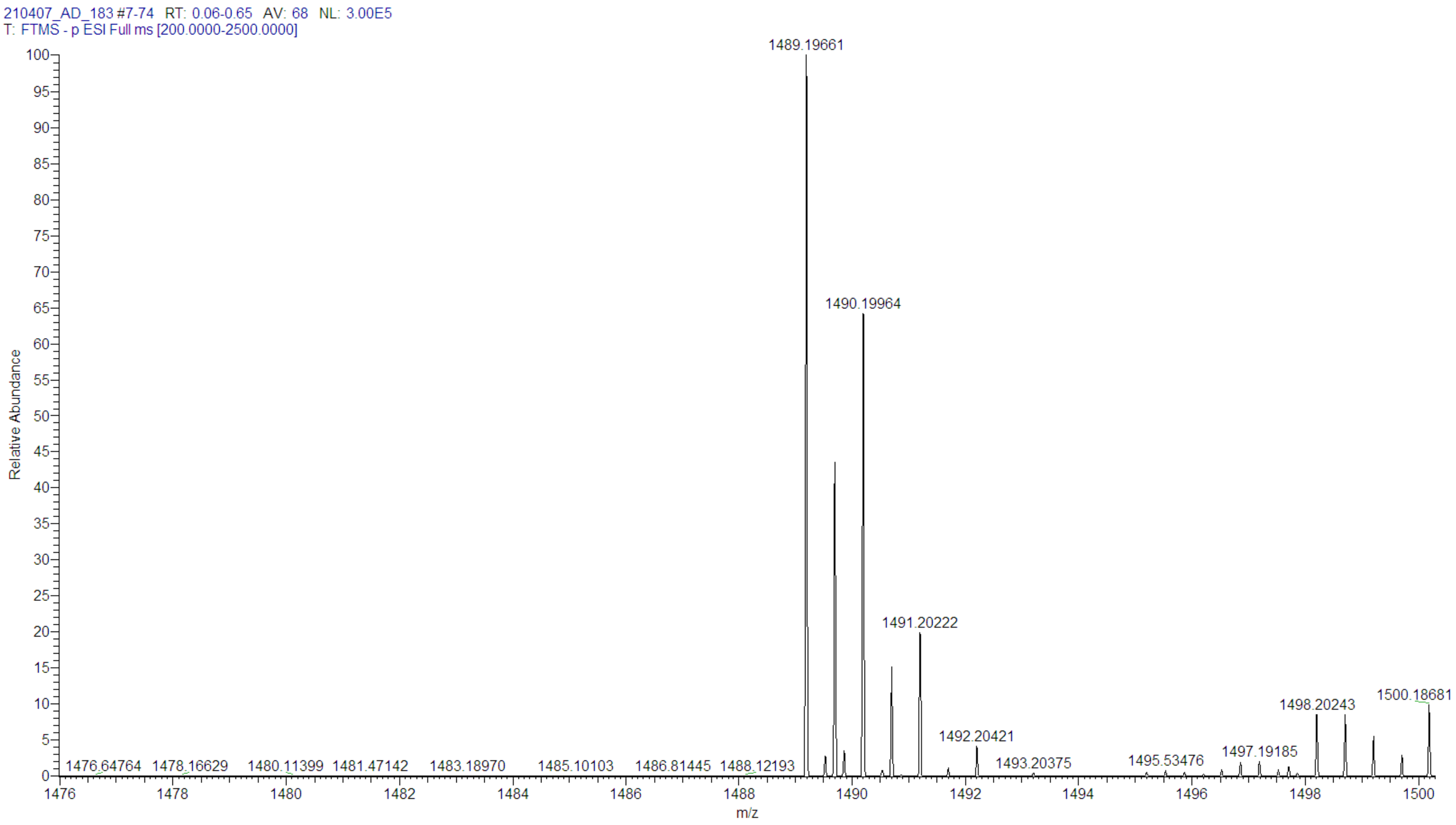

| m7Gppp(m6A)pGmpG | |
|---|---|
| **Chemical structure** | 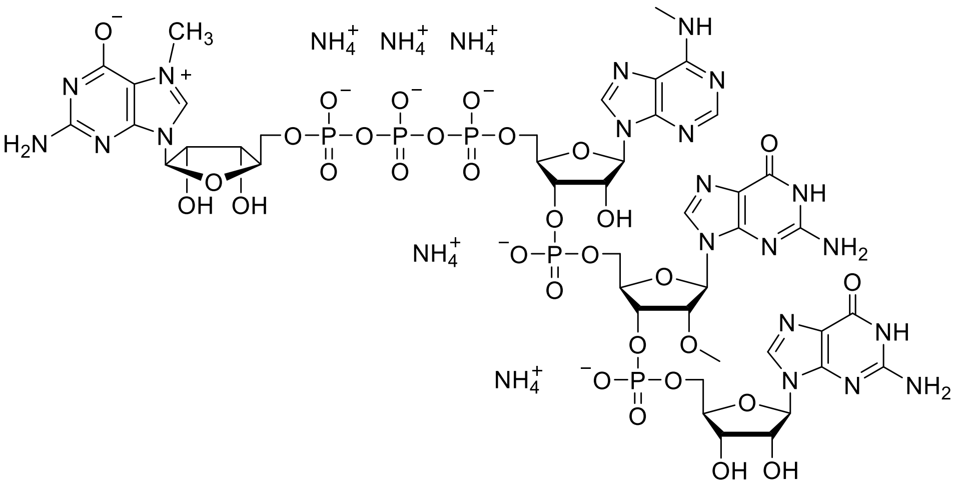 |
| **RP HPLC** | 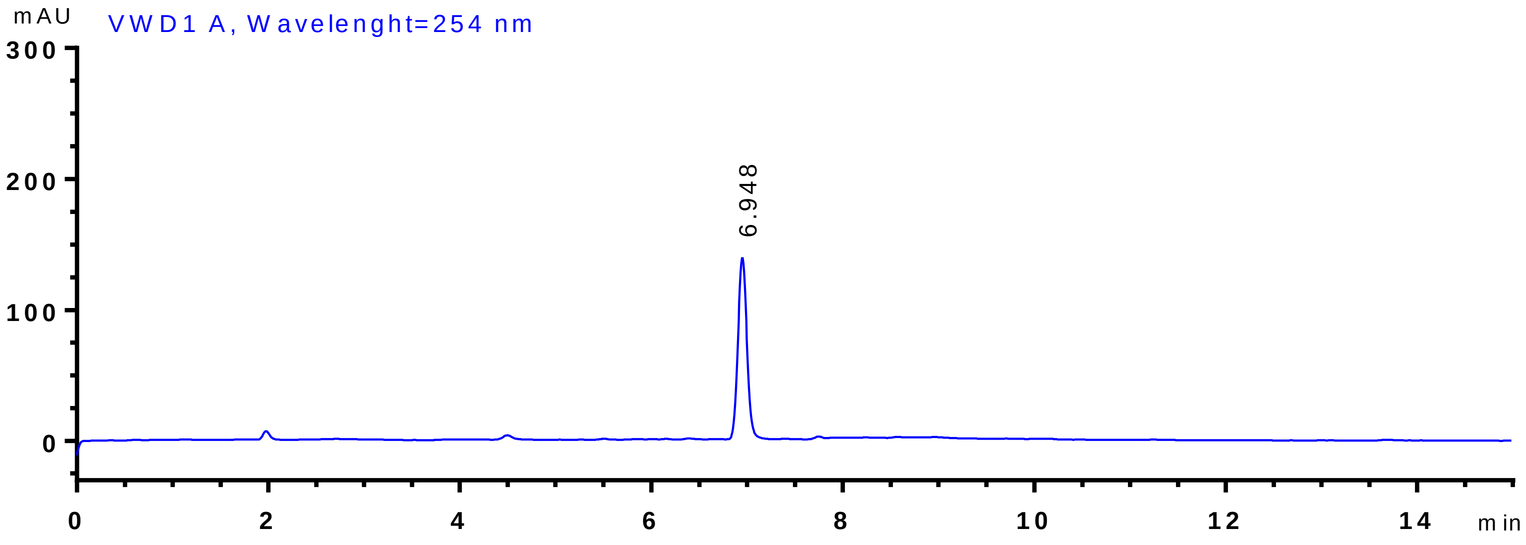 |

HR ESI (-) MS

Calc. [M-H]$^-$ $C_{43}H_{56}N_{20}O_{31}P_5^-$ 1503.21139)

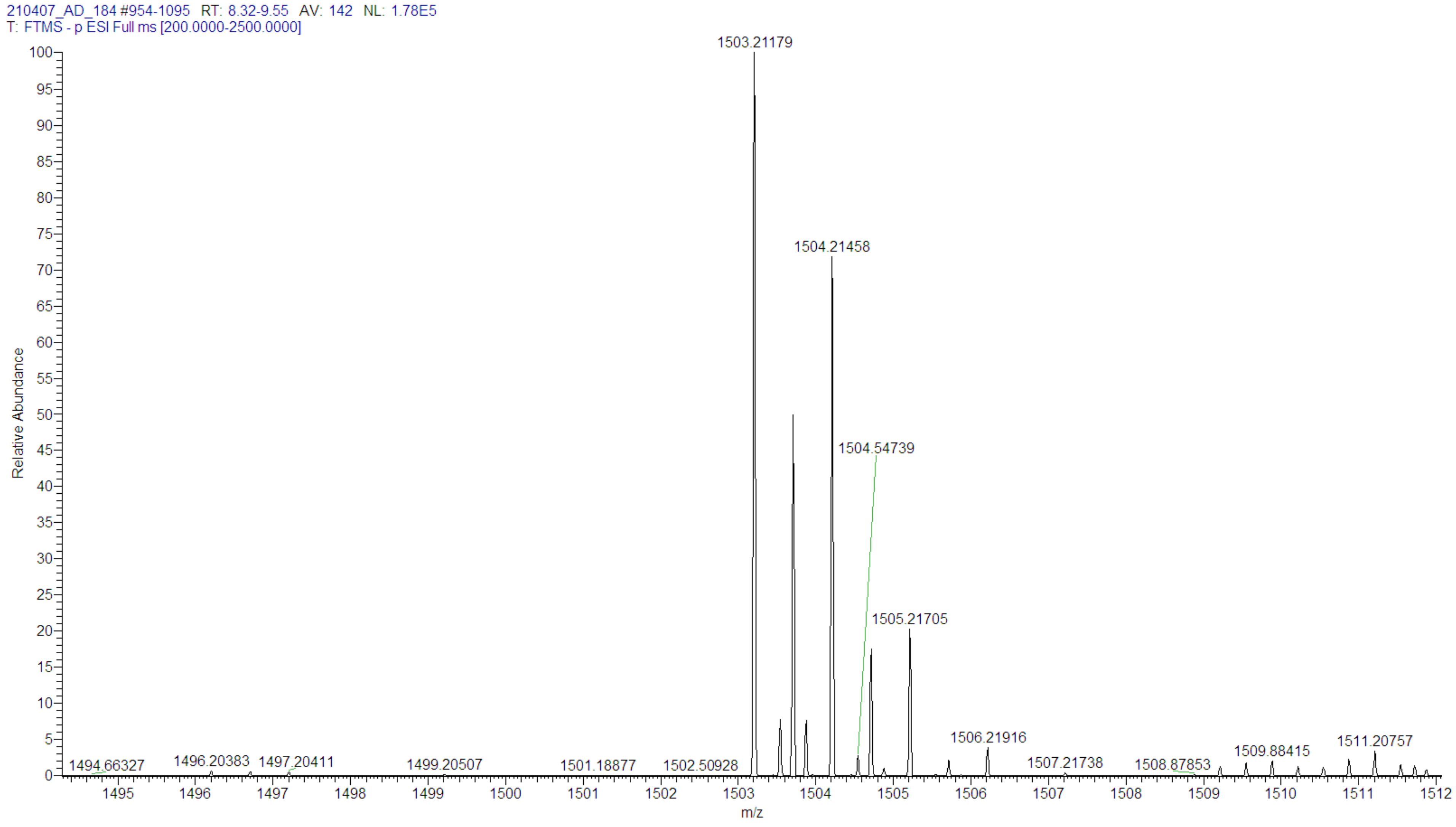

**Table S2.** List of oligonucleotides used in the study

| Oligonucleotide | Sequence | Purpose |
|---|---|---|
| IVTshF | CAGTAATACGACTCACTATAGGGGAAGCGGGCATGCGGCCAGCCATAGCCGATCA (10) | template for IVT of short RNA |
| IVTshR | TGATCGGCTATGGCTGGCCGCATGCCCGCTTCCCCTATAGTGAGTCGTATTACTG (10) | template for IVT of short RNA |
| DNAzyme | TGATCGGCTAGGCTAGCTACAACGAGGCTGGCCGC (10) | short RNA trimming |
| IFIT1_F | GATCAGCCATATTTCATTTTGAATC | qPCR |
| IFIT1_R | GAAAATTCTCTTCAGCTTTTCTGTG | qPCR |
| IFIT2_F | AAGAGGAAGATTTCTGAAGAGTGC | qPCR |
| IFIT2_R | TCTCCAAGGAATTCTTATTGTTCTC | qPCR |
| IFIT3_F | GAAGGAACTGGGCCGCCTGCTAAG | qPCR |
| IFIT3_R | GCCCTGGCCCATTTCCTCACTACC | qPCR |
| RIG-I_F | ATGTGCTCCTACAGGTTGTGG | qPCR |
| RIG-I_R | ACACTGGGATCTGATTCGCAA | qPCR |
| MDA5_F | GAGTCAAAGCCCACCATCTGA | qPCR |
| MDA5_R | CAGACCTTCTTCTGCCACTGT | qPCR |
| PKR_F | CAGACCCCTTGTCCAGACTTC | qPCR |
| PKR_R | CGGGAAGGGTAAGATGGCTTT | qPCR |
| OAS1_F | TCCTCAGTCCTCTCACCACTT | qPCR |
| OAS1_R | GAGCCTGGACCTCAAACTTCA | qPCR |
| GAPDH_F | ACCCACTCCTCCACCTTTGAC | qPCR |
| GAPDH_R | TGTTGCTGTAGCCAAATTCGTT | qPCR |
| hDXOfor | actggtggacagcaaatgggtcgc**ggatccATGGATCCCAGGGGGACCAAG** | cloning of full-length hDXO wt and mut into pET28-N-6xHis-TEV |
| hDXOrev | tcagtggtggtggtggtggtg**ctcgag*TTA*TTTGGGAGAGGGAGTCTTGG** | cloning of full-length hDXO wt and mut into pET28-N-6xHis-TEV |
| hDXOmutF | **AGCCACCCTCTGCTCTTCTCAGGGGCGGTAGCATGCACAGACCCCCAAGC** | cloning of full-length hDXO mut into pET28-N-6xHis-TEV |
| hDXOmutR | **GCTTGGGGGTCTGTGCATGCTACCGCCCCTGAGAAGAGCAGAGGGTGGCT** | cloning of full-length hDXO mut into pET28-N-6xHis-TEV |
| hDCP2F | GACGCGGCCGCCCCTTCACCTCACTTCATCAAAGAATTCTGCTG | cloning of 1-350 aa hDCP2 mut into pET28-N-6xHis-TEV |
| hDCP2R | GACGGATCCCTTATGGAGACCAAACGGGTGGAG | cloning of 1-350 aa hDCP2 mut into pET28-N-6xHis-TEV |

**Table S3** List of plasmids used in the study

| Plasmid | Genotype | Source |
|---|---|---|
| phDXOwt | [pET28M N-6xHis-TEV] hDXO wt | this work |
| phDXOmut | [pET28M N-6xHis-TEV] hDXO mut (E234A D236A) | this work |
| phDCP2wt | [pET28M N-6xHis-SUMOTag] hDCP2$^{1-350}$wt | gift from Prof. Megerditch Kiledjian |
| phDCP2mut | [pET28M N-6xHis-SUMOTag] hDCP2$^{1-350}$mut (E147Q E148Q) | this work |
